# Evaluating the impact of two different diets on the protein profile expression of the brain, liver, and intestine of barramundi

**DOI:** 10.1101/2025.11.03.686387

**Authors:** Mohadeseh Montazeri Shatouri, Igor Pirozzi, Pinar Demir Soker, Zeshan Ali, Ardeshir Amirkhani, Paul A Haynes

## Abstract

Commercial feed formulations are increasingly being evaluated for their nutritional impacts on aquaculture species, yet the molecular consequences of commonly used commercial diets remain underexplored. This study investigated the effects of two commercial diets, diet A (higher land animal protein) and diet B (higher fish meal content), on protein expression in the brain, liver, and intestine of barramundi (*Lates calcarifer*). A 12-week feeding trial was conducted with controlled water quality, and proteomic profiling was performed using data-independent acquisition. Differential analysis revealed consistent changes between diets across all tissues, with a higher percentage of differentially abundant proteins observed in between-diet comparisons (12.99% in brain, 12.73% in liver, and 16.59% in intestine) than within-diet controls (<8%), confirming a measurable dietary effect size. In total, 3,901 proteins in the brain, 3,660 in the liver, and 5,025 in the intestine were quantified. Functional enrichment highlighted upregulation of ferroptosis pathways, downregulation of apelin signaling in the brain, and increased digestive proteases in the liver. ICP-MS confirmed elevated iron concentrations in the brain, liver and intestine of fish fed on diet B. These findings demonstrate that molecular pathways linked to iron metabolism, digestion, and growth regulation are very sensitive to dietary composition, highlighting how proteomics can help identify subtle impacts of compositional differences in aquaculture feeding. Despite the absence of significant differences in physiological parameters, the proteomic alterations observed across tissues likely represent adaptive metabolic adjustments of each organ to the varying nutrient availability between diets.

## 1. Introduction

Barramundi (*Lates calcarifer*) are native to the Indo-Pacific region. They are found as far north as Taiwan, as far south as Fraser Island on the Australian east coast, eastwards as far as southeastern Papua New Guinea and westwards as far as the Persian Gulf. In the wild, the fish are a diadromous species, returning to estuarine or marine water to breed [1]. They are now also an important aquaculture species in the Indo-Pacific region, produced in both fresh and saltwater. The species is grown to a harvest size of between 400 and 4000 g, depending on the production system and market [2].

To sustain growth, barramundi, like all animals, require the dietary intake of certain nutrients. The dietary concentration required of these nutrients is largely driven by the energetic content of the food eaten by the fish. Several studies have been undertaken to examine the requirements for protein in barramundi diets. Most of these studies suggest a relatively high protein requirement, consistent with the carnivorous/piscivorous nature of the fish [3]. Despite these insights, research on the individual amino acid requirements remains limited; to date, only four of the ten essential amino acids have been thoroughly investigated, with existing estimates focusing on methionine/total sulfur amino acids, lysine, arginine, and tryptophan [4]. In addition, lipids - being the most energy-dense nutrient - have received considerable attention, as they provide nearly double the energy of protein and significantly more than carbohydrates [2]. Proteomics has been used to investigate food intake and appetite in other monogastric animals [5]. Omic-based studies on barramundi have so far been limited to phenotype characterization [6], speciation and adaptation [7], sensory and volatile characteristics [8], and hepatic transcriptomes [9]. In a previous study, the use of animal protein sources as a replacement for fishmeal in fish diets had a positive impact on the feed conversion ratio, variable growth rate, final weight, and survival rate of different types of fish species of different size groups [10]. However, little is known about how changes in physiological processes associated with food intake in fish are reflected at the proteome level. Advances in proteomic technologies now offer a powerful means to explore the molecular mechanisms underpinning these nutritional requirements. High-throughput proteomics allows for the simultaneous profiling of thousands of proteins, providing insights into the regulatory networks and metabolic pathways that are modulated by dietary composition [11]. This type of molecular-level investigation is critical because it can reveal early biomarkers of nutritional stress or suboptimal dietary formulations well before changes in growth performance or health become apparent.

In this study, we conducted a 12-week feeding trial using two different commercial diets (designated diet A and diet B) provided by Skretting Australia. These diets differed in ingredient composition but shared the same crude protein and crude fat levels (crude protein/crude fat: 45/20). Diet A contained less fish meal and more land-animal protein sources than diet B and was also characterized by lower levels of methionine as an essential amino acid. These compositional differences provided an opportunity to investigate how subtle shifts in dietary protein quality and ingredient origin affect tissue-level protein expression and nutrient regulation.

Our aim was to compare the proteomic profiles of key tissues including brain, liver, and intestine to gain a more comprehensive understanding of how diet influences metabolic regulation and cellular function in barramundi. These three tissues were selected for their pivotal roles in maintaining overall physiological balance [12]. Liver plays a central role in the digestion, transport, metabolism, and storage of nutrients, as well as detoxification, immunity and health [13]. Brain integrates neural and hormonal signals to regulate feeding behaviour [14], and intestine is the primary site for nutrient digestion, motility, secretion, and absorption [12].

We hypothesized that the nutrient composition differences between diet A and diet B would lead to distinct alterations in proteomic profiles of different tissues. The insights gained from this study will improve our understanding of the dietary requirements of farm-raised barramundi.

## 2. Materials and Methods

### 2.1 Feeding, handling and treatment of barramundi

The feeding trial was conducted over a period of 12 weeks, using two different commercial diets manufactured by Skretting Australia (Cambridge, TAS, Australia). Relative to diet B, diet A contained 60% less fish meal (wild catch sardine), 100% more rendered poultry meal, 33% less hydrolysed feather meal, 1.7 times more high-quality blood meal, 39% less phospholipids, 22% less methionine, and 1.3 times more iron. Levels of other components were essentially the same between the two diets, including crude protein, crude fat, lupin, faba, total amino acids, total fatty acids, starch, fibre, vitamins, zinc, copper, and antioxidants.

The experimental design involved stocking 25 barramundi, each weighing approximately 500 g, into two 1000 L aquaculture tanks per diet treatment. The experimental tanks were integrated within a 30,000 L recirculating aquaculture system which provided mechanical and biological filtration and allowed control of water flow. Feeding was administered twice daily on weekdays, at 9 am and 2 pm, and once at 9 am on weekends.

Fish were anaesthetized (AQUI-S), and tissue samples from the brain, liver, and intestine of 8 fish fed on diet A and 8 fish fed on diet B were collected after the measurement of growth rate parameters. The tissue samples were randomly assigned into two groups of four from each diet, to facilitate subsequent analysis using four replicates each for both within – diet and between – diet quantitative comparisons. The samples were stored on dry ice and transferred to -80°C for storage until further processing.

### 2.2 Water quality

Water quality variables were recorded daily (at 8:30 am and 3:30 pm) using portable electronic instruments. The average water temperature during the experiment was approximately 28°C ± 1. Salinity ranged from 30% to 33%. Dissolved oxygen concentration remained above 7 mg L^−1^. Throughout the experiment oxygen saturation remained between 120% and 180%, and pH ranged from 6.9 to 8.4. Total ammonia nitrogen (TAN) concentration was measured regularly using a colorimetric test kit, with the average TAN concentration ≤ 0.8 mg L^−1^.

### 2.3 Calculated production indices

Performance indices including growth rate, food intake, and food conversion ratios were calculated using the formulae below[15]: Specific growth rate (% d^−1^) = (Ln (final weight) – Ln (initial weight)) / days x 100 Relative food intake (g kgBW^−1^ d^−1^) = daily food intake per fish (g) / (geometric mean body weight of fish (kg)) Food conversion ratio (FCR) = food intake per tank (g) / wet weight gain per tank (g)

### 2.4 Protein extraction from brain, liver and intestine

The tissues collected from fish grown on both diets, stored at -80°C, were ground on dry ice. Approximately 20 mg aliquots from brain, liver and intestine were transferred to 2 mL Eppendorf tubes kept on dry ice (8 biological replicates per tissue). A volume of 200 µL of RIPA buffer (50 mM Tris, pH 7.5, 150 mM NaCl, 1% NP40, 1 mM EDTA, 0.1% SDS containing 1% v/v protease inhibitor cocktail) was added to the tube, and the tissue was homogenized using a TissueLyser (IKA, T 10 basic ULTRA-TURRAX) for 30 seconds. Subsequently, the sample was sonicated twice using a probe sonicator (40 Hz × 2 pulses, each time 15 seconds, with 30-second intervals on ice). The samples were then centrifuged for 15 minutes at 13,000 rpm at 4°C to remove debris. The supernatant was transferred to a clean tube. Proteins were reduced using 20 mM Dithiothreitol (DTT) for 45 minutes. The reduced protein extract was then alkylated with 50 mM Iodoacetamide for 30 minutes in the dark. Alkylation was stopped by adding 40 mM DTT.

Protein extracts were precipitated using methanol/chloroform. First, 800 µL of methanol (4 times the volume of the lysis buffer), 200 µL of chloroform (equal to the volume of the lysis buffer), and 600 µL of water (3 times the volume of the lysis buffer) were added to the supernatant. The mixture was vortexed and centrifuged for 1.5 minutes at 13,000 rpm. The upper phase was removed, and 800 µL of methanol was added to the tube, followed by centrifugation for 5 minutes at 13,000 rpm. The supernatant was removed, and the pellet was air-dried in a fume hood. The protein pellet was solubilized in 8 M urea.

### 2.5 Protein quantification and digestion

Bicinchoninic acid (BCA) assay kit (Thermo Scientific, San Jose, CA, USA) was used to measure the concentration of protein in the sample extract. From each sample, 100 µg of protein was transferred to a 1.5 mL tube. Trypsin (1 µg) was added to each sample, and digestion occurred overnight at 37°C. Digestion was halted by adding TFA to adjust the pH to 3. Samples were desalted using SDB-RPS (3M Empore) 200 µL stage tips [16]. The membranes were first conditioned with 100 µL of acetonitrile. Subsequently, 100 µL of 1% trifluoroacetic acid (TFA)/30% methanol and 100 µL of 0.2% TFA were sequentially loaded onto the membrane. Samples were loaded onto the conditioned tips and centrifuged. The columns were then washed with 0.2% TFA to remove unbound material. Tryptic peptides were eluted with 200 µL of 80% ACN, 5% ammonium hydroxide into a new tube. The eluted peptides were dried using a vacuum centrifuge and resuspended in 35 µL of 0.1% formic acid. A micro-BCA assay was performed to determine the peptide concentration.

### 2.6 Peptide fractionation for spectral library generation

A pooled sample containing 200 µg of protein was prepared for each tissue separately by combining 12.5 µg each from all 16 replicates, and fractionated using high-pH RP-HPLC, with 17 fractions collected. The fractionation was performed using an Agilent 1260 HPLC system equipped with a quaternary pump (G1311B), an autosampler (G1329B) with a 900 µL injection loop and syringe, a column compartment (G1316A), a fraction collector (G1364C), and a thermostat for the autosampler and fraction collector (G1330B). The analysis was conducted using ChemStation chromatography software. A Zorbax 300Extend- C18 column (2.1 × 150 mm, 3.5 µm, 300 Å) was used, along with a 96-well, 2 mL deep well-rounded bottom polypropylene collection plate. The gradient elution method consisted of an 85-minute gradient, starting with 97% mobile phase A (milli-Q water with 6 mM ammonium hydroxide) and 3% mobile phase B (6 mM ammonium hydroxide in 90:10 ACN: MilliQ water) for 10 minutes, followed by 55 minutes of 70% mobile phase A and 30% mobile phase B, 10 minutes of 30% mobile phase A and 70% mobile phase B, 5 minutes of 10% mobile phase A and 90% mobile phase B, and finally 5 minutes of 97% mobile phase A and 3% mobile phase B, all at a flow rate of 0.3 mL/min and a column temperature of 25°C. UV detection was performed at wavelengths of 210 nm and 280 nm.

### 2.7 Nanoflow Liquid Chromatography–Tandem Mass Spectrometry (nanoLC-MS/MS) analysis of pooled samples for library generation

Fractions prepared by HpH reverse phase fractionation for library generation were analyzed on a Q-Exactive HF-X mass spectrometer (Thermo Fisher Scientific) interfaced with an UltiMate 3000 UHPLC (Thermo Fisher Scientific) column using Data-Dependent Acquisition (DDA) mode with a full scan followed by fragmentation of the top 10 most abundant ions. A 90-minute gradient was employed, beginning with 98% mobile phase A and 2% mobile phase B for 10 minutes, followed by 60 minutes of 65% mobile phase A and 35% mobile phase B, 8 minutes of 5% mobile phase A and 95% mobile phase B, 5 minutes of 5% mobile phase A and 95% mobile phase B, and 4 minutes of 98% mobile phase A and 2% mobile phase B. The flow rate was set at 300 nL/min, using a 75 µm × 30 cm C18 column (Dr. Maisch ReproSil-Pur 120 C18-AQ) maintained at 45°C. The analysis was conducted in positive ion mode with automatic gain control (AGC) set at 3×106 for ion accumulation, a maximum trapping time of 50 milliseconds, a scan range of 350 to 1650 m/z, a full scan resolution of 60,000, MS2 resolution of 15,000, with HCD fragmentation of the top 10 ions, a normalized collision energy of 27.5 and dynamic exclusion enabled for 15 seconds.

### 2.8 nanoLC-MS/MS Data-Independent Acquisition proteomic analysis of tissue samples

Tryptic peptides prepared from tissue samples taken from fish fed on different diets were analysed on a Q-Exactive HF-X mass spectrometer (Thermo Fisher Scientific) interfaced with an UltiMate 3000 UHPLC (Thermo Fisher Scientific) using Data-Independent Acquisition (DIA) mode, employing the same column and gradient used for the library generation fractions. Twenty-one dynamic windows with precursor ions within isolation windows ranging from 22.0 to 589.0 m/z were subjected to fragmentation by HCD using a normalized collision energy of 27.5. MS2 scan resolution was set at 30k, and an AGC target value of 2×105 was used.

### 2.9 Data analysis

Raw files obtained from 17 fractions were analysed using MSFragger version 21.1 to produce a spectral library using a barramundi fasta file of 58,000 protein sequences entries downloaded in February 2024 from UniProt. The library generated by MSFragger from each tissue was used for DIA-NN (1.8.1) search for 16 samples of different two diets. The report.pg_matrix file generated by DIA-NN was used for statistical analysis by fragpipe analyst [17] using DIA type, Max-LFQ Intensity, with a p-value cutoff of p< 0.05, fold change of ≥ 1.3 and ≤ 0.76, variance stabilizing normalization and MLA imputation. Partial least squares discriminant analysis (PLS-DA) was performed using MetaboAnalyst 6.0 [18], and volcano plots of the three tissue samples were generated using Microsoft excel. Functional analysis of identified proteins including gene ontology (GO) and KEGG pathways were performed using the String database [19].

### 2.10 Parallel Reaction Monitoring Analysis

Parallel Reaction Monitoring (PRM) analysis was used for the validation of the label-free shotgun proteomics results, to measure quantitative changes in specific proteins in Barramundi’s tissue fed on diet A and diet B. Skyline-daily was used to create an inclusion list of unique peptides, for each selected protein, with their mass to charge ratio [20]. The obtained PRM data were imported into Skyline and subject to quality control analysis. The sum of areas of all transition peaks for each peptide were collected and log2-transformed for four biological replicates of diet A and four biological replicates of diet B, and Student t-test analysis was used to compare protein abundance between two diets.

### 2.11 Total Iron analysis

Approximately 20 mg of liver, and 50 mg of brain and intestine tissues from each biological replicate (n= 4 per tissue) was transferred to a microwave digestion tube (Multiwave 3000 Digestion Microwave, Anton Paar Australia pty.Ltd). A total of 5 mL of 69% nitric acid and 1mL of 30% hydrogen peroxide was added to each. The microwave digestion settings were as follows: temperature, 180 °C; pressure, 500 psi; power, 1000−1800 w; ramp time, 20 min; and hold time, 20 min. After complete digestion and cooling for 10 min, the resultant solutions were transferred to 50 mL falcon tubes, and the volume of the replicates was adjusted to 30 mL with Milli-Q water. Samples were diluted 9000x times and analyzed by inductively coupled plasma-mass spectrometry (ICP−MS) with a limit of detection of parts per billion (ppb) [21].

## 3. Results & Discussion

### 3.1. Growth performance

At the end of the 12-week feeding trial, the average body weight of barramundi reached 683 g for fish fed with Diet A and 684 g for those fed with Diet B, showing no significant difference between treatments. Weights were recorded at both the beginning and end of the feeding period to monitor growth progression. At the start of the trial, fish averaged approximately 500 g in weight.

Feed intake averaged 864 g for Diet A and was modestly lower at 828 g for Diet B; however, this variation was not statistically significant. In contrast, the FCR was significantly lower in fish fed Diet B (1.210) compared to those fed Diet A (1.265), indicating slightly better feed efficiency in the Diet B group.

### 3.2. Numerical summary of proteins identified in different tissues

Data independent analysis quantitative proteomics was used to analyse four replicates of three tissue samples of barramundi fed on two different commercial diets designated as diet A and B. The number of proteins identified and quantified in each replicate for diet A range from 2868 to 3342 for liver, 3270 to 3501 for brain and 4295 to 4482 for intestine, and for diet B range from 2941 to 3356 for brain, 2051 to 3069 for liver, and 4471 to 4545 for intestine, as shown in Table 1. Details of all these identified proteins are presented in supplementary data S1.

**Table 1.**
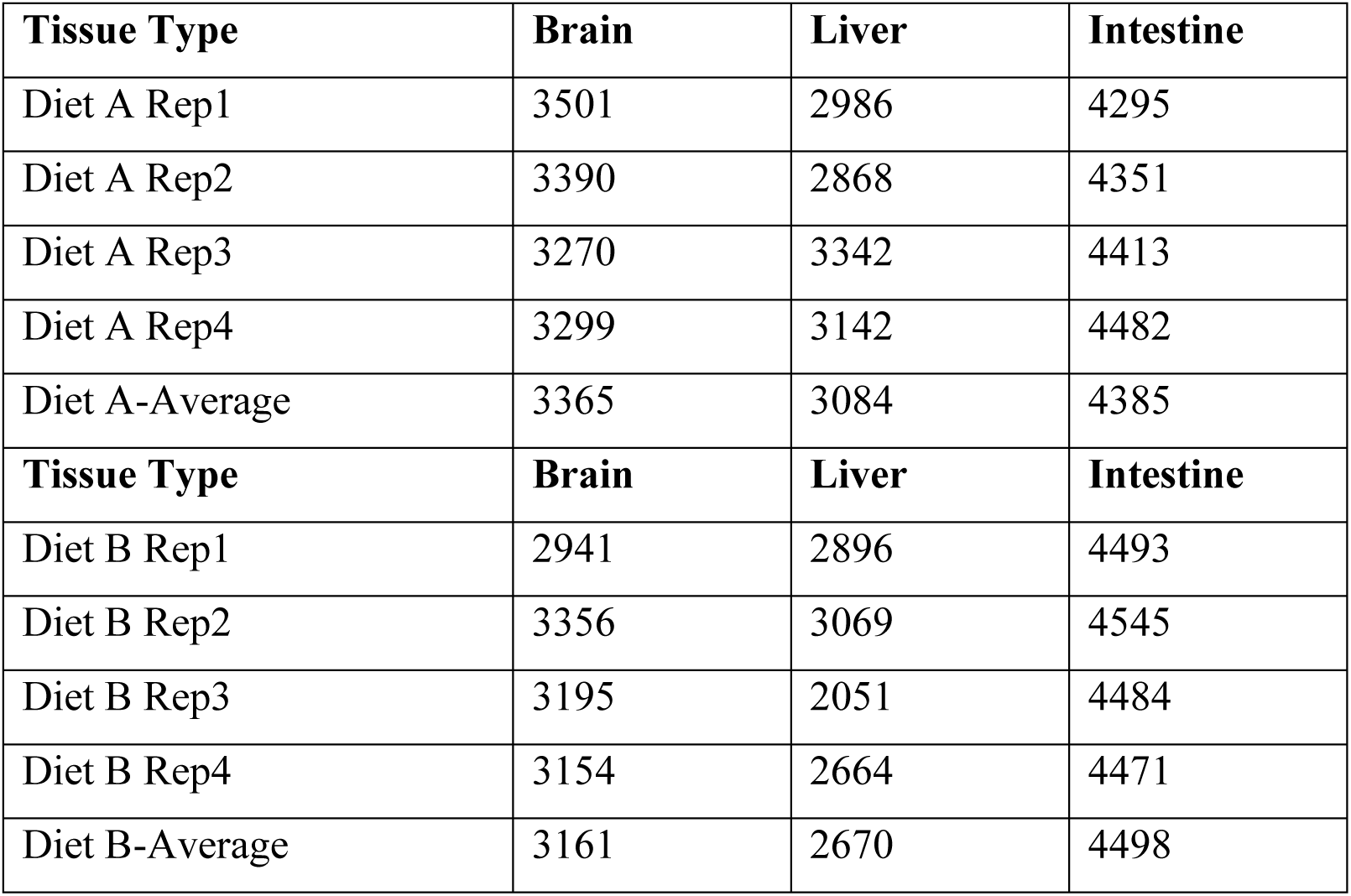
Summary of number of proteins identified in tissue replicates

### 3.3 Estimating effect size of within-diet vs between-diet comparisons

In biological experiments, quantitative differences at the molecular level can stem from three main sources: biological variability, reflecting the natural diversity between organisms, subjects, or tissue samples; technical variability, which arises from the way we handle and measure those samples, including factors like instrumentation, assay methods, and preparation steps; and induced biological change, which is what we are trying to measure in order to indicate the effect of what external factors are imposed on the biological system [22]. To determine how much the observed changes in protein expression were influenced by diet, we first performed within-diet comparisons as a control. This involved comparing results from 4 replicates of Diet A with results from a second set of four replicates of Diet A (A vs A) and performing a similar analysis with results from two sets of four replicates of Diet B (B vs B). These results were then compared to the variability observed between 4 replicates of Diet A and 4 replicates of Diet B (A vs B). Expressing the number of differentially abundant proteins as a percentage of the total proteins identified in each comparison allows us to generate an approximate measure of effect size. If dietary composition had no significant effect, we would expect similar levels of effect size in both within- and between-diet comparisons. However, a noticeably higher percentage of differentially expressed proteins between diets would suggest that the dietary treatments had a greater effect size and thus a measurable impact on the proteomic profile.

Our analysis clearly showed that protein expression differences were greater between the two diets than within the same diet, across all three tissues (see Tables 2, 3, 4). For example, in brain samples (Table 2), comparisons within the same diet showed only 6.01% (Diet A) and 2.41% (Diet B) of proteins were differentially abundant. But when comparing Diet A to Diet B, this jumped to 12.99%. A similar trend was seen in the liver in Table 3 (7.86% and 5.77% vs 12.73%) and intestine in Table 4 (2.20% and 5.21% vs 16.59%).

**Table 2.**
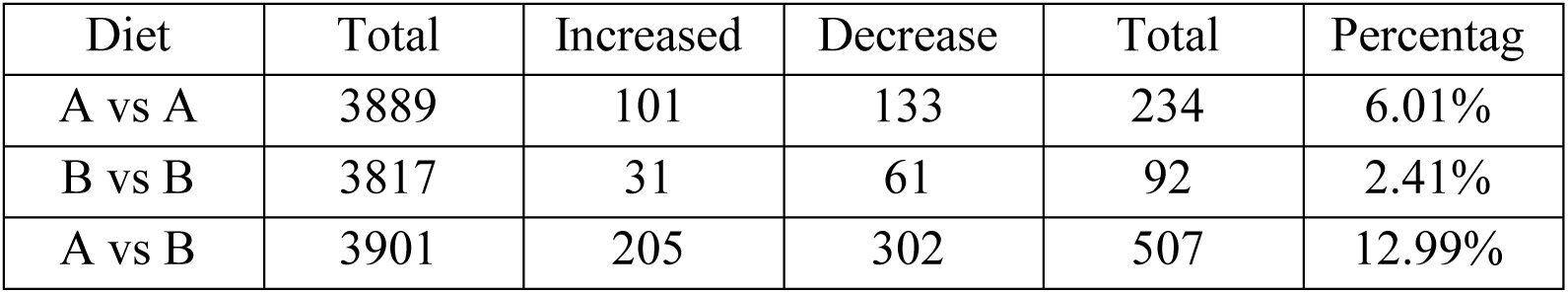
Comparison of brain samples within and between diets

| Diet | Total | Increased | Decrease | Total | Percentag |
| --- | --- | --- | --- | --- | --- |
| A vs A | 3889 | 101 | 133 | 234 | 6.01% |
| B vs B | 3817 | 31 | 61 | 92 | 2.41% |
| A vs B | 3901 | 205 | 302 | 507 | 12.99% |

**Table 3.**
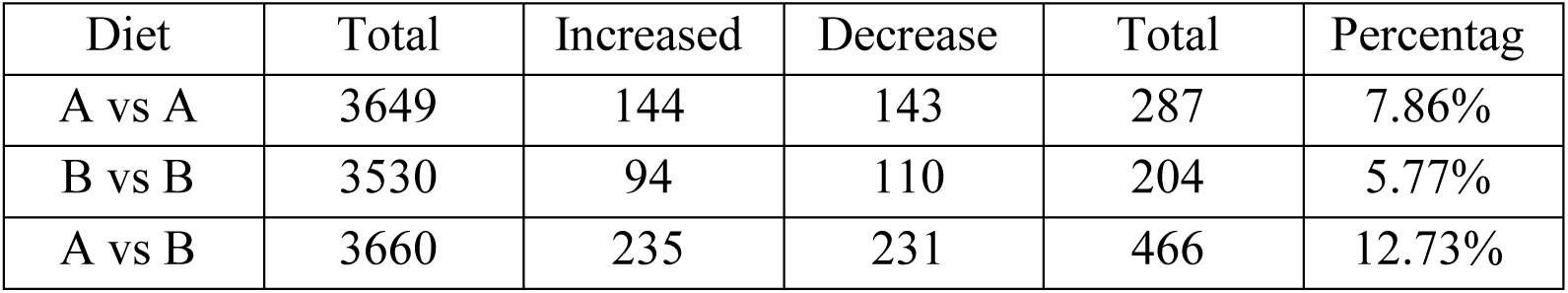
Comparison of liver samples within and between diets

| Diet | Total | Increased | Decrease | Total | Percentag |
| --- | --- | --- | --- | --- | --- |
| A vs A | 3649 | 144 | 143 | 287 | 7.86% |
| B vs B | 3530 | 94 | 110 | 204 | 5.77% |
| A vs B | 3660 | 235 | 231 | 466 | 12.73% |

**Table 4.** Comparison of intestine samples within and between diets

| Diet | Total | Increased | Decrease | Total | Percentag |
| --- | --- | --- | --- | --- | --- |
| A vs A | 5030 | 48 | 63 | 111 | 2.20% |
| B vs B | 5003 | 135 | 126 | 261 | 5.21% |
| A vs B | 5025 | 485 | 349 | 834 | 16.59% |

The consistent differences across all tissues between the two diets suggest that what the fish were fed had a measurable effect on the proteomic profile in each organ. Even though the diets were both commercially formulated and suitable for use in feeding trials, they still had a clear and consistent impact on the protein expression profiles in barramundi. By comparing replicates within the same diet, we were able to confirm that the differences seen between diets are not just due to biological or random variation. This supports the idea that the proteins identified as differentially abundant between the two diets are altered in response to dietary changes and could serve as useful index for evaluating the adaptive response of fish to the available nutrients.

### 3.4 Statistical analysis of protein expression in brain

We employed the Limma software package to perform statistical comparisons by fitting a linear model and applying empirical Bayes moderation to estimate variance. Protein abundances were log2-transformed, and group means were compared to calculating fold changes. This is equivalent to comparing geometric means on the raw scale but provides greater statistical robustness, particularly in the presence of missing values. Proteins detected in at least 3 out of 4 replicates per group with a p-value < 0.05 and a fold change of ≥ 1.3 and ≤ 0.76 were considered as differentially abundant proteins (DAPs).

Analysis of PLS-DA of the brain proteome data revealed evidence of the clustering between protein expression of the brain tissue (Figure 1). There is greater dispersion among diet B samples, suggesting higher inter-individual variability in response to diet B. The volcano plot provided in figure 2 gives a comprehensive overview of protein expression in the brain tissue. The clear separation of increased and decreased proteins in brain indicates that brain tissue exhibits a wide range of differential protein expression. This may reflect subtle but meaningful shifts in protein function that were not apparent in the physiological measurements, suggesting that proteomic analysis can reveal diet-induced responses even when overall diet composition and physiological outcomes appear similar. As shown in Table 2 a total of 302 proteins were decreased, and 205 proteins were increased in brain in fish fed on diet B. Full details of all these differentially abundant proteins are presented in Supplementary data S2.

**Figure 1.**
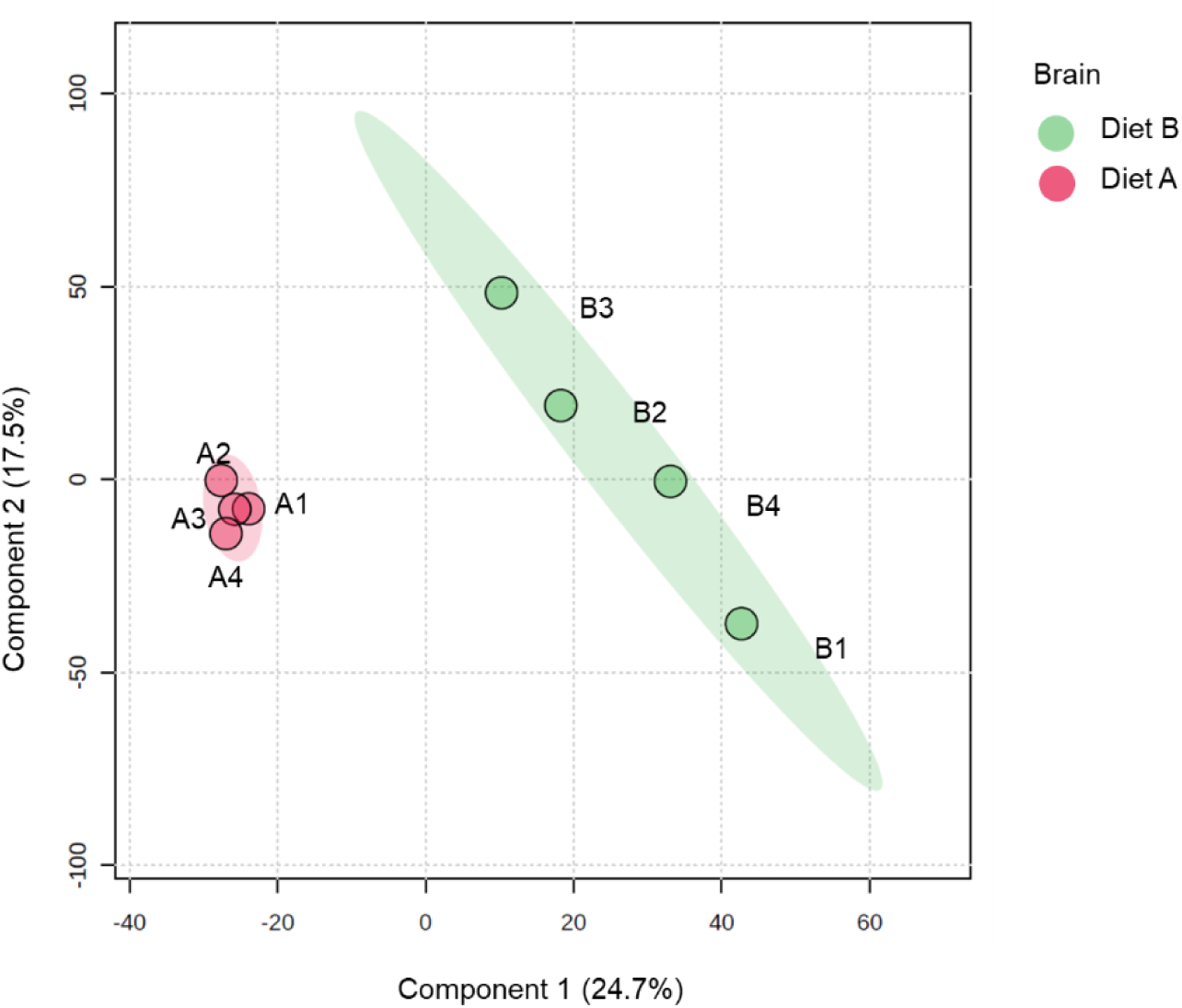
Partial least squares discriminant analysis (PLS-DA) of brain of barramundi fed on diet A and diet B displayed at 95% confidence intervals highlights the clustering of the sample based on their response to different diets (n= 4).

**Figure 2.**
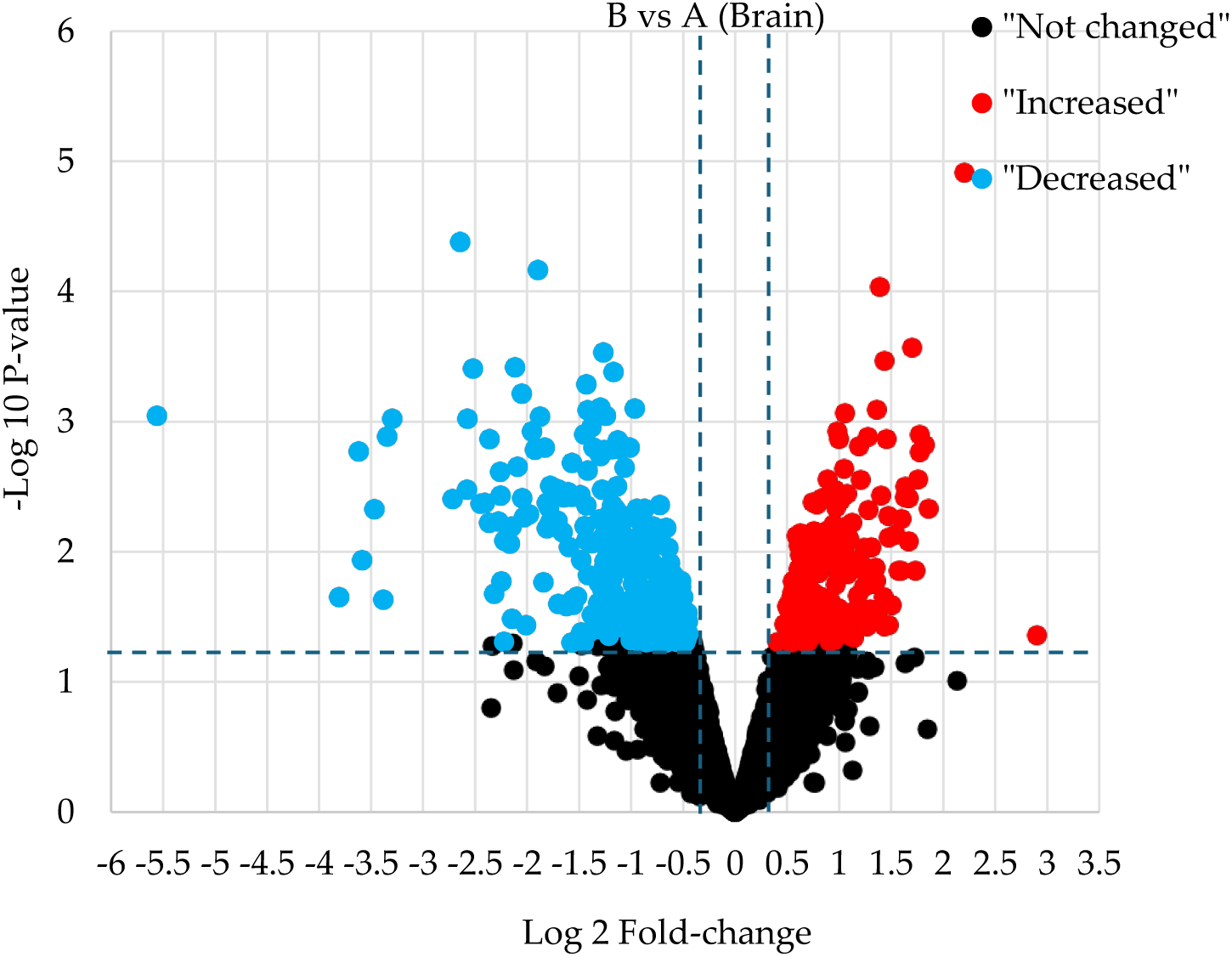
Volcano plot illustrates significantly differentially abundant proteins in brain. The −log10 (P value) is plotted against the log2 (fold change: diet B/diet A).

#### 3.4.1 Differentially abundant proteins in brain

The brain regulates nearly all biological functions and behaviours, including eating and digestion. Its response to food and metabolic status, along with its ability to integrate information, influences an individual’s nutritional and emotional state. This interaction shapes the balance between hunger and satiety, the experience of pleasure, and the development of goal-directed behaviours [23]. Table 5 presents the top ten DEPs in the brain (presented in detail in supplementary data S2) when comparing the different diets, sorted by fold change.

**Table 5.**
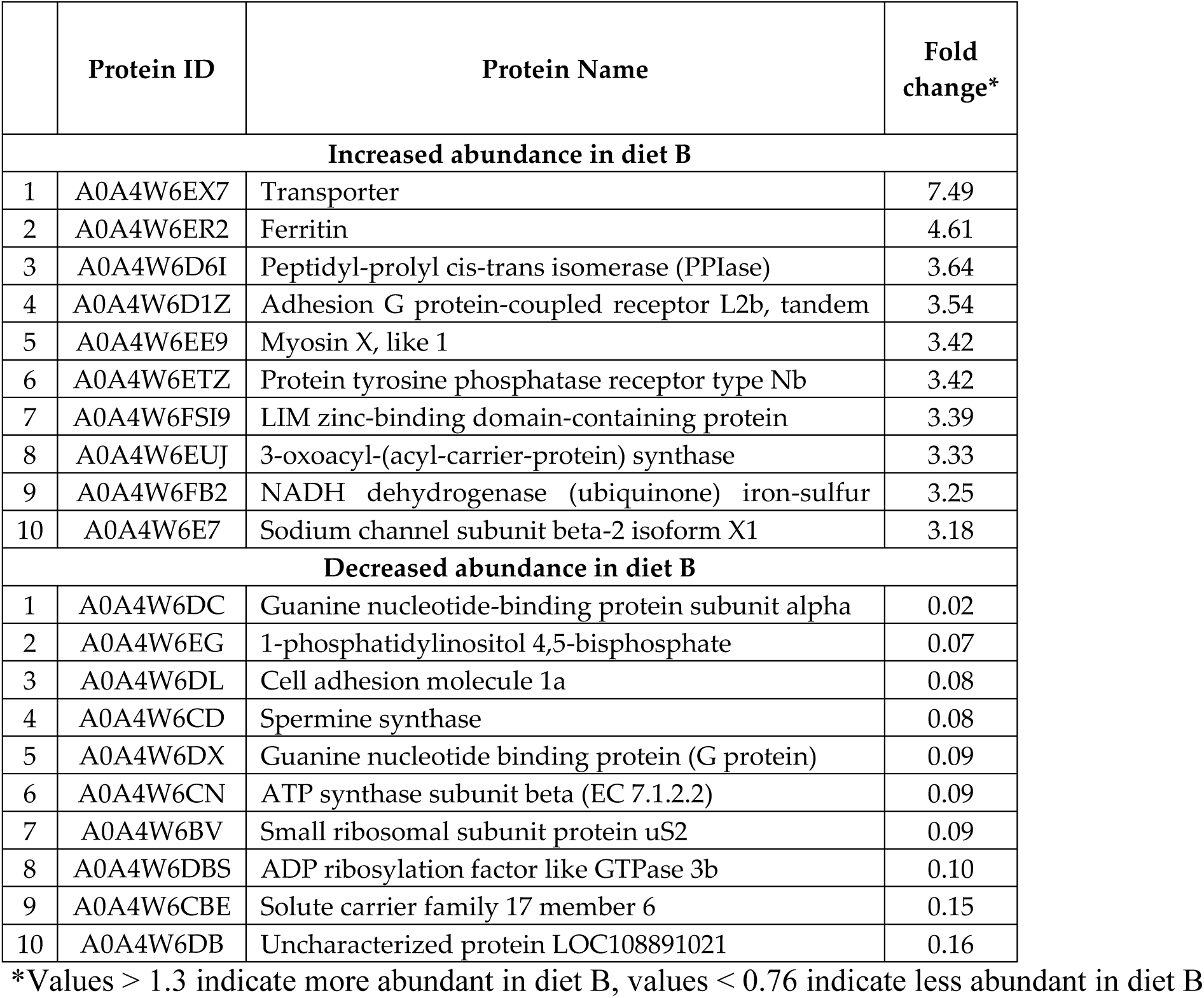
Top ten DAPs in brain tissue

|  | Protein ID | Protein Name | Fold change* |
| --- | --- | --- | --- |
| **Increased abundance in diet B** |  |  |  |
| 1 | A0A4W6EX7 | Transporter | 7.49 |
| 2 | A0A4W6ER2 | Ferritin | 4.61 |
| 3 | A0A4W6D6I | Peptidyl-prolyl cis-trans isomerase (PPIase) | 3.64 |
| 4 | A0A4W6D1Z | Adhesion G protein-coupled receptor L2b, tandem | 3.54 |
| 5 | A0A4W6EE9 | Myosin X, like 1 | 3.42 |
| 6 | A0A4W6ETZ | Protein tyrosine phosphatase receptor type Nb | 3.42 |
| 7 | A0A4W6FSI9 | LIM zinc-binding domain-containing protein | 3.39 |
| 8 | A0A4W6EUJ | 3-oxoacyl-(acyl-carrier-protein) synthase | 3.33 |
| 9 | A0A4W6FB2 | NADH dehydrogenase (ubiquinone) iron-sulfur | 3.25 |
| 10 | A0A4W6E7 | Sodium channel subunit beta-2 isoform X1 | 3.18 |
| **Decreased abundance in diet B** |  |  |  |
| 1 | A0A4W6DC | Guanine nucleotide-binding protein subunit alpha | 0.02 |
| 2 | A0A4W6EG | 1-phosphatidylinositol 4,5-bisphosphate | 0.07 |
| 3 | A0A4W6DL | Cell adhesion molecule 1a | 0.08 |
| 4 | A0A4W6CD | Spermine synthase | 0.08 |
| 5 | A0A4W6DX | Guanine nucleotide binding protein (G protein) | 0.09 |
| 6 | A0A4W6CN | ATP synthase subunit beta (EC 7.1.2.2) | 0.09 |
| 7 | A0A4W6BV | Small ribosomal subunit protein uS2 | 0.09 |
| 8 | A0A4W6DBS | ADP ribosylation factor like GTPase 3b | 0.10 |
| 9 | A0A4W6CBE | Solute carrier family 17 member 6 | 0.15 |
| 10 | A0A4W6DB | Uncharacterized protein LOC108891021 | 0.16 |

\*Values > 1.3 indicate more abundant in diet B, values < 0.76 indicate less abundant in diet B

Peptidyl-prolyl cis-trans isomerase (PPIase) (EC 5.2.1.8) increased in the brain, which is involved in immune regulation [24]. This suggests an adaptive response to different dietary compositions. Protein tyrosine phosphatase receptor type Nb (PTPRN) was increased in the brain. This family of proteins are highly expressed in brain regions like the hypothalamus and pituitary [25]. Notably, PTPRN is required for normal accumulation of dopamine and serotonin in the brain and is involved in insulin and peptide hormone secretion in endocrine cells [26]. An increase in PTPR-Nb in fish fed on diet B could signal altered neuroendocrine activity, such as adjustment of hormone or neurotransmitter release.

Another protein increased in the brain is 3-Oxoacyl-(Acyl-Carrier-Protein) Synthase which is an enzyme involved in the fatty acid biosynthesis pathway [27]. Even though the crude lipid ratio and fatty acids in both diets are similar, total diet composition might have impacted on the brain by, for example, increasing the expression of protein tyrosine phosphatase receptor type Nb to increase fatty acid synthesis (FAS) to compensate for the required fatty acids locally. Indeed, brain FAS is sensitive to diet, and is regulated by nutrient availability that increases in states of excess energy or certain deficiencies [28]. FAS activity in the hypothalamus is known to influence feeding behaviour by producing lipid signals that activate peroxisome proliferator-activated receptor alpha [28].

Several proteins involved in G-protein coupled receptor (GPCR) signaling were markedly lower in abundance in brain of the fish fed on diet B. Notably, a Guanine nucleotide-binding protein Gα subunit and another G-protein component were reduced, alongside a phosphatidylinositol 4,5-bisphosphate phosphodiesterase. These decreases suggest an overall dampening of GPCR signal transduction in the brains of fish fed on diet B. G-proteins (Gα, β, γ) transmit signals from GPCRs, and phosphatidylinositol 4,5- bisphosphate phosphodiesterase enzymes generate second messengers downstream of certain GPCRs. Diet can strongly influence neuromodulatory systems; for example, diet imbalances or difference sources might reduce the activity of neurohormonal pathways. GPCRs are central to regulating appetite, satiety, and metabolism [29]. The observed effect may be linked to subtle nutritional differences between Diet A and Diet B. These differences could have modulated neurohormonal pathways that regulate feeding behaviour.

Cell adhesion molecule 1a (CAM1a) is another protein that was significantly reduced in the brain of fish fed diet B, implying reduced cell–cell adhesion in the brain. Cell adhesion molecules are crucial for synapse formation, maintenance, and neural plasticity [30]. A decrease in CAM could mean that fish fed on diet B had fewer or less stable synaptic contacts. One possible factor is the differences in the absorption of dietary fatty acids like omega-3 fatty acids which are known to promote synaptic protein expression and plasticity [31].

#### 3.4.2 Functional annotation of the most enriched GO terms in brain

Gene ontology (GO) terms of the DAPs in the brain tissue, illustrated in figure 3 (full details in Supplementary data S3), indicate that fish fed on diet B experience the downregulation of biological pathways such as adenosine triphosphate (ATP) metabolic process, glycolytic, and carbohydrate process.

**Figure 3.**
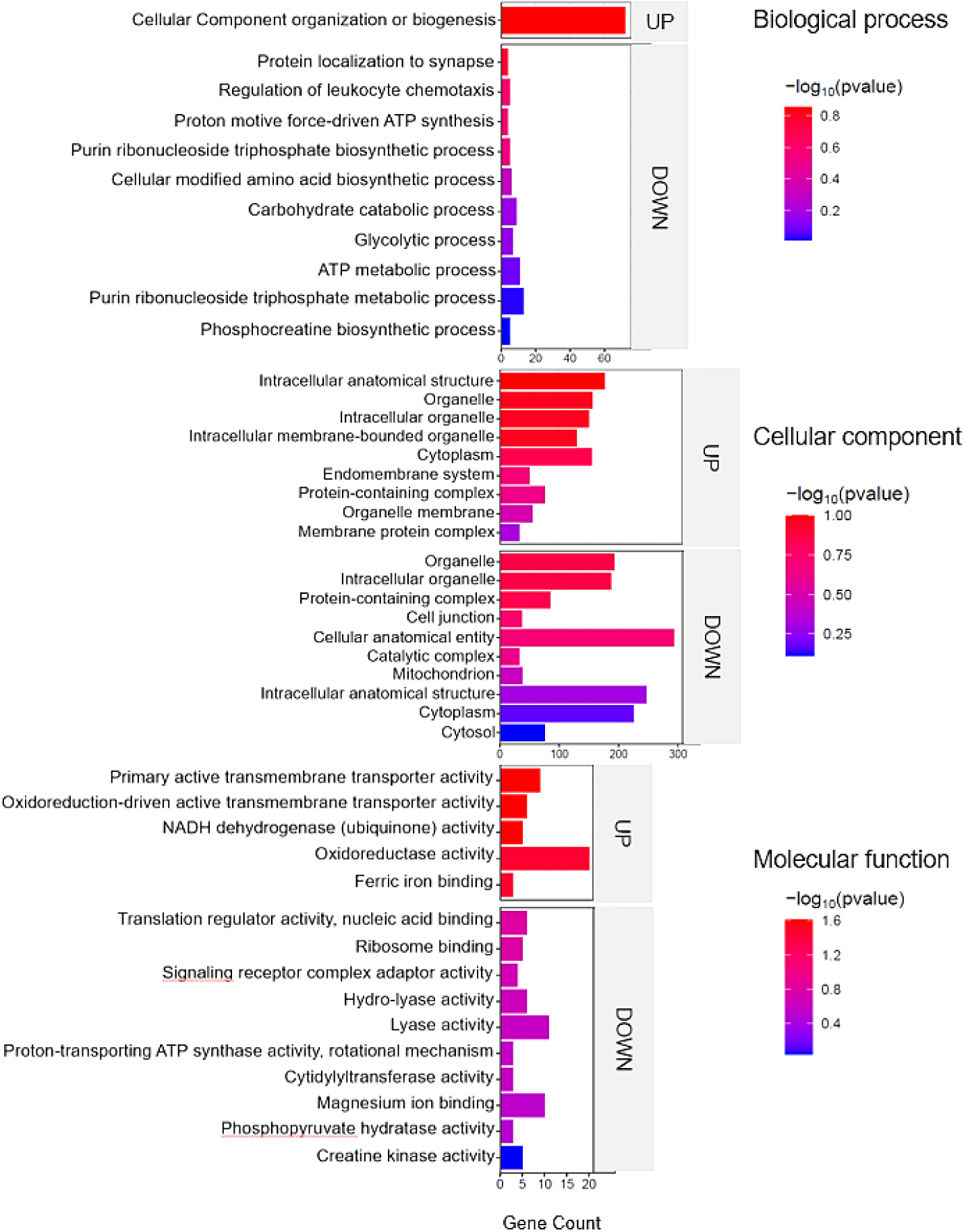
Top enriched GO terms in the DAPs of brain tissue from barramundi. A higher –log₁₀(p-value) (red) indicates greater statistical significance, while lower –log₁₀(p-value) (blue) values indicate less significant enrichment. Up indicates increased abundance in the brain of fish fed on diet B, while Down indicates decreased abundance in the brain of fish fed on diet B.

The downregulation of these pathways points to reduced energy metabolism and cellular renewal. Given the comparable diets and absence of significant physiological differences, this effect may reflect subtle nutritional differences are only detectable at the proteomic level [32]. This reduction in ATP-related processes often accompanies oxidative stress or inflammation, as organisms may divert energy toward repair and immune functions instead of growth [32]. Therefore, growth rate is suppressed, which is a well-known adaptation to poor nutrition. Inadequate nutrient intake in fish typically results in slower growth and smaller body size, as the fish conserves energy [33]. This is a common outcome of nutritional condition, reflected by poor weight gain and low feed efficiency in deficient diets [33]. Although no significant physiological differences were observed, the proteomic shifts in fish fed diet B may indicate subtle metabolic adaptations to the altered nutrient composition, representing an early molecular response to the available nutrients.

There is an upregulation of cellular components including intracellular organelle membrane, and cytoplasm, which might reflect cellular remodelling or immune activation. One plausible interpretation is the activation of autophagy. This is a process of organelle membrane formation and cytoplasmic component recycling that happens as an adaptive response to nutrient composition. In fish, autophagy is induced during starvation as a survival mechanism, helping to recycle cellular constituents to provide nutrients to vital organs [34]. This process involves increased formation of double-membrane autophagosomes (derived from organelle membranes) that engulf cytoplasmic components for degradation, leading to the enrichment of membrane and cytoplasm-associated proteins. Such cellular remodelling can also tie into metabolic and adaptive responses in fish [35].

Meanwhile, downregulation of GO terms such as catalytic complex, mitochondrion, and cytosol might indicate a disruption in energy production, aligning with the depressed metabolic pathways noted above. This disruption is often linked to oxidative stress and iron deficiency. For instance, both oxidative stress and iron deficiency can impair mitochondrial function. Iron is a critical cofactor for many mitochondrial enzymes and the electron transport chain; iron deficiency is known to reduce oxidative phosphorylation capacity by limiting iron–sulfur cluster and heme synthesis in complexes I–III of the electron transport system [36]. In severe iron deficiency, mitochondria may even show structural abnormalities (e.g. loss of cristae) that compromise ATP production [36]. Proteomic evidence of mitochondrial protein downregulation in the brains of fish fed on diet B suggests a modulation of mitochondrial energy metabolism, potentially reflecting an adaptive response to subtle nutritional differences between the diets. Although physiological parameters did not differ significantly and the diets were compositionally comparable, these proteomic patterns indicate early molecular adjustments associated with variations in nutrient availability.

Upregulation was also observed of molecular functions including oxidative activity, nicotinamide adenine dinucleotide hydride (NADH) dehydrogenase, and active transmembrane transporter activity in the fish fed on diet B. These increases suggest that the cells were mounting a defensive response against oxidative responses. Oxidoreductase enzymes include antioxidants and components of the electron transport chain that manage redox reactions. When oxidative response is elevated (for example, due to iron-mediated reactive species oxygen (ROS) or inflammation), organisms often induce antioxidant enzymes to counteract the damage. Indeed, studies on fish under fasting or stress show a significant upregulation of antioxidant defence proteins like superoxide dismutase (SOD), catalase (CAT), and glutathione peroxidase (GPX) [37]. In one study in Common Dentex (*Dentex dentex*) liver, starved fish increased their SOD, CAT, and GPX activities by 20–50% as a compensatory mechanism to neutralize the excess ROS, although oxidative damage still occurred [37]. The observed rise in oxidoreductase activity in fish fed on diet B aligns with this pattern. It likely reflects the fish activating antioxidant and redox-balancing pathways to cope with the oxidative imbalances. For example, the enhancement of NADH dehydrogenase (a component of mitochondrial Complex I) might indicate stimulated electron transport chain activity, potentially to maintain ATP production to regenerate NAD⁺ for metabolic balance. Similarly, increased transmembrane transporter activity could be related to transporters involved in ion homeostasis and nutrient uptake, or export of toxic byproducts, which are often upregulated during cellular strain to restore homeostasis.

#### 3.4.3 Energy metabolism and ferroptosis

Figure 4 shows the most enriched KEGG pathways of DAPs in brain tissue. Oxidative phosphorylation (highlighted in Figure 4) is a cellular process that utilizes the reduction of oxygen to produce high-energy phosphate bonds in the form of ATP. This process involves a series of oxidation-reduction reactions where electrons are transferred from NADH and the reduced form of flavin adenine dinucleotide (FADH2) to oxygen. These reactions occur within various protein, metal, and lipid complexes in the mitochondria, collectively known as the electron transport chain (ETC). The ETC uses NADH and FADH2, which are generated from different catabolic processes within the cell [38]. However, electrons can leak from the cell and react with oxygen and produce superoxide anions. Excessive amounts of superoxide anions as a source of ROS can result in the oxidation of biological molecules such as lipids, proteins, and DNA [39].

**Figure 4.**
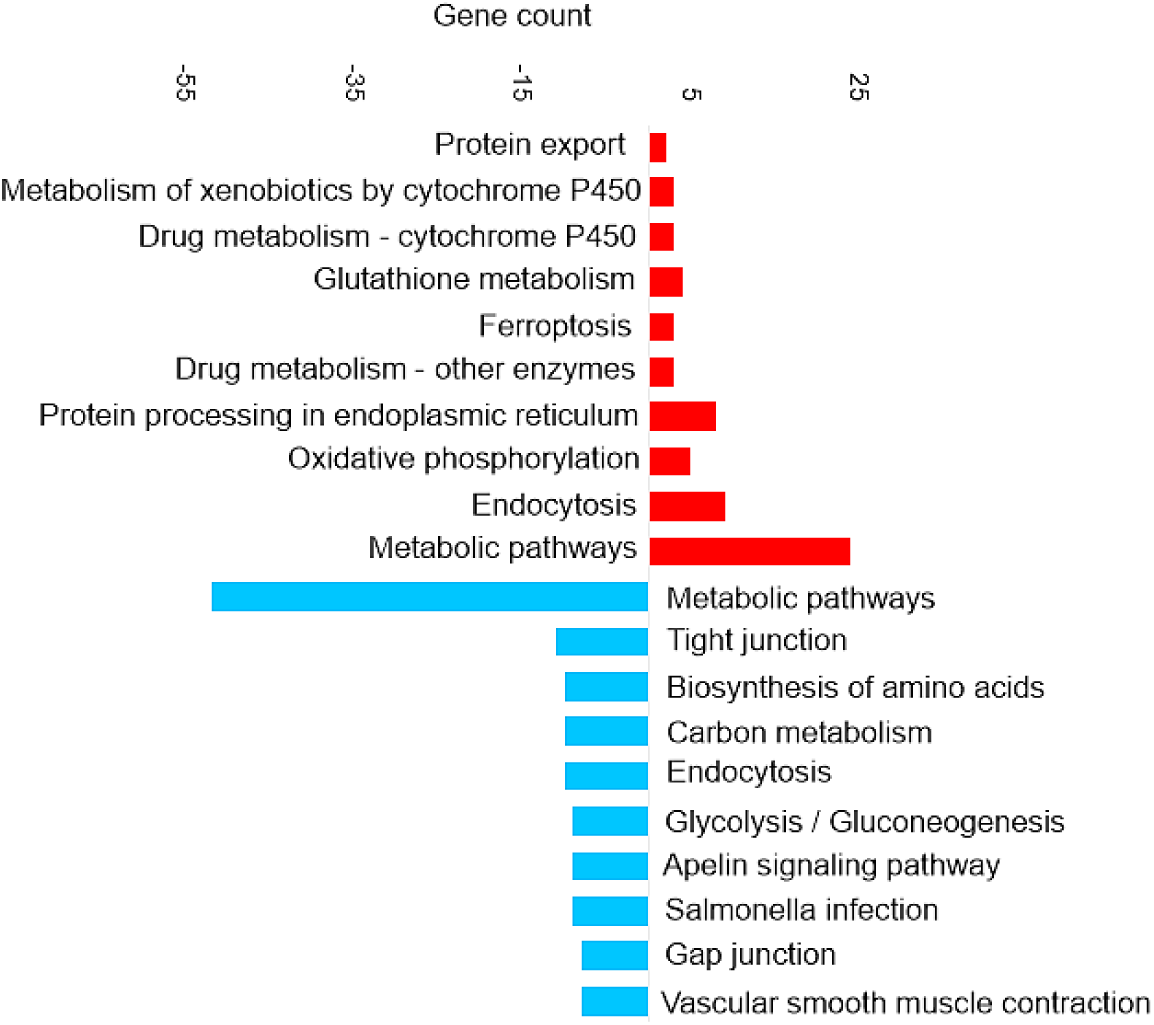
Top KEGG pathways in the differentially abundant proteins in brain tissue

Ferroptosis upregulation (highlighted in Figure 4) in the brain is related to the increase of ferritin. Ferroptosis was identified as an iron-dependent form of programmed cell death [40]. The unique process of ferroptosis includes the dysregulation of iron metabolism and the accumulation of ROS [41,42]. Some characteristics of ferroptosis are cytological changes, including decreased or vanished mitochondria cristae, a ruptured outer mitochondrial membrane, and a condensed mitochondrial membrane [43].

In the present study, although ferroptosis-related proteins were upregulated in fish fed on Diet B, there were no significant differences in physiological parameters or feed intake between the two dietary groups. This suggests that the observed ferroptosis activation represents an adaptive regulatory response to subtle nutritional differences, rather than a pathological outcome. Such molecular adjustments likely reflect normal cellular mechanisms maintaining iron and redox homeostasis under slightly varying dietary conditions.

#### 3.4.4 Glutathione metabolism and antioxidant response

The Glutathione metabolism KEGG pathway was also upregulated (highlighted in Figure 4), which includes the increase of glutathione transferase (EC 2.5.1.18). Glutathione (GSH) is a critical endogenous antioxidant found in all eukaryotic cells [44]. Increasing the supply of cysteine or its precursors via oral or intravenous administration enhances GSH synthesis and prevents GSH deficiency in humans and animals under various nutritional and pathological conditions [45]. Cysteine depletion can trigger iron-dependent nonapoptotic cell death – ferroptosis [46]. Even though cysteine itself is not an essential amino acid, methionine is the metabolic precursor for cysteine [47]. The higher essential amino acid content of methionine in diet B may have induced compensatory responses in the brain, as reflected by the upregulation of glutathione metabolism. This shift likely represents an antioxidant strategy to counter ROS accumulation.

#### 3.4.5 Appetite regulation and digestive response

The downregulation of certain KEGG pathways in the brain, such as apelin signalling (highlighted in Figure 4) may represent compensatory mechanisms, attempting to mitigate the effects of dietary composition and slightly lower food intake. Apelin is a food intake-regulating peptide [48]. Guanine nucleotide-binding protein subunit alpha is one of the proteins involved in the apelin signaling pathway that decreased. Research on common crab has shown that Pyr-apelin-13 supplementation can promote food intake and growth by regulating the mRNA expression levels of key genes [49]. Apelin has been shown to influence appetite by interacting with regulatory signals such as leptin and ghrelin. Additionally, apelin levels are elevated in obesity disorders associated with hyperinsulinemia, conditions where feeding behaviour and energy balance are disrupted. These observations suggest that apelin might play a role in regulating feeding behaviour and maintaining energy homeostasis [48]. It has been shown previously that infusion of apelin reduced the juice volume, protein and trypsin outputs in a dose-dependent manner [48]. Trypsin is one of the serine proteinases in fish viscera that breaks down the protein [50]. KEGG pathway analysis suggests that the dietary composition might include compounds that impact the digestibility or bioavailability of the nutrients by reducing the trypsin activation. For example, rats fed raw soybeans showed decreased body weight and increased pancreas weight that is due to the presence of trypsin inhibitors in the soybean since soybean can increase the release of cholecystokinin (CCK) [51]. Trypsin as a digestive proteinase has been shown to reduce the release of CCK [51], which is a peptide hormone and a neurotransmitter expressed mainly in a subpopulation of small intestinal endocrine cells (I-cells) and in cerebral neurons [52]. CCK is believed to make the rat feel full, and higher levels of CCK might lead to eating less [51]. In this study the nutritional profile for both diets show similar level of faba bean which has trypsin inhibitor activity [53].

### 3.5 Statistical analysis of protein expression in liver

Proteomic profiling coupled with multivariate analysis is a powerful approach to discern dietary effects on fish liver physiology. In the present context, PLS-DA analysis was applied to highlight diet-driven clustering of liver proteomes, a strategy similarly utilized in other fish nutrition omics studies [54]. The PLS-DA plot of proteins identified in the liver samples (Figure 5) shows a distinct separation between the two dietary groups along the first two principal components. The ellipses representing the 95% confidence intervals are well separated, indicating a high degree of discriminative power between diet A and diet B at the proteomic level in the liver. Samples from diet A are tightly clustered together and distinctly separated from the diet B samples, which also form a cohesive cluster. The results underscore that the fish liver proteome is highly responsive to dietary composition, which induces marked shifts in hepatic protein expression profiles [55,56].

**Figure 5.**
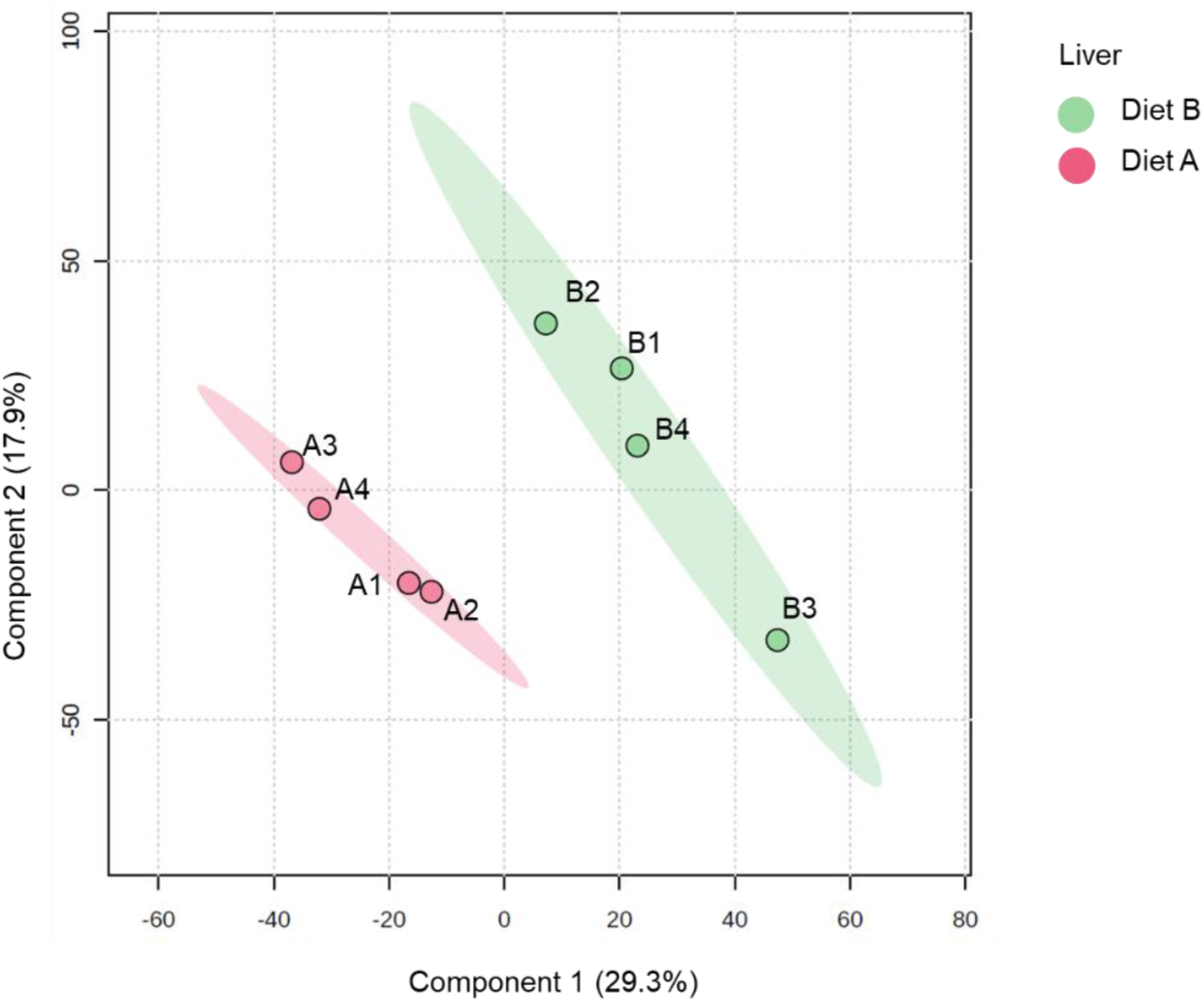
Partial least squares discriminant analysis (PLS-DA) of liver of barramundi fed on diet A and diet B displayed at 95% confidence intervals highlights the clustering of the sample based on their response to different diets (n= 4).

The volcano plot provided presented in figure 6 shows the protein expression in the liver tissue. For liver, 231 proteins were decreased, and 235 proteins were increased in the liver of fish fed on diet B, as shown in table 3. Full details of all DAPs in liver are presented in Supplementary data S4. Notably, these diet-induced proteomic changes are similar to a previous study on Spotted scat (*Scatophagus argus*) where dietary supplementation of oil applied showed a significant differential protein expression in the liver between groups [57].

**Figure 6.**
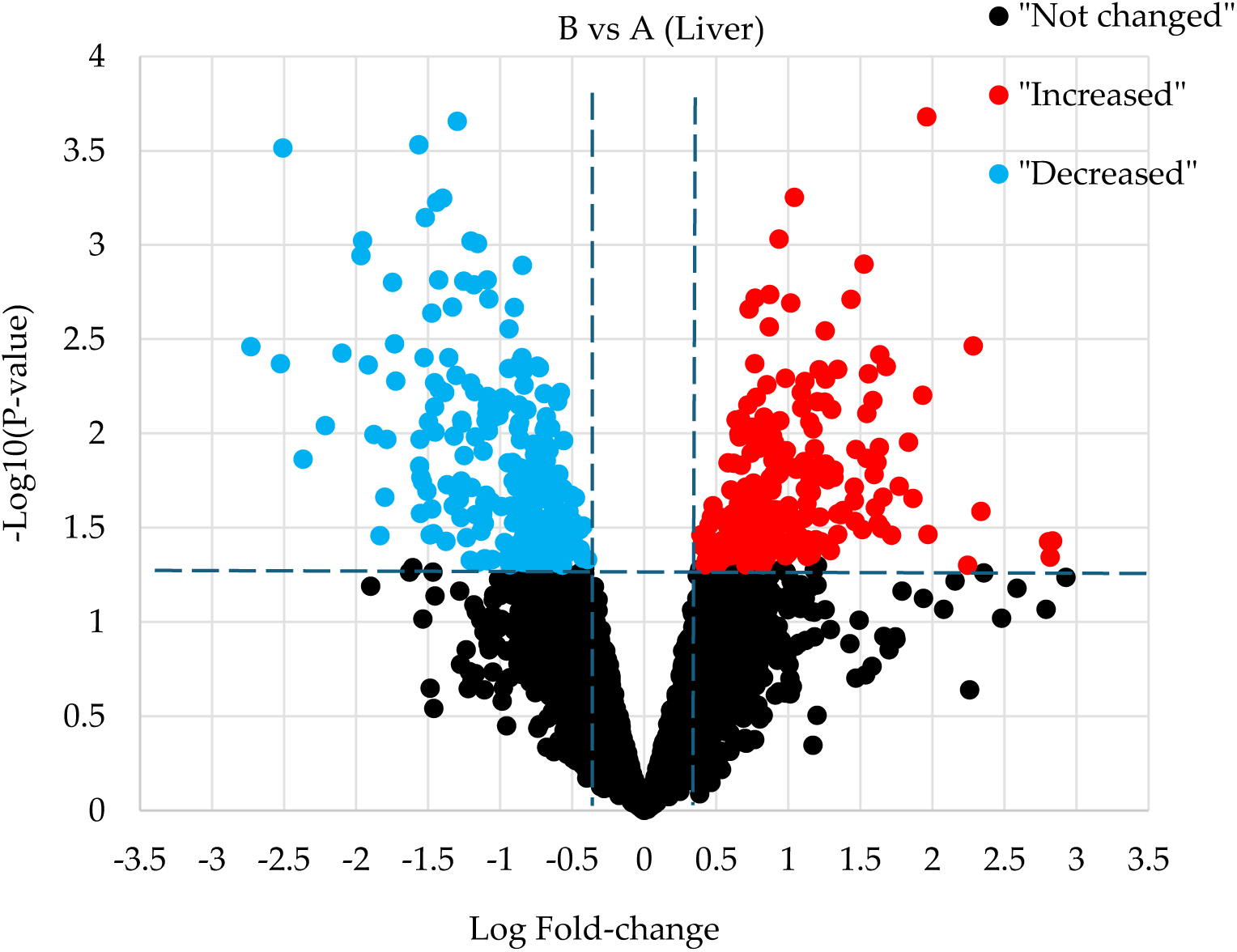
Volcano plot illustrates significantly differentially abundant proteins in liver. The −log10 (P value) is plotted against the log2 (fold change: diet B/diet A).

#### 3.5.1 Differentially abundant proteins in liver

The liver is the primary site where absorbed nutrients are processed after leaving the gut. It plays a central role in metabolizing carbohydrates, fats, and proteins; activating and storing vitamins; and detoxifying and eliminating both internal and external compounds [58]. Table 6 presents the top 10 DAPs, sorted by fold change, in the liver of fish when comparing the different dietary formulations.

**Table 6.**
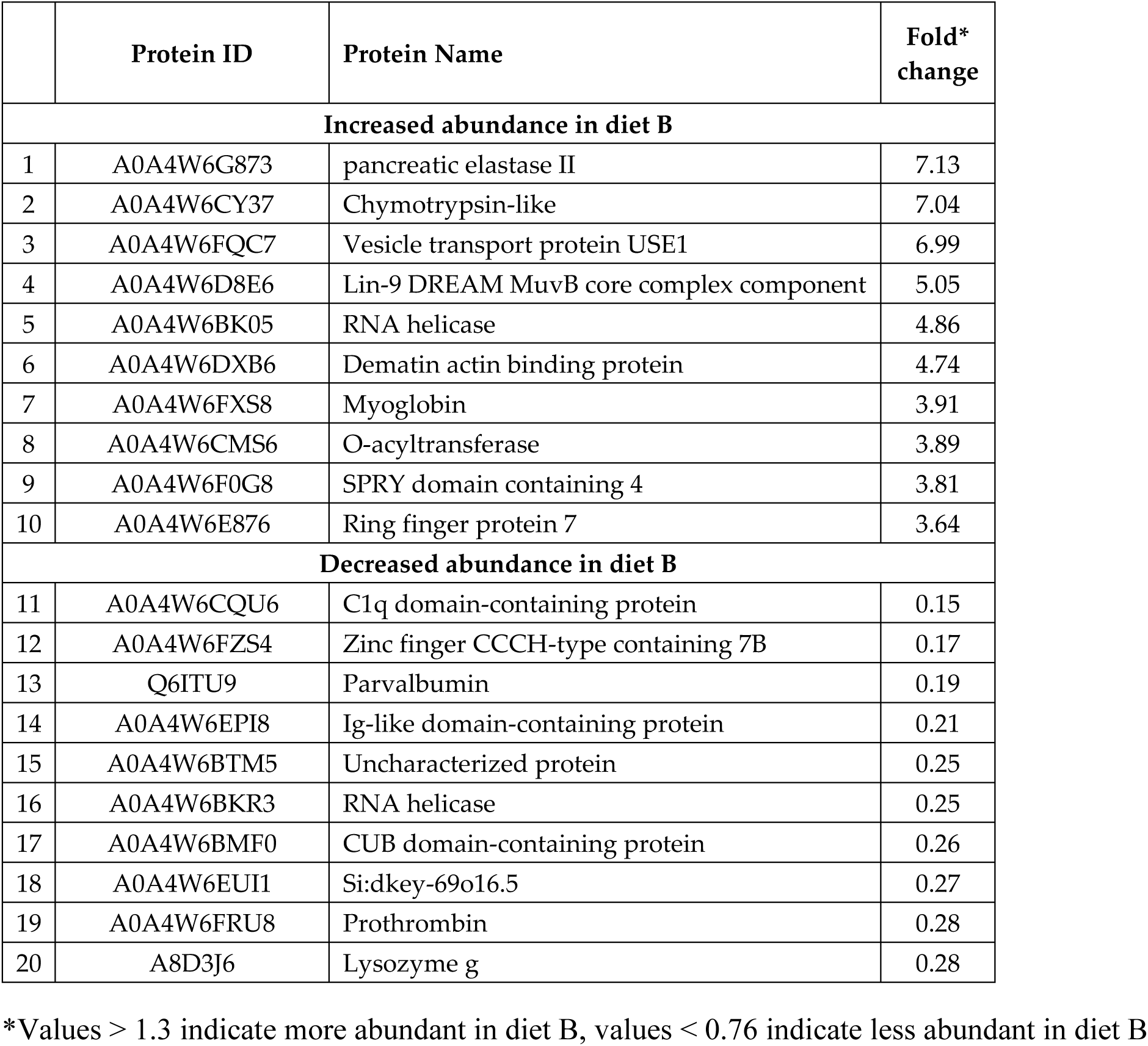
Top ten DAPs in the Liver tissue

#### 3.5.2 Proteolytic enzymes and nutrient digestion

The synthesis of pancreatic enzymes and their secretion into the duodenum are essential for digestion of dietary macromolecules and absorption of nutrients [59]. In particular, the activities of proteolytic enzymes are modified either by the protein level and the source of protein in the diet or by food intake [60,61]. Our results are similar to these findings, as Table 6 shows an increase in the amount of pancreatic elastase II and chymotrypsin. Elastase II, which is closely related to the chymotrypsin family [62], exhibits a broad specificity for substrates containing medium to large hydrophobic amino acids in the P1 position [59]. Diet B includes a higher proportion of hydrolysed feather meal . Feather meal includes a high proportion of keratin, which has a high concentration of hydrophobic amino acids [63]. Keratin is resistant to digestion by the proteases pepsin or trypsin [64]. The increase in the amount of proteases expressed in the liver of fish fed on diet B could be an adaptive response to these compositional differences to regulate the digestion of the fish.

#### 3.5.3 RNA processing and cellular response

RNA helicase (DDX5) is another protein (highlighted in Table 6) that was increased in the liver with a fold change of 4.86. DDX5 (also known as p68) is one of the prototypic members of the DEAD box family of RNA helicases. DDX5 and related DDX17 (p72) are involved in a variety of cellular processes, including transcription, pre-mRNA and rRNA processing, alternative splicing and miRNA processing, and they are also dysregulated in a range of cancers [65–68]. The RNA helicase DDX5 functions as a coactivator of the tumor suppressor protein p53, playing a crucial role in the DNA damage response by facilitating the transcription of genes like CDKN1A (p21), which are essential for cell cycle arrest [69,70]. However, studies have shown that while DDX5 is necessary for p53-dependent induction of p21 and subsequent cell cycle arrest, it does not participate in the activation of pro-apoptotic genes [70]. This suggests that DDX5 may influence the cellular decision to undergo cell cycle arrest rather than apoptosis in response to DNA damage, as a result of conditions triggered by ferroptosis, thereby contributing to cell survival under certain conditions. Lysozyme g was also decreased in the liver of fish fed on diet B that contains more methionine, which is an important secretory innate immune system component [71]. Lyzozyme activities were influenced when *Scophthalmus maximus* were stressed [72]. In European Seabass, fish fed with methionine supplementation showed an increase in the level of lysozyme, and a higher survival was observed in fish fed with supplemented diets [73].

#### 3.5.4 Functional annotation of the most enriched GO terms in liver

Gene ontology (GO) terms of the DAPs in the liver tissue are illustrated in figure 7, with full details provided in Supplementary data S5. GO enrichment analysis of liver tissue revealed changes in key biological pathways. In the biological process category, both increased and decreased proteins were predominantly associated with metabolic processes, including cellular metabolic processes, small molecule metabolic processes, nitrogen compound metabolism, and biosynthetic pathways. The cellular component analysis showed that increased proteins were enriched in ribosomal subunits, endoplasmic reticulum protein-containing complexes, and vesicle transport components. In contrast, decreased proteins were linked to the cytoplasm, proteasome complex, mitochondria, and extracellular space. In the molecular function category, increased proteins were associated with catalytic and hydrolase activity as well as ribosome structural components, while decreased proteins were related to peptidase and endopeptidase inhibitor functions. These findings suggest that dietary composition modulates liver function by altering metabolic, biosynthetic, and proteolytic pathways.

**Figure 7.**
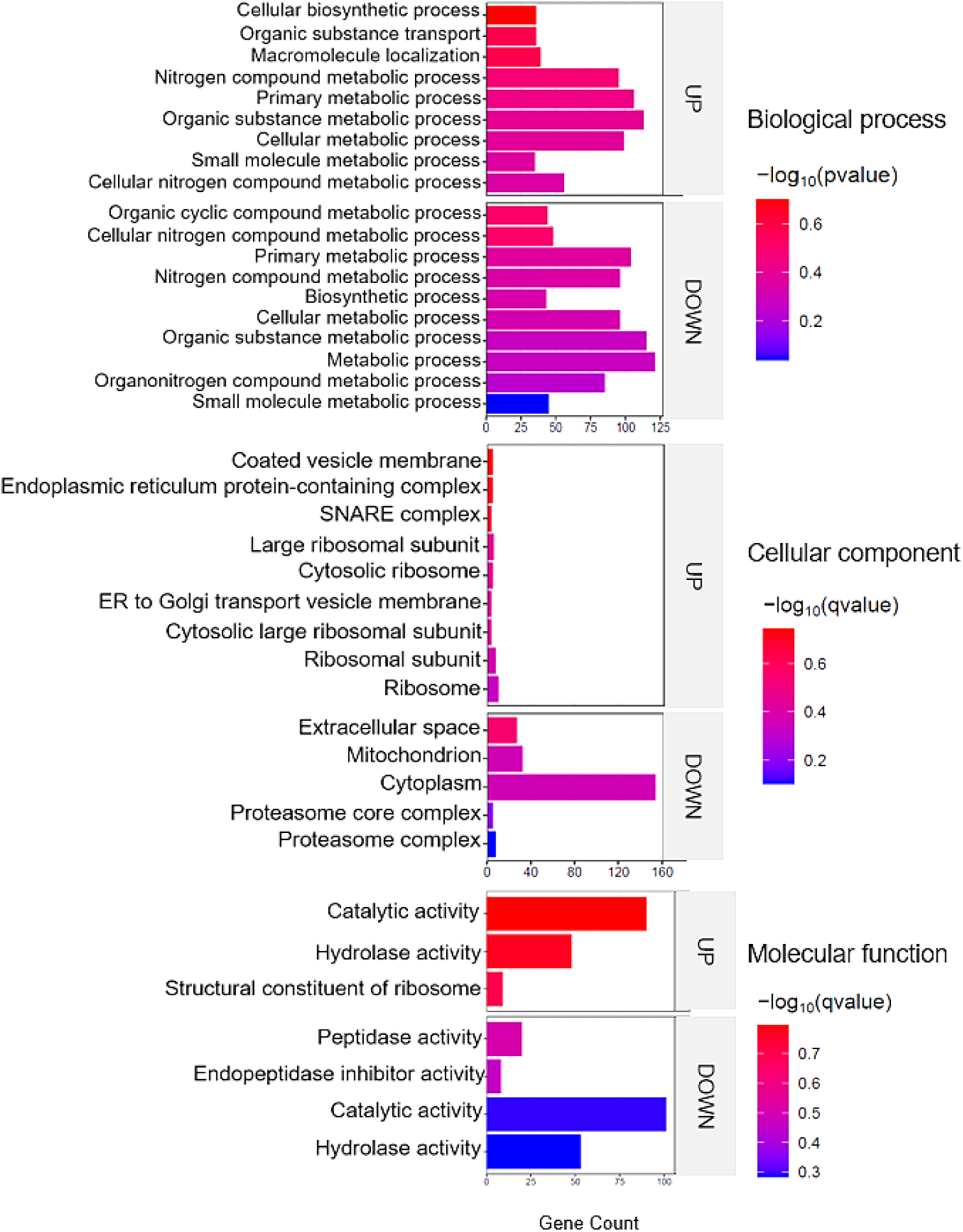
Top enriched GO terms in the DAPs of liver of barramundi. A higher –log₁₀(p-value) (red) indicates greater statistical significance, while lower –log₁₀(p-value) (blue) values indicate less significant enrichment. Up indicates increased abundance in the liver of fish fed on diet B, while Down indicates decreased abundance in the liver of fish fed on diet B.

#### 3.5.5 Ferroptosis and oxidative pathways

Figure 8 shows the most enriched KEGG pathways of DAPs in liver. Ferroptosis and glutathione metabolism are two of the upregulated pathways in liver tissue, as shown in figure 8. This is similar to the results discussed in the brain tissue in section 3.4.3. Proteins involved in the ferroptosis pathway include glutathione peroxidase, glutathione synthetase, Lys phosphatidylcholine acyltransferase 3, and one uncharacterized protein (for details, see Supplementary data S5)

**Figure 8.**
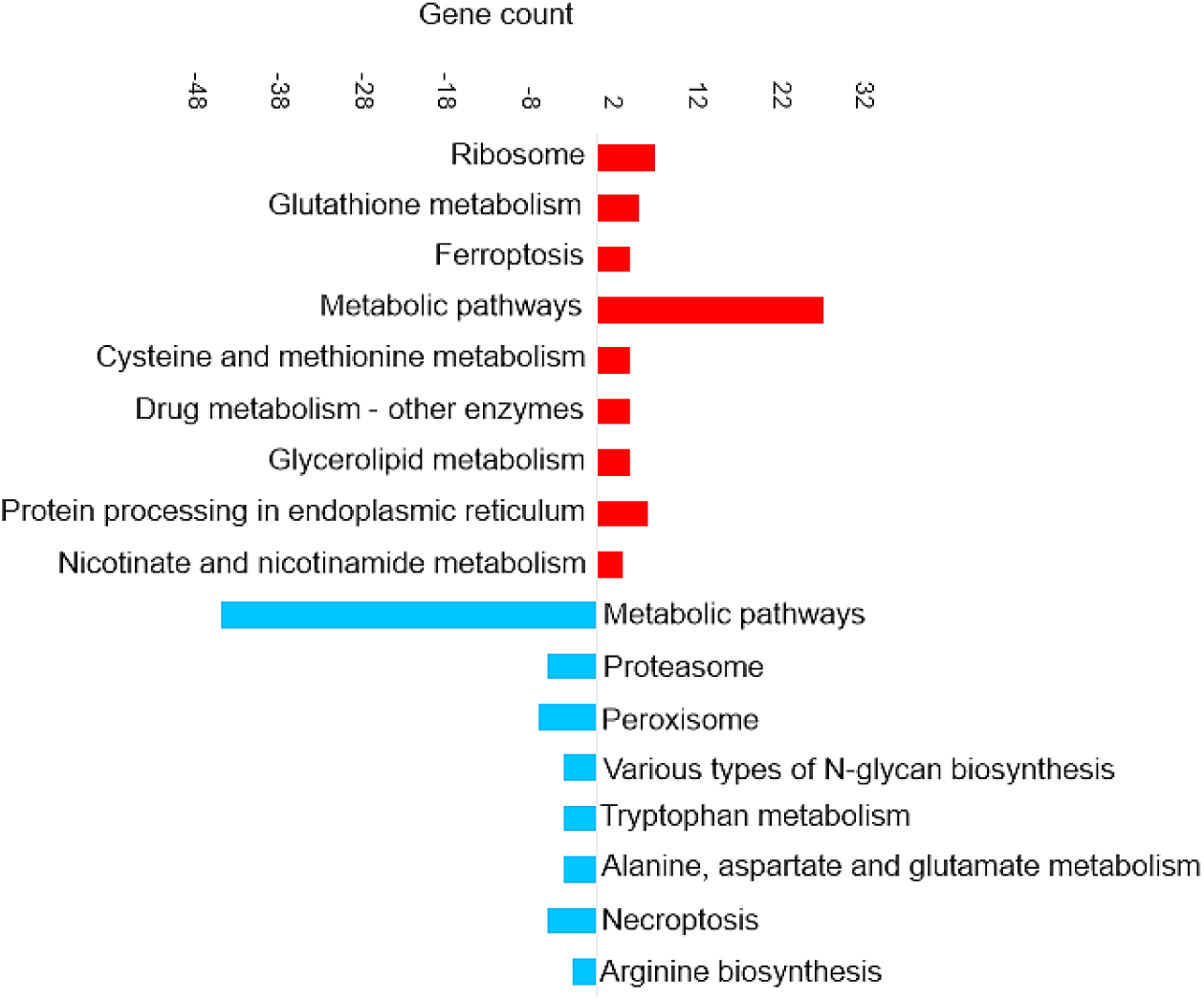
Top KEGG pathways of the differentially abundant proteins in liver tissue

### 3.6 Statistical analysis of protein expression in Intestine

The PLS-DA analysis of the intestine samples (presented in Figure 9) shows a similar separation between the dietary groups as was observed for brain and liver tissues, although the replicates are not as tightly clustered together. Figure 10 shows the differential protein expression in intestine tissue between fish fed on diet B and diet A. The observed fold change values are higher than for the other tissues which may reflect the role of the intestine in nutrient absorption and barrier functions, with involvement in metabolic regulation, indicating tissue-specific sensitivity to dietary interventions. As shown in Table 5, 349 proteins were increased, and 485 proteins were decreased in the intestine of fish fed on diet B when compared with diet A (for details, see Supplementary data S6).

**Figure 9.**
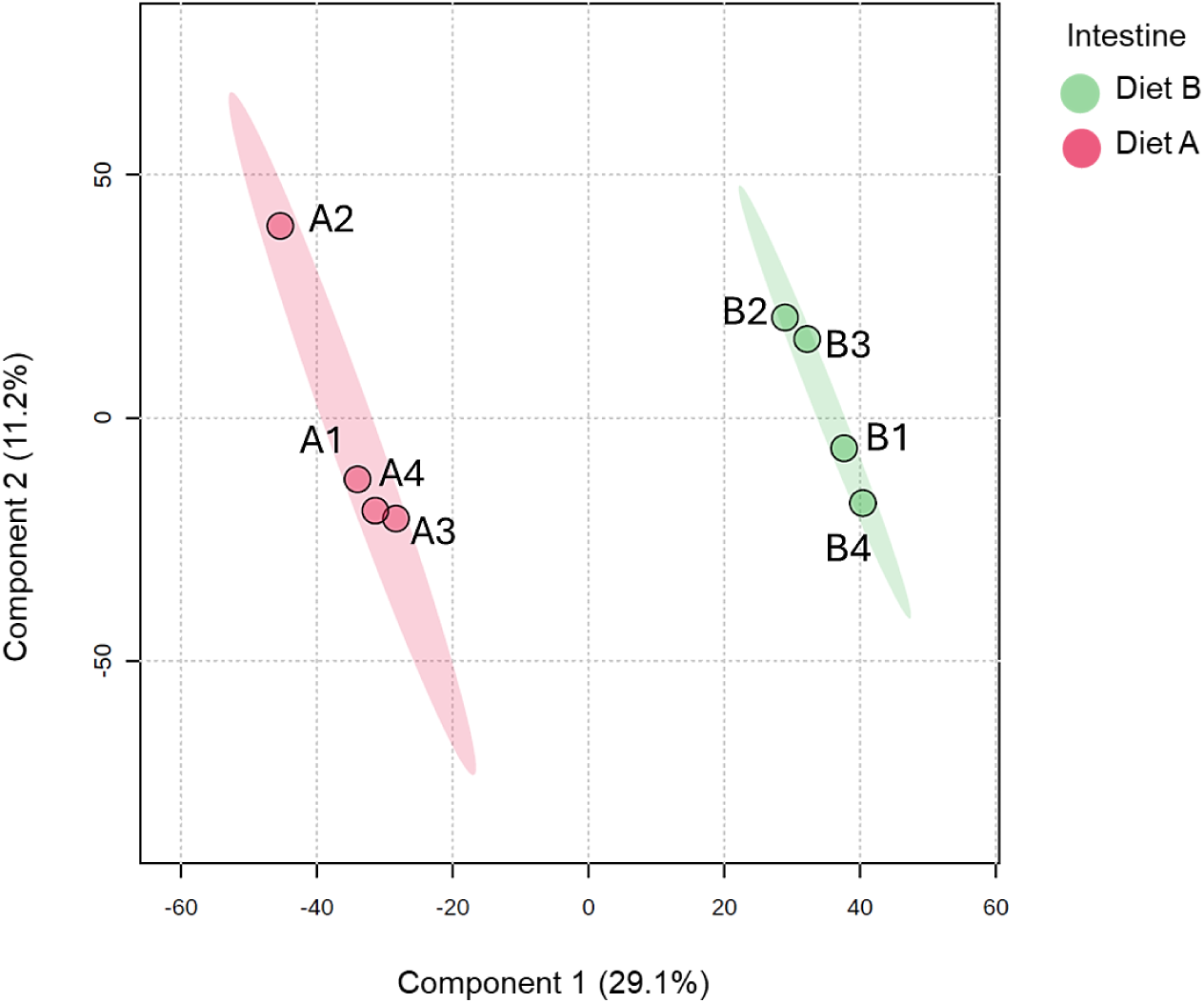
Partial least squares discriminant analysis (PLS-DA) of intestine of barramundi fed on diet A and diet B displayed at 95% confidence intervals highlights the clustering of the sample based on their response to different diets (n= 4).

**Figure 10.**
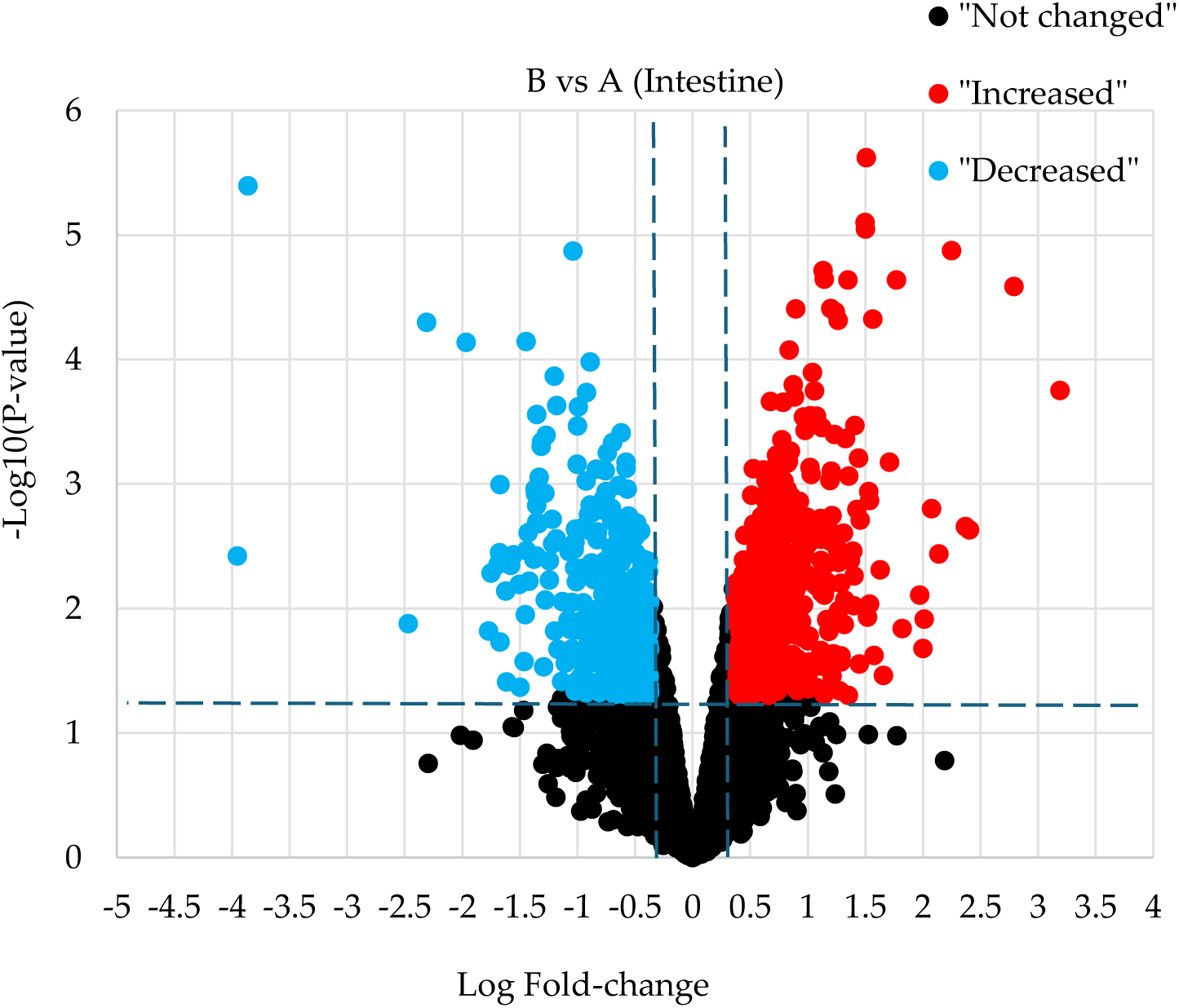
Volcano plot illustrates significantly differentially abundant proteins in intestine. The −log10 (P value) is plotted against the log2 (fold change: diet B/diet A).

#### 3.6.1 Differentially abundant proteins in intestine

The intestine plays a central role in nutrient digestion and absorption, acting as a vital interface between the external environment and the internal physiology of an organism. It is not only responsible for the enzymatic breakdown and uptake of macronutrients such as proteins, lipids, and carbohydrates, but also for the absorption of micronutrients, including vitamins and minerals [74]. The intestinal epithelium is highly specialized, containing a variety of transporters, enzymes, and structural proteins that regulate barrier integrity and nutrient passage [75]. Fish intestines are vital for immunological response, nutrition absorption and environmental interactions [76]. Proteomic profiling of intestinal tissues is therefore a powerful approach to assessing how dietary interventions as an environmental factor affect gut function and nutrient utilization. Table 7 presents the top ten proteins as sorted by fold change which were increased and decreased in the intestine tissue of fish fed on diet B compared with fish fed on diet A.

**Table 7.**
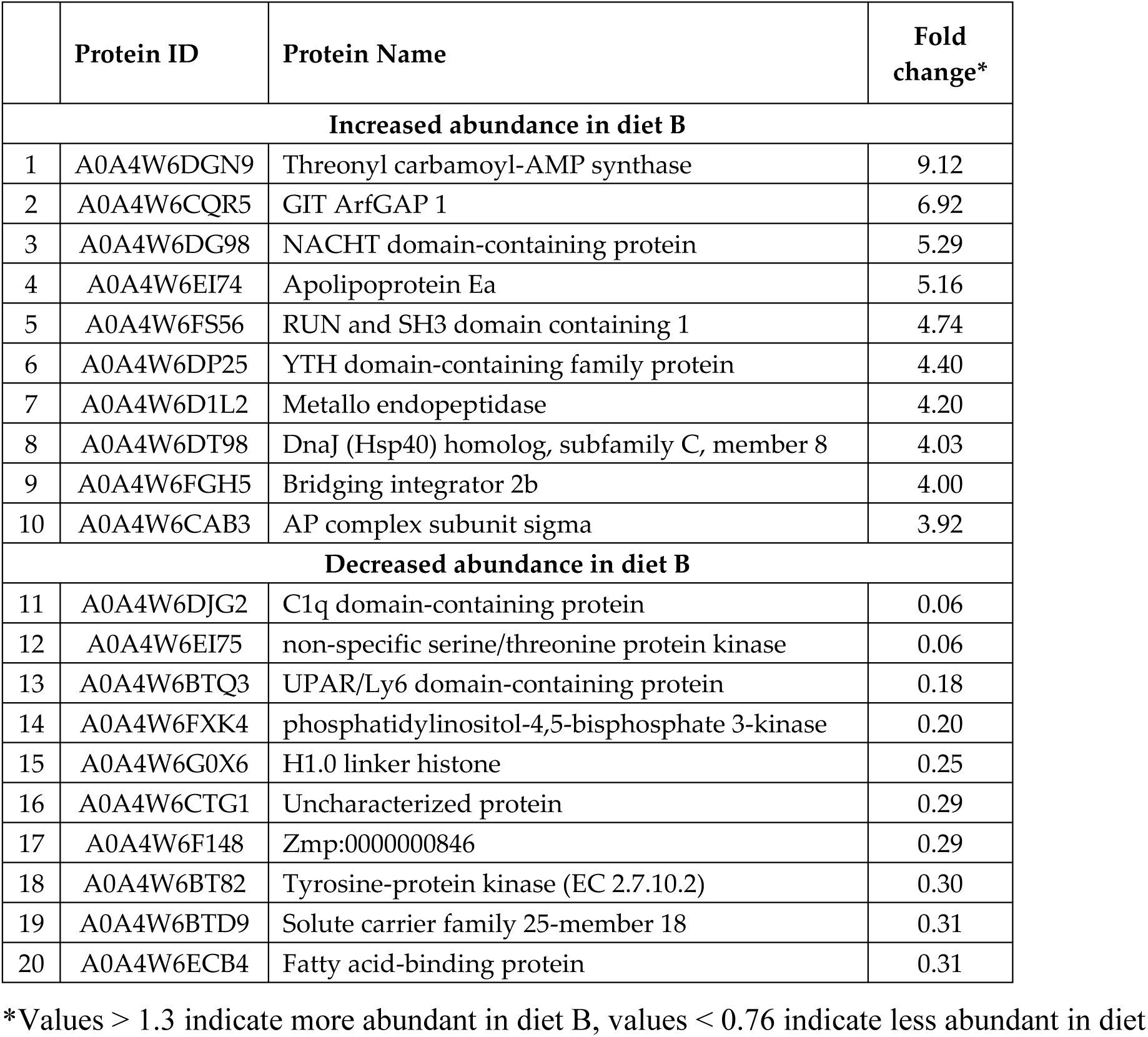
Top ten DAPs in the intestine tissue

#### 3.6.2 Immune signaling and adaptive response

NACHT domain-containing protein (highlighted in Table 7) increased with a fold-change of 5.29. This protein is found in eukaryotic pattern recognition receptors involved in promoting anti-microbial defense, mediating their oligomerization into immune signaling complexes. The NACHT module is an oligomerization domain that facilitates assembly of multiprotein, immune-signaling complexes in response to microbial ligands or disrupted physiological processes [77]. Another protein increased is apolipoprotein Ea (ApoE) with a 5.16-fold change. ApoE is also implicated in infections with herpes simplex type-1, hepatitis C, and human immunodeficiency viruses [78]. Oxidative responses have been shown to augment ApoE secretion from adipocytes, and ApoE overexpression protected cells from hydrogen peroxide-induced damage [79].

#### 3.6.3 Lipid metabolism and nutrient transport

Fatty acid-binding protein (highlighted in Table 7) decreased with a fold change of 0.31. Fatty acid-binding protein is essential for digesting and absorbing dietary fats by transporting bile acids in intestinal cells. It helps break down cholesterol and may be linked to bile acid transport in the intestine [80]. High-fat feeding of intestinal fatty acid-binding protein in mice resulted in reduced weight gain and fat mass relative to wild-type mice [81]. The mRNA levels of fatty acid synthase and sterol regulatory element-binding protein-1 (SREBP1) genes increased with increasing dietary methionine in juvenile cobia [82].

#### 3.6.4 Functional annotation of the most enriched GO terms in intestine

Figure 11 presents the most enriched GO terms represented within the set of DAPs from intestinal tissue. GO enrichment analysis of DAPs in intestine tissue revealed distinct shifts in biological processes, cellular components, and molecular functions.

**Figure 11.**
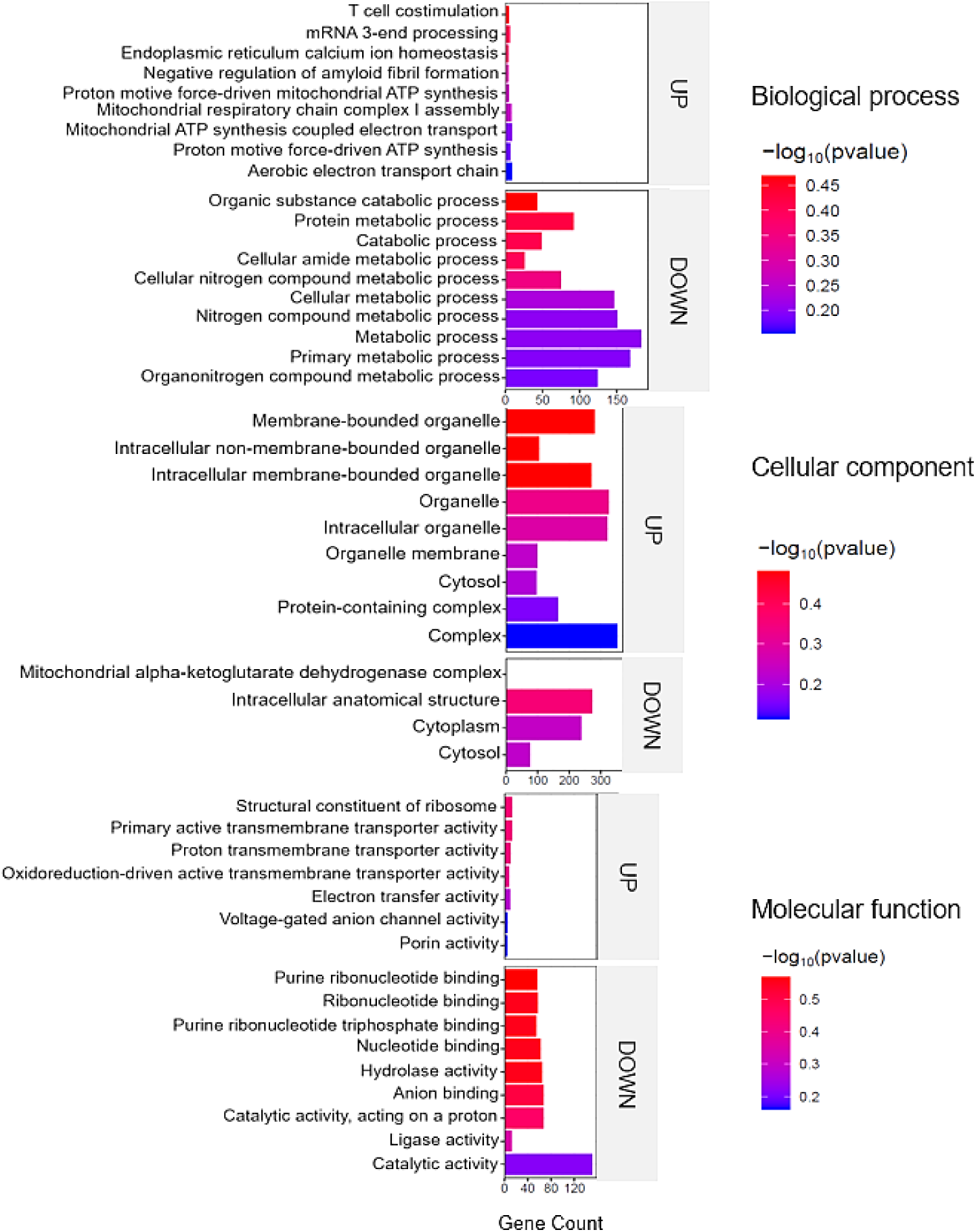
Top enriched GO terms in the DAPS of the intestine of barramundi. A higher –log₁₀(p-value) (red) indicates greater statistical significance, while lower –log₁₀(p-value) (blue) values indicate less significant enrichment. Up indicates increased abundance in the intestine of fish fed on diet B, while Down indicates decreased abundance in the intestine of fish fed on diet B.

Among the upregulated proteins, pathways associated with mitochondrial energy production were significantly enriched, including the aerobic electron transport chain and proton motive force-driven ATP synthesis, suggesting an increased mitochondrial activity and energy demand. Correspondingly, upregulated cellular components were primarily linked to the mitochondrial membrane, protein-containing complexes, and membrane-bounded organelles, reinforcing the role of mitochondria in the observed metabolic response.

In contrast, proteins decreased in abundance were associated with metabolic and catabolic processes, particularly those involving nitrogen-containing compounds, protein metabolism, and primary cellular metabolism. These trends were further supported by the enrichment of cytoplasmic and intracellular organelle components, suggesting a suppression of general metabolic pathways.

At the molecular function level, increased proteins were associated with transmembrane transporter activities, including those involved in electron and proton transport, aligning with enhanced mitochondrial and membrane transport function. Conversely, downregulated functions included catalytic activity, nucleotide binding, and hydrolase activity, indicating a general downregulation of biosynthetic and enzymatic processes. Because ferroptosis recurs in all tissues and promotes ROS generation, it is likely tied to the shifts observed in metabolic proteins. Cells produce ROS in multiple locations including cytoplasm, membranes,

ER, mitochondria, and peroxisomes [83], but mitochondria are the biggest source, creating about 90 % of the total ROS [84].

#### 3.6.5 Amino acid biosynthesis and nitrogen regulation

Figure 12 shows the most enriched KEGG pathways in the set of DAPs in the intestine (full details in Supplementary data S7). Upregulated pathways in the intestine shown in figure 12 included metabolic pathways primarily driven by proteins involved in amino acid biosynthesis and glutathione metabolism. This suggests that diet B triggered a broad metabolic response, with particular emphasis on nitrogen and redox metabolism. Several proteins associated with amino acid biosynthesis such as phosphoserine aminotransferase, ribulose-phosphate 3-epimerase, phosphoglycerate mutase, and argininosuccinate lyase were increased in the intestine. Key proteins in the general metabolic pathways are associated with glutathione metabolism, such as glutathione transferase, glutathione S-transferase kappa, thioredoxin domain-containing protein, and spermidine synthase.

**Figure 12.**
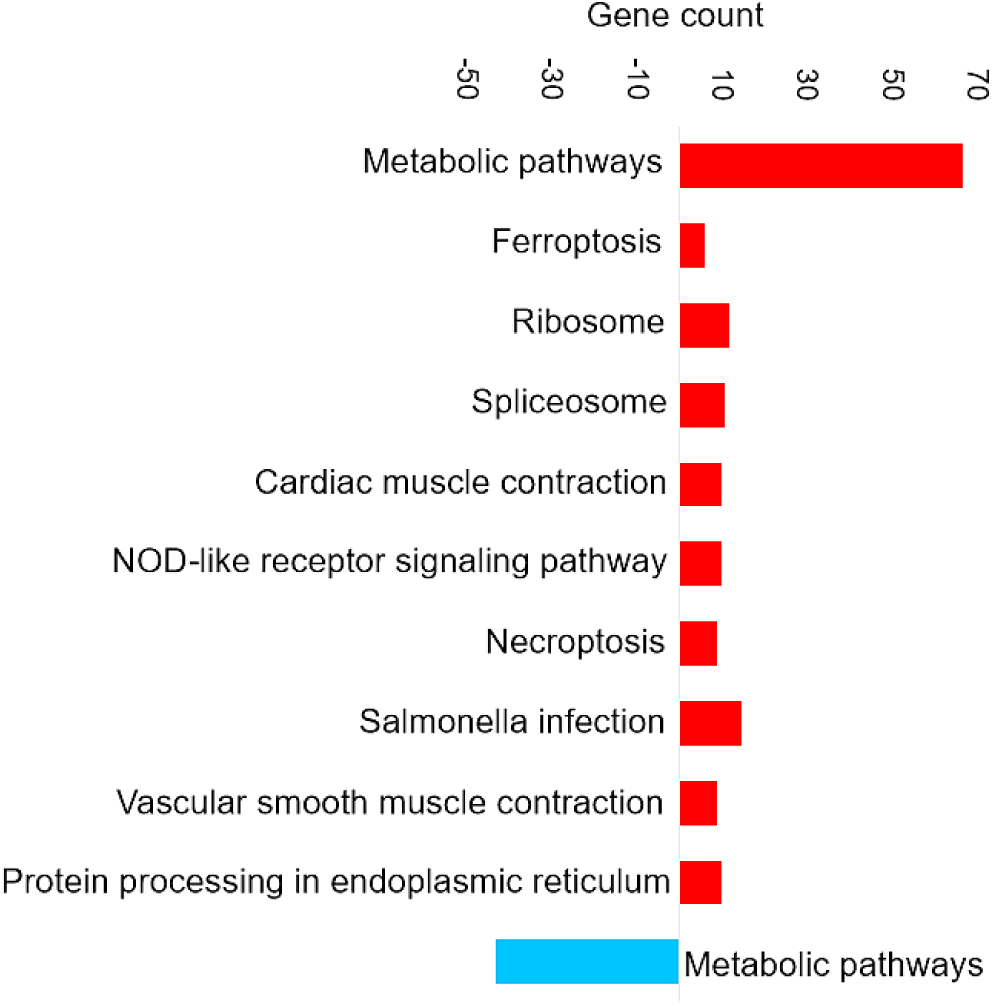
Top KEGG pathways of the differentially abundant proteins in Intestine tissue

Phosphoserine aminotransferase is part of the phosphorylated pathway of serine biosynthesis [85]; serine is a non-essential proteinogenic amino acid that serves as a precursor for glycine, tryptophan, and cysteine [86]. In addition to its role in protein synthesis, serine is a key intermediate in the production of phospholipids and nucleobases [87]. Under environmental conditions in plants, the activity of this phosphorylated pathway is alleviated which suggests that supplying serine is important for non-photosynthetic cells under harsh conditions [88].

Argininosuccinate lyase is involved in the urea cycle, in which argininosuccinate lyase catalyses the fourth reaction in this cycle, resulting in the breakdown of argininosuccinate lyase to arginine and fumarate [89]. Fish have particularly high requirements for dietary arginine, because it is abundant in proteins as a peptide bound amino acid and in tissue fluid as phosphoarginine, a major reservoir of ATP, while its de novo synthesis is limited or even completely absent [90]. Arginine also plays a crucial role in regulating the endocrine system, which releases hormones directly into the bloodstream [90]. The teleost endocrine system includes the organs and tissues that produce hormones which, together with the central nervous system (hypothalamus), control and regulate various body functions. These include sexual activity and reproduction, growth, osmotic pressure, and general metabolic activities such as the storage of fat and the utilization of foodstuffs, blood pressure, and specific aspects of skin colour [91]. Although feed intake was slightly lower in diet B, this did not appear to limit amino acid availability. Arginine may regulate endocrine pathways and could have contributed to the improved feed conversion ratio observed. In addition, amino acid metabolism is central to iron and lipid homeostasis and redox balance, and thus may play a role in modulating ferroptosis [92].

### 3.7 PRM Validation

A series of PRM experiments were performed to validate the results from label-free shotgun proteomics analysis of Barramundi tissues. The PRM results indicated that the differential changes in abundance of nine selected proteins measured by PRM, as shown in Figure 13, 14 and 15, agreed with the label-free shotgun proteomics results. Quantification was based on the integrated peak areas of the most intense y- and b-type fragment ions selected by Skyline from the spectral library for each peptide. The list of monitored precursor ions, fragment ions (transitions), and their corresponding m/z values are provided in Supplementary data S8.

**Figure 13.**
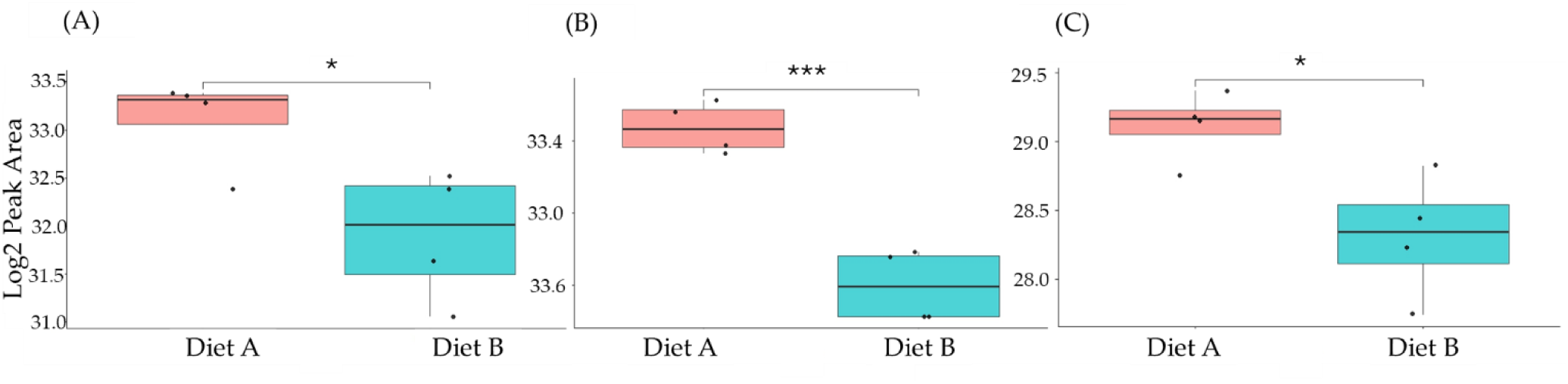
PRM validation of three proteins changed in abundance in response to dietary composition in brain tissue in Barramundi. (A) = Guanine nucleotide binding protein (G protein), (B) = ATP synthase subunit beta (EC 7.1.2.2), (C) = ADP ribosylation factor like GTPase 3b. Asterisks indicate a statistically significant difference between diet A and diet B, according to a Student t-test (* = p-value ≤ 0.05, ** = p-value ≤ 0.01, *** = p-value ≤ 0.001).

**Figure 14.**
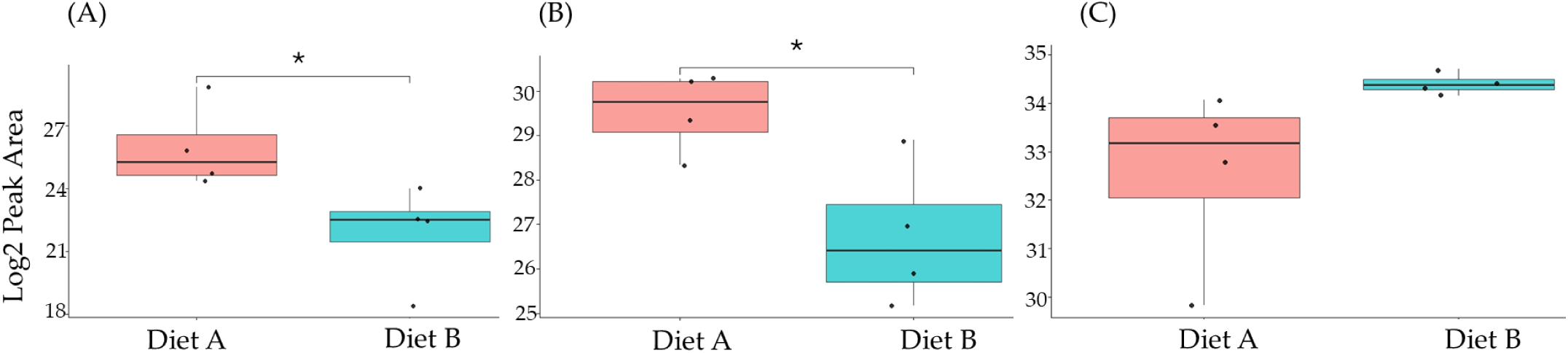
PRM validation of two proteins changed in abundance in response to dietary composition in liver tissue in Barramundi. (A) = Lysozyme g, (B) = C1q domain-containing protein, (C) = pancreatic elastase. Asterisks indicate a statistically significant difference between diet A and diet B, according to a Student t-test (* = p-value ≤ 0.05, ** = p-value ≤ 0.01, *** = p-value ≤ 0.001).

The proteins that were selected for confirmatory PRM analysis included Guanine nucleotide binding protein (G protein), ATP synthase subunit beta (EC 7.1.2.2), and ADP ribosylation factor like GTPase 3b that were all decreased significantly in abundance in the brain of fish fed on diet B (figure 13). In the liver, Lysozyme g and C1q domain-containing protein were decreased significantly in fish fed on diet B, while pancreatic elastase protein was increased in abundance (figure 14). In the intestine, Apolipoprotein E was increased significantly in abundance in the fish fed on diet B, while non-specific serine/threonine protein kinase and phosphatidylinositol-4,5-bisphosphate 3-kinase were significantly decreased (figure 15).

**Figure 15.**
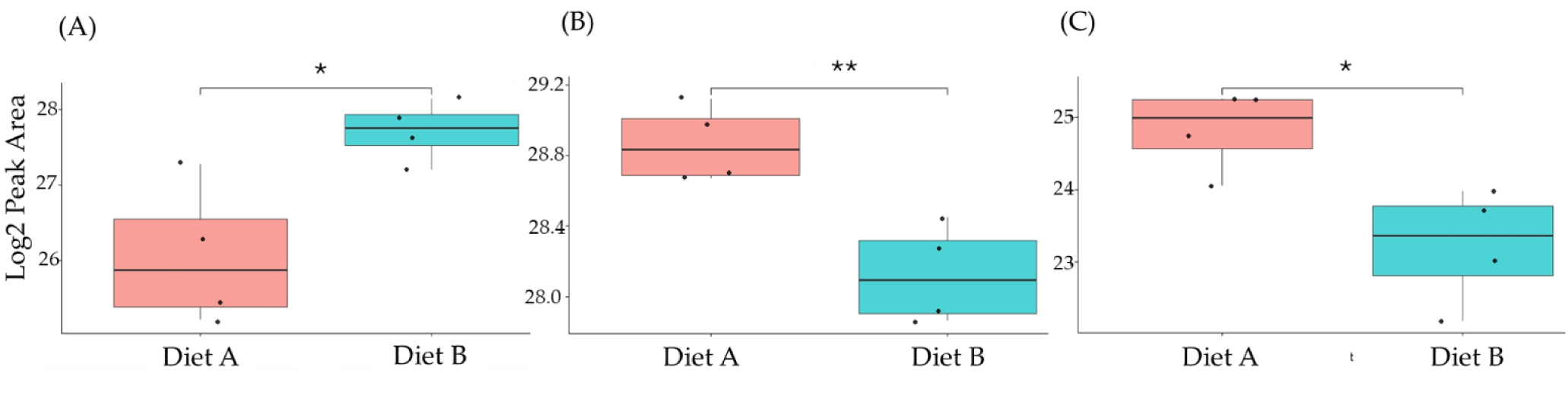
PRM validation of three proteins changed in abundance in response to dietary composition in intestine tissue in Barramundi. (A) = Apolipoprotein Ea, (B) = non-specific serine/threonine protein kinase, (C) = phosphatidylinositol-4,5-bisphosphate 3-kinase. Asterisks indicate a statistically significant difference between diet A and diet B, according to a Student t-test (* = p-value ≤ 0.05, ** = p-value ≤ 0.01, *** = p- value ≤ 0.001).

### 3.8 Iron analysis

Since the proteomic results in all three tissues demonstrated the upregulation of the ferroptosis signalling pathway, total iron analysis was performed to validate the quantitative proteomic results by confirming the accumulation of iron in the selected tissues including brain, liver, and intestine (full details in Supplementary data S8). It has been shown that iron overload disrupts iron homeostasis, induces oxidative responses, promotes ferroptosis and compromises the functional competence of chondrocytes [93].

#### 3.8.1 Total Iron analysis in brain tissue

Iron is a crucial micronutrient in fish physiology, especially for brain function [94]. Iron is a vital functional component of many proteins and enzymes in organisms. It is actively involved in a series of physiological processes including erythropoiesis, oxygen delivery, energy metabolism, and immune response [95]. The ferroptosis pathway was upregulated in the brain of the fish fed on diet B, as discussed in section 3.4.3. As shown in figure 16, the concentration of iron measured in the brain of the fish fed on diet B was higher on average (B= 0.605 ± 0. 073, A= 0.514 ± 0.196 µg/g) although the difference was not statistically significant (Supplementary data S9).

**Figure 16.**
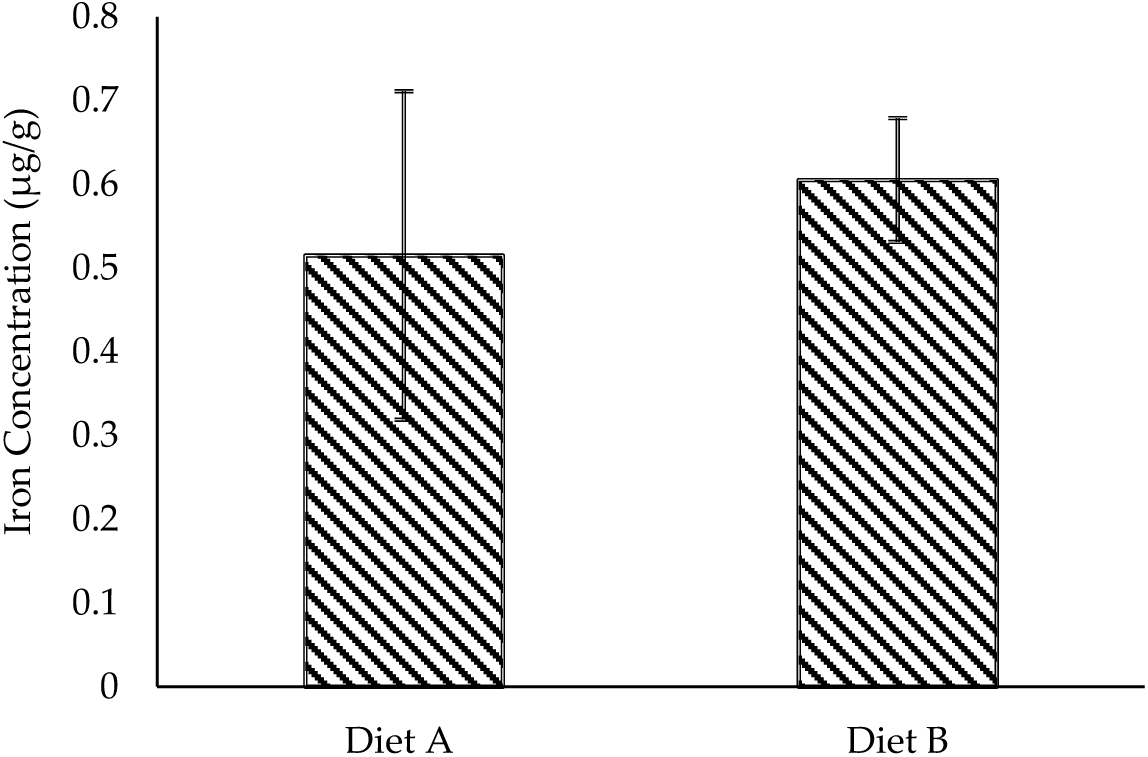
Total iron concentration (µg/g) brain tissue of the barramundi fed on diet A and B. Error bars represent the standard deviation (n=4).

However, elevated brain iron levels must be interpreted with caution as even small changes can be important. While sufficient iron is necessary for optimal brain development and performance, excessive iron accumulation can lead to oxidative responses due to the Fenton reaction, producing free radicals that damage neural tissue [96]. In this study, it is important to note that the growth parameter differences between two diets were not significant and the FCR was significantly lower for diet B. This suggests that changes in the expression of the proteins in the fish tissues are a physiological response to different nutrient availability, not necessarily a stress response.

#### 3.8.2 Total Iron analysis in liver tissue

As shown in figure 17 the concentration of iron in the liver of the barramundi fed on diet B was 123.86 ± 79.14 µg/g while for diet A it was lower at 64.96 ± 20.10 µg/g. However, due to the variability between biological replicate analysis, this result is not statistically significant.

**Figure 17.**
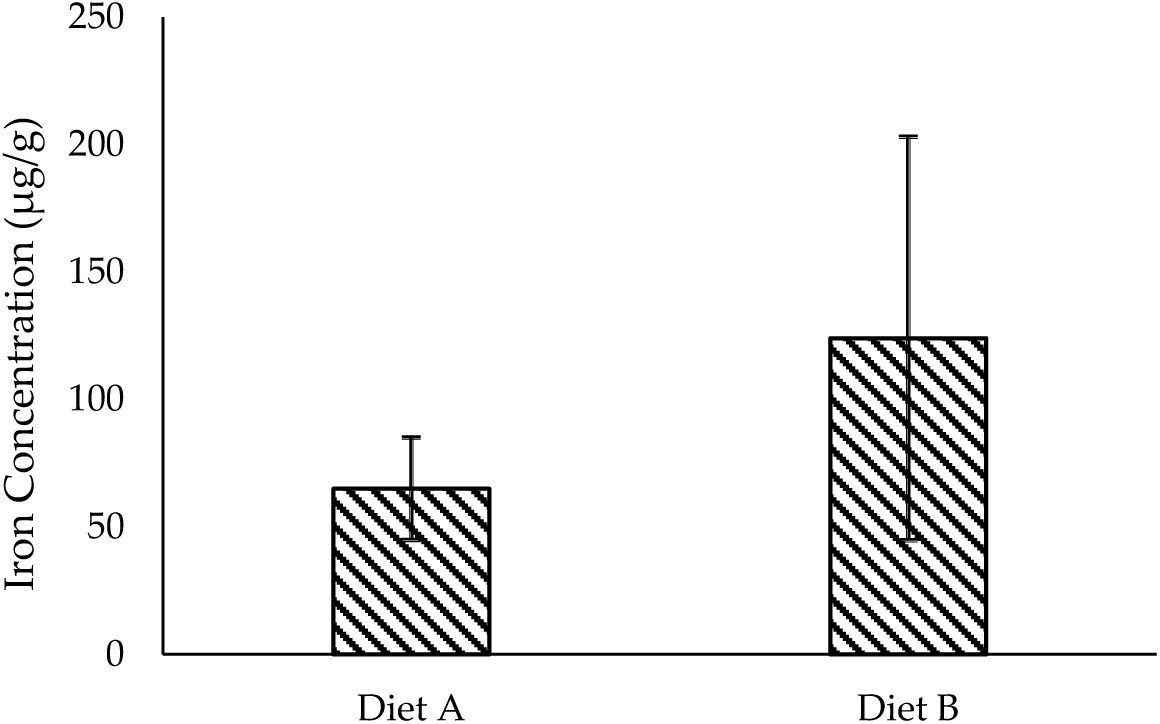
Total iron concentration (µg/g) liver tissue of the barramundi fed on diet A and B. Bars represent the standard deviation (n=4).

#### 3.8.3 Total Iron analysis in intestine tissue

As shown in Figure 18, the iron concentration in the intestinal tissue of barramundi was noticeably higher in fish fed on diet B (0.803 ± 0.227 µg/g) compared to those fed on diet A (0.425 ± 0.132 µg/g), and this difference was statistically significant. This finding suggests that while iron may be present in the intestinal lumen of fish fed on diet B, its absorption or subsequent transport into circulation might be less efficient than in fish fed on diet A. In contrast, the lower intestinal iron levels in fish fed on diet A may reflect more efficient uptake and systemic transport, potentially supported by components in the diet containing more land animal proteins. Several factors are known to affect Fe absorption, including the amount and chemical form of Fe, age and Fe status of the animal, physiological conditions of the gastrointestinal tract (e.g., pH) and other dietary components (e.g., phytic acid, ascorbic acid, citrate) [97]. In fish, ferric iron (Fe³⁺) is released from food in the acidic environment of the stomach and binds to mucin, which may help keep it soluble in the small intestine. Although the exact mechanism of iron absorption in marine fish remains unclear, it is believed that mucus secretion plays a key role in maintaining iron solubility. However, metal–mucus complexes, dietary reducing agents like ascorbic acid, and other gut conditions may help enhance iron solubility and absorption [94].

**Figure 18.**
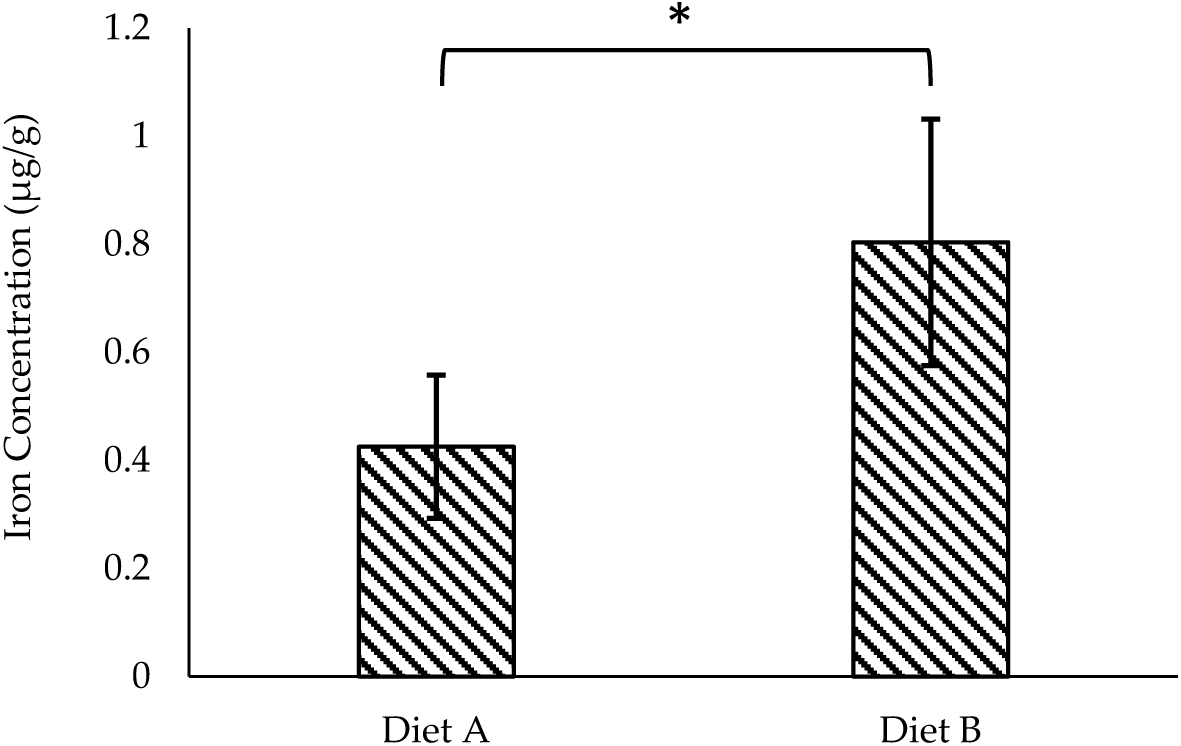
Total iron concentration (µg/g) in intestine tissue of the barramundi fed on diet A and B. Bars represent the standard deviation (n=4). Asterisk (*) indicates a statistically significant difference between groups as determined by Student’s t-test (p < 0.05).

## 4. Conclusions

Aquaculture faces ongoing pressure to identify alternative ingredients that support more sustainable feed development. As the industry shifts toward using alternative protein sources to reduce reliance on fishmeal, it becomes increasingly important to understand how these changes may affect fish health and overall well-being [98]. This study presents the first detailed quantitative proteomic analysis of barramundi (*Lates calcarifer*) using a DIA-MS-based approach. It aimed to understand how two commercial diets with the same crude protein and fat content, but different ingredient compositions, affect different tissues of the fish at a molecular level. By examining the brain, liver, and intestine, we found that even small differences in dietary protein sources can lead to noticeable changes in proteins related to metabolism, nutrient processing, and immune function. These results are aligned with a previous study performed on the rainbow trout on the transcriptomic level, in which GO and pathway analysis of the corresponding genes revealed that nutrition affects pathways of neuroendocrine peptides in the brain, and in the liver, pathways mediating intermediary metabolism, xenobiotic metabolism, proteolysis, and cytoskeletal regulation of cell cycle [99].

In the brain, we observed changes in energy production and oxidative response pathways, glutathione metabolism and ferroptosis. Notably, fish on diet B exhibited slightly lower feed intake and proteomic evidence of adaptive physiological response. Key appetite-regulating signals were also altered. For example, components of the apelin signaling pathway were downregulated in fish fed on diet B. Apelin peptides are known to act as appetite stimulants in vertebrates; in mammals, apelin increases food intake and improves energy balance [100], and recent experiments suggest a similar appetite-promoting role in fish [100].

In the liver, the diet influenced digestive enzymes and RNA-processing proteins, such as elastase II, chymotrypsin, and DDX5. We also observed activation of antioxidant and ferroptosis-related pathways, which is consistent with an adaptive response. It is well known that diet quality and ingredient type can influence hepatic metabolism in fish. For example, completely replacing fishmeal with plant proteins has been shown to alter hepatic proteomic profiles and induce metabolic pathways in other species [99]. One interesting observation was the activation of proteins linked to ferroptosis, which is a type of cell death triggered by too much iron and oxidative response [101].

The intestine showed increased activity in amino acid biosynthesis, glutathione metabolism, and mitochondrial energy production. Proteins linked to serine and arginine production were more abundant, possibly helping the fish to adapt to different nutrient composition. At the same time, proteins involved in fat transport and general metabolism were reduced, suggesting a shift in how nutrients were being absorbed and processed. Some stress-related proteins, like NACHT domain-containing proteins and ApoE, were also elevated, reflecting normal physiological adjustments to varying nutrient availability rather than a sign of immune activation.

The iron analysis performed for all three tissues demonstrated a higher level of iron for the fish fed on diet B. However, it was only statistically significant in the intestine.

Altogether, this research highlights how what fish eat, despite having comparable feed intake, influences their biology at the protein level. When the immune system is activated, it demands specific nutrients, which can lead to competition between what is needed for basic maintenance, immune function, and growth-related protein deposition. This is especially important in aquaculture, where nutrition plays a key role in fish health. As feed is the largest cost in aquaculture production, optimizing diet formulation is essential for both animal welfare and economic sustainability [98]. In this study, the growth performance between fish fed on two diets were not significant, and the FCR was significantly lower for fish fed on diet B. Lower FCR in aquaculture is regarded as high feed efficiency [102]. Although fish fed on diet B showed upregulation of ferroptosis and higher iron content in the intestine, it seems upregulation of glutathione metabolism as an antioxidant mediated the ferroptosis [103], and changes in the protein level is are adaptive response of the fish to compositional differences to the diet, in order to maintain optimal growth performance. It is also possible that some of these observed changes reflect regulation at the proteoform level, as proteins can exist in multiple modified or processed forms that influence their biological function [104].

## 5. Limitations

Although this study provides valuable insight into the tissue-specific proteomic responses of *Lates calcarifer* to two commercial diets, several limitations should be acknowledged. The proteomic analysis focused on total protein abundance, and post-translational modifications or specific proteoforms were not directly assessed. These modifications may play critical roles in metabolic regulation and warrant further investigation using top-down or targeted proteomic approaches. While this study identified molecular signatures associated with dietary composition, complementary measurements such as metabolomics or transcriptomics could provide additional layers of evidence linking molecular changes to physiological outcomes. Finally, the feeding trial was conducted under controlled laboratory conditions, which may not fully represent environmental variations in commercial aquaculture systems. Future work should evaluate how these molecular responses translate under production-scale settings.

## Supplementary Materials

The following supporting information can be found in the supplementary data: Supplementary data S1 - Proteins identified in each tissue; Supplementary data S2 - differentially expressed proteins of brain tissue, Diet A1-4, Diet B1-4; Supplementary data S3 - functional analysis of differentially expressed proteins of brain using String database; Supplementary data S4 - differentially expressed proteins of liver tissue, Diet A1-4, Diet B1-4; Supplementary data S5 - functional analysis of differentially expressed proteins of liver using String database; Supplementary data S6 - differentially expressed proteins of intestine tissue, Diet A1-4, Diet B1-4; Supplementary data S7 - functional analysis of differentially expressed proteins of intestine using String database; Supplementary data S8 - list of isolated peptides and corresponding quantification from PRM analysis; Supplementary data S9 - iron concentration measured by ICP-MS in brain, liver and intestine.

## Author Contributions

Conceptualization, IP, PDS, MMS and PAH; methodology IP, PDS, MMS and PAH; software, MMS, ZA; Mass Spectrometry, AA, ZA; resources, PDS, PAH; data curation, MMS, ZA; writing - original draft preparation, MMS; writing - review and editing, MMS, PAH, IP and PDS; visualization, MMS; supervision, PAH, IP. All authors have read and agreed to the published version of the manuscript.

## Data Availability Statement

The mass spectrometry proteomics data have been deposited to the ProteomeXchange Consortium via the jPOST [105] repository with the dataset identifier PXD069810 (https://repository.jpostdb.org/entry/JPST004137(.

## Acknowledgments

The feeding trial was conducted at Port Stephens Fisheries Institute and commercial diets were provided by Skretting Australia. Aspects of this research were conducted at the Australian Proteome Analysis Facility. The authors wish to thank Macquarie University for providing Ph.D. scholarship funding, and Wayne O’Connor for continued encouragement and support.

