## Supplementary Tables 1 to 9 for "Evaluating the impact of two different diets on the protein profile expression of the brain, liver, and intestine of barramundi": Figures and Tables.docx

**3.** Results and Discussion

**Table 1.** Summary of number of proteins identified in tissue replicates

| **Tissue Type** | **Brain** | **Liver** | **Intestine** |
| --- | --- | --- | --- |
| Diet A Rep1 | 3501 | 2986 | 4295 |
| Diet A Rep2 | 3390 | 2868 | 4351 |
| Diet A Rep3 | 3270 | 3342 | 4413 |
| Diet A Rep4 | 3299 | 3142 | 4482 |
| Diet A-Average | 3365 | 3084 | 4385 |
| **Tissue Type** | **Brain** | **Liver** | **Intestine** |
| Diet B Rep1 | 2941 | 2896 | 4493 |
| Diet B Rep2 | 3356 | 3069 | 4545 |
| Diet B Rep3 | 3195 | 2051 | 4484 |
| Diet B Rep4 | 3154 | 2664 | 4471 |
| Diet B-Average | 3161 | 2670 | 4498 |

**Table 2.** Comparison of brain samples within and between diets

| Diet | Total | Increased | Decreased | Total changed | Percentage |
| --- | --- | --- | --- | --- | --- |
| A vs A | 3889 | 101 | 133 | 234 | 6.01% |
| B vs B | 3817 | 31 | 61 | 92 | 2.41% |
| A vs B | 3901 | 205 | 302 | 507 | 12.99% |

**Table 3.** Comparison of liver samples within and between diets

| Diet | Total | Increased | Decreased | Total changed | Percentage |
| --- | --- | --- | --- | --- | --- |
| A vs A | 3649 | 144 | 143 | 287 | 7.86% |
| B vs B | 3530 | 94 | 110 | 204 | 5.77% |
| A vs B | 3660 | 235 | 231 | 466 | 12.73% |

**Table 4.** Comparison of intestine samples within and between diets

| Diet | Total | Increased | Decreased | Total changed | Percentage |
| --- | --- | --- | --- | --- | --- |
| A vs A | 5030 | 48 | 63 | 111 | 2.20% |
| B vs B | 5003 | 135 | 126 | 261 | 5.21% |
| A vs B | 5025 | 485 | 349 | 834 | 16.59% |

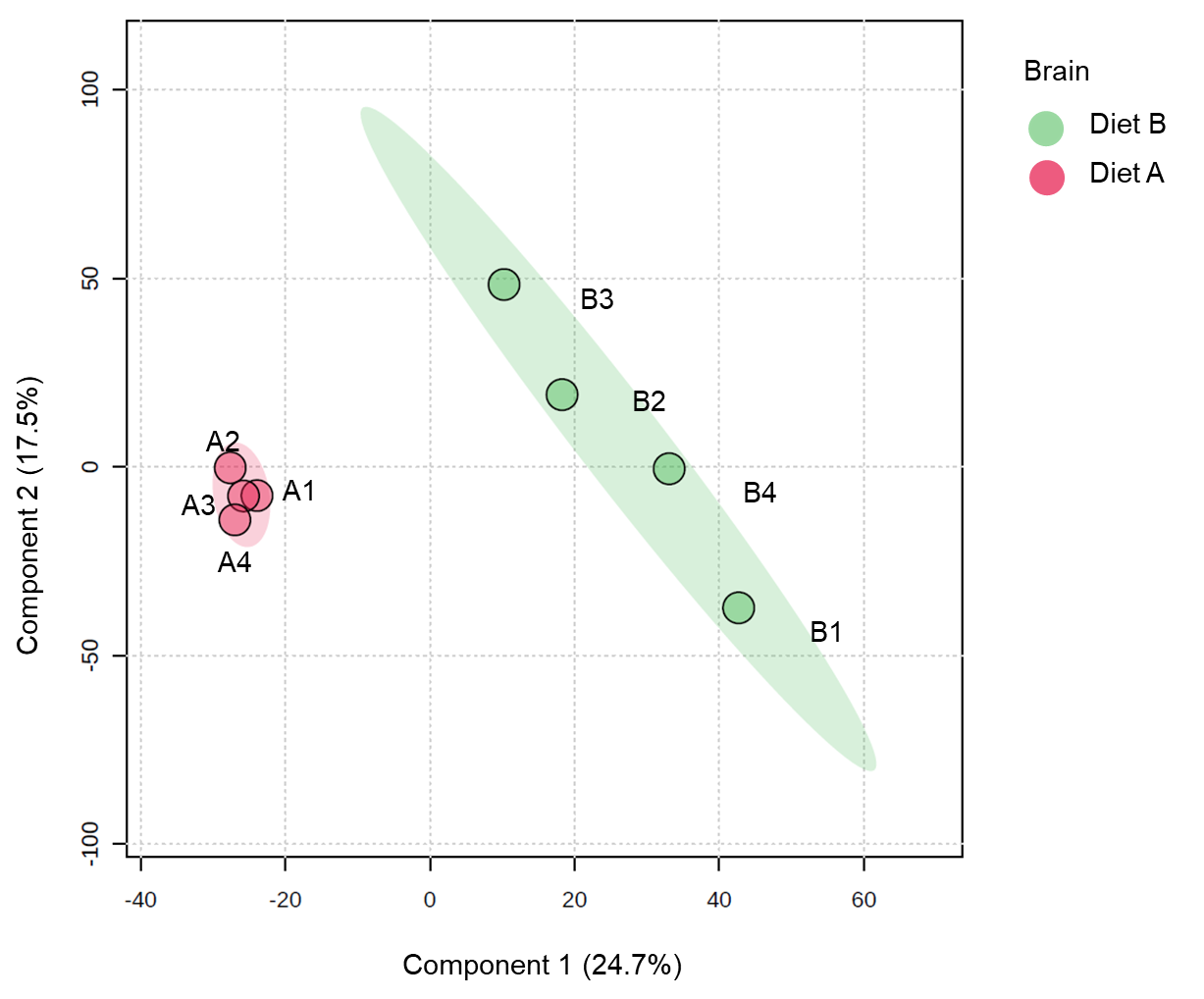

**Figure 1.** Partial least squares discriminant analysis (PLS-DA) of brain of barramundi fed on diet A and diet B displayed at 95% confidence intervals highlights the clustering of the sample based on their response to different diets (n= 4).

B vs A (Brain)

-Log 10 P-value

| Log 2 Fold-change | "Not changed" |
| --- | --- |
| 2.20503 | 4.91322 |
| -2.64252 | 4.38062 |
| -1.89382 | 4.16592 |
| 1.39156 | 4.03256 |
| 1.70227 | 3.56838 |
| -1.26746 | 3.53058 |
| 1.43594 | 3.46604 |
| -2.11572 | 3.41652 |
| -2.52041 | 3.40619 |
| -1.1673 | 3.38125 |
| -1.42878 | 3.28505 |
| -2.04981 | 3.21483 |
| -1.29433 | 3.10814 |
| -0.962251 | 3.09891 |
| 1.3623 | 3.0893 |
| -1.41808 | 3.08771 |
| -1.30645 | 3.07321 |
| 1.05894 | 3.06544 |
| -1.24264 | 3.04483 |
| -5.55929 | 3.04418 |
| -1.87613 | 3.03717 |
| -2.57304 | 3.0226 |
| -3.29542 | 3.02225 |
| -1.38179 | 2.95723 |
| -1.95355 | 2.92465 |
| 0.982493 | 2.92314 |
| -1.45132 | 2.90229 |
| 1.77686 | 2.89634 |
| -3.34314 | 2.88554 |
| 1.2788 | 2.88017 |
| 1.45991 | 2.86475 |
| -2.35864 | 2.86458 |
| 1.00253 | 2.86429 |
| -1.12674 | 2.85616 |
| 1.8243 | 2.82093 |
| -1.11968 | 2.81009 |
| 1.19334 | 2.8089 |
| -1.83094 | 2.8004 |
| -1.01326 | 2.79898 |
| -1.35997 | 2.7987 |
| -1.11942 | 2.78953 |
| -1.18714 | 2.78662 |
| -1.92221 | 2.7836 |
| -1.25636 | 2.78033 |
| -3.6193 | 2.7699 |
| 1.77737 | 2.76476 |
| -1.29375 | 2.73399 |
| -1.56814 | 2.68212 |
| -2.09127 | 2.65164 |
| -1.0608 | 2.64617 |
| 1.0505 | 2.63753 |
| -1.41549 | 2.62334 |
| -2.25967 | 2.61464 |
| 1.76197 | 2.55642 |
| 0.888988 | 2.55382 |
| 1.20904 | 2.55167 |
| -1.77491 | 2.50478 |
| -1.13363 | 2.50445 |
| 1.63766 | 2.49995 |
| -1.70221 | 2.47909 |
| -1.27716 | 2.47656 |
| -2.5785 | 2.47581 |
| 0.96389 | 2.47395 |
| -1.60662 | 2.45586 |
| 1.07932 | 2.44564 |
| -1.48636 | 2.43586 |
| -2.25313 | 2.43233 |
| 1.40446 | 2.4323 |
| -1.57383 | 2.42173 |
| 0.864278 | 2.41913 |
| -1.6468 | 2.41742 |
| 1.63283 | 2.41679 |
| -2.04393 | 2.41182 |
| 1.66863 | 2.41146 |
| 0.831712 | 2.40668 |
| -2.71638 | 2.40641 |
| 1.01515 | 2.39231 |
| 0.746154 | 2.37786 |
| -2.40432 | 2.37717 |
| -1.80739 | 2.37404 |
| -2.44419 | 2.36773 |
| 0.790384 | 2.36327 |
| -0.724336 | 2.36004 |
| -1.18057 | 2.35885 |
| -1.4293 | 2.35713 |
| -1.78831 | 2.33943 |
| 0.972747 | 2.33923 |
| -0.944106 | 2.33111 |
| -0.869037 | 2.33092 |
| 1.86432 | 2.33081 |
| -1.15204 | 2.32859 |
| -3.46679 | 2.32739 |
| 1.28404 | 2.32082 |
| -1.12664 | 2.30144 |
| -0.907436 | 2.29908 |
| -1.20619 | 2.29443 |
| -1.97847 | 2.28606 |
| 1.47383 | 2.27558 |
| -1.07205 | 2.27304 |
| -0.923871 | 2.26534 |
| -2.02923 | 2.26361 |
| -0.936083 | 2.25207 |
| 1.60154 | 2.25005 |
| -1.30111 | 2.24929 |
| -1.22335 | 2.24106 |
| -1.7088 | 2.23746 |
| -2.27358 | 2.23066 |
| -1.77545 | 2.22774 |
| 1.12633 | 2.22274 |
| -2.3653 | 2.22124 |
| 0.946312 | 2.20802 |
| -1.44527 | 2.19647 |
| -1.16553 | 2.19491 |
| 1.07232 | 2.19409 |
| -0.783906 | 2.19396 |
| -2.14763 | 2.19177 |
| -0.662151 | 2.18397 |
| -1.80796 | 2.18077 |
| -1.09977 | 2.17515 |
| 0.888034 | 2.16119 |
| 0.758851 | 2.1562 |
| -1.65608 | 2.15146 |
| 0.631249 | 2.14013 |
| 1.03244 | 2.13572 |
| 1.54272 | 2.13005 |
| -1.16056 | 2.12413 |
| -0.852725 | 2.12138 |
| 0.593532 | 2.11966 |
| -0.923047 | 2.1175 |
| -1.27132 | 2.11501 |
| -1.2153 | 2.11152 |
| -1.23107 | 2.11015 |
| 1.47738 | 2.10706 |
| 1.06395 | 2.10194 |
| 1.09743 | 2.09894 |
| -1.05333 | 2.09688 |
| -1.00016 | 2.09607 |
| -0.964795 | 2.09173 |
| -1.4303 | 2.08694 |
| -2.21858 | 2.08575 |
| 1.67148 | 2.07767 |
| -1.24471 | 2.0736 |
| -1.37636 | 2.06626 |
| -2.16313 | 2.06131 |
| -0.72532 | 2.05624 |
| 0.614044 | 2.04721 |
| 0.719494 | 2.04696 |
| 0.886891 | 2.04664 |
| 0.970521 | 2.04548 |
| 0.8012 | 2.03934 |
| -1.5966 | 2.03635 |
| 1.31068 | 2.03338 |
| -0.640077 | 2.03098 |
| -1.07218 | 2.03085 |
| 1.24208 | 2.03058 |
| -1.2333 | 2.02573 |
| 1.0492 | 2.02089 |
| -0.887949 | 2.02035 |
| 0.904064 | 2.01991 |
| -1.17557 | 2.01785 |
| 0.974779 | 1.99784 |
| -1.03076 | 1.99124 |
| -1.19474 | 1.98917 |
| 0.708915 | 1.98636 |
| -1.12733 | 1.97349 |
| 0.624038 | 1.97272 |
| 0.711658 | 1.97187 |
| 0.880493 | 1.97075 |
| -1.12085 | 1.97035 |
| -0.879859 | 1.96607 |
| 0.767406 | 1.94769 |
| 0.709204 | 1.94371 |
| -1.4796 | 1.93551 |
| -3.58503 | 1.93526 |
| -1.25225 | 1.92997 |
| 1.14999 | 1.9283 |
| -0.79988 | 1.92332 |
| -0.97713 | 1.91728 |
| -0.626091 | 1.91167 |
| -1.09254 | 1.91143 |
| 0.957635 | 1.90967 |
| -1.01933 | 1.90401 |
| 0.798921 | 1.90208 |
| -1.0879 | 1.90098 |
| 1.22189 | 1.89939 |
| -1.20704 | 1.8888 |
| -0.949031 | 1.88837 |
| -1.13672 | 1.88665 |
| -1.16224 | 1.88626 |
| 0.659374 | 1.88446 |
| -0.655017 | 1.88227 |
| 1.05255 | 1.87826 |
| 1.35295 | 1.87668 |
| -1.11918 | 1.86998 |
| 0.612115 | 1.86996 |
| -0.727114 | 1.86861 |
| -1.26699 | 1.86776 |
| 0.61722 | 1.85537 |
| 1.5731 | 1.8544 |
| -0.77603 | 1.85378 |

| Log 2 Fold-change | "Increased" |
| --- | --- |
| 0.406829 | 1.30561 |
| 0.473641 | 1.44065 |
| 0.497627 | 1.43079 |
| 0.502271 | 1.5813 |
| 0.511714 | 1.56337 |
| 0.511954 | 1.44708 |
| 0.517284 | 1.30946 |
| 0.521655 | 1.57482 |
| 0.528063 | 1.39144 |
| 0.534879 | 1.63124 |
| 0.541452 | 1.4287 |
| 0.543655 | 1.32077 |
| 0.544934 | 1.54339 |
| 0.547398 | 1.30254 |
| 0.551176 | 1.36063 |
| 0.552442 | 1.68566 |
| 0.55284 | 1.46383 |
| 0.557344 | 1.77277 |
| 0.561886 | 1.62417 |
| 0.571444 | 1.77838 |
| 0.573752 | 1.40018 |
| 0.573914 | 1.37518 |
| 0.577373 | 1.7163 |
| 0.578847 | 1.54379 |
| 0.581165 | 1.68882 |
| 0.582912 | 1.34599 |
| 0.586321 | 1.37496 |
| 0.586473 | 1.42315 |
| 0.589727 | 1.66202 |
| 0.591232 | 1.62469 |
| 0.591635 | 1.39172 |
| 0.593532 | 2.11966 |
| 0.599534 | 1.39291 |
| 0.604164 | 1.62954 |
| 0.610829 | 1.40695 |
| 0.611663 | 1.46604 |
| 0.612115 | 1.86996 |
| 0.614044 | 2.04721 |
| 0.614167 | 1.4562 |
| 0.615895 | 1.52022 |
| 0.61722 | 1.85537 |
| 0.624038 | 1.97272 |
| 0.625476 | 1.81577 |
| 0.628604 | 1.64973 |
| 0.631249 | 2.14013 |
| 0.631662 | 1.39407 |
| 0.634471 | 1.59453 |
| 0.635897 | 1.81163 |
| 0.637021 | 1.502 |
| 0.638486 | 1.37876 |
| 0.64246 | 1.43234 |
| 0.650833 | 1.74063 |
| 0.656841 | 1.48353 |
| 0.65861 | 1.43467 |
| 0.659374 | 1.88446 |
| 0.660533 | 1.45614 |
| 0.661618 | 1.81747 |
| 0.665699 | 1.30501 |
| 0.665971 | 1.45023 |
| 0.667471 | 1.38952 |
| 0.673662 | 1.47441 |
| 0.674371 | 1.55692 |
| 0.68337 | 1.63604 |
| 0.684852 | 1.34029 |
| 0.691093 | 1.51454 |
| 0.692233 | 1.71786 |
| 0.693985 | 1.65126 |
| 0.696065 | 1.3844 |
| 0.697298 | 1.4771 |
| 0.699629 | 1.77256 |
| 0.701736 | 1.57318 |
| 0.703664 | 1.30632 |
| 0.705506 | 1.40296 |
| 0.705728 | 1.7684 |
| 0.708915 | 1.98636 |
| 0.709204 | 1.94371 |
| 0.710171 | 1.69337 |
| 0.711658 | 1.97187 |
| 0.714257 | 1.38539 |
| 0.717203 | 1.40845 |
| 0.719494 | 2.04696 |
| 0.73326 | 1.34519 |
| 0.737622 | 1.66513 |
| 0.744832 | 1.67515 |
| 0.746154 | 2.37786 |
| 0.748974 | 1.53354 |
| 0.757019 | 1.64956 |
| 0.758851 | 2.1562 |
| 0.760391 | 1.45947 |
| 0.764607 | 1.68091 |
| 0.767406 | 1.94769 |
| 0.781379 | 1.59127 |
| 0.785543 | 1.45438 |
| 0.790384 | 2.36327 |
| 0.794998 | 1.60327 |
| 0.798921 | 1.90208 |
| 0.8012 | 2.03934 |
| 0.802095 | 1.39332 |
| 0.808764 | 1.58946 |
| 0.810268 | 1.83701 |
| 0.81907 | 1.4894 |
| 0.821656 | 1.65572 |
| 0.822187 | 1.40071 |
| 0.831712 | 2.40668 |
| 0.84417 | 1.50075 |
| 0.864278 | 2.41913 |
| 0.864315 | 1.55869 |
| 0.869091 | 1.46532 |
| 0.873098 | 1.49574 |
| 0.880493 | 1.97075 |
| 0.886891 | 2.04664 |
| 0.888034 | 2.16119 |
| 0.888988 | 2.55382 |
| 0.895513 | 1.55187 |
| 0.904064 | 2.01991 |
| 0.905735 | 1.3077 |
| 0.906512 | 1.43506 |
| 0.908792 | 1.62749 |
| 0.916484 | 1.6227 |
| 0.929432 | 1.54445 |
| 0.940525 | 1.54838 |
| 0.946312 | 2.20802 |
| 0.949034 | 1.49839 |
| 0.954451 | 1.31903 |
| 0.957635 | 1.90967 |
| 0.96389 | 2.47395 |
| 0.967011 | 1.7457 |
| 0.967549 | 1.49643 |
| 0.970521 | 2.04548 |
| 0.972747 | 2.33923 |
| 0.974015 | 1.42905 |
| 0.974779 | 1.99784 |
| 0.982493 | 2.92314 |
| 0.99279 | 1.56743 |
| 0.993613 | 1.82173 |
| 1.00253 | 2.86429 |
| 1.00854 | 1.55165 |
| 1.01515 | 2.39231 |
| 1.03244 | 2.13572 |
| 1.0492 | 2.02089 |
| 1.0505 | 2.63753 |
| 1.05255 | 1.87826 |
| 1.05894 | 3.06544 |
| 1.06395 | 2.10194 |
| 1.07232 | 2.19409 |
| 1.0741 | 1.41916 |
| 1.07932 | 2.44564 |
| 1.08361 | 1.82879 |
| 1.09689 | 1.35266 |
| 1.09743 | 2.09894 |
| 1.12633 | 2.22274 |
| 1.12806 | 1.53194 |
| 1.14493 | 1.33908 |
| 1.14999 | 1.9283 |
| 1.18113 | 1.65934 |
| 1.18461 | 1.57208 |
| 1.19334 | 2.8089 |
| 1.20904 | 2.55167 |
| 1.21101 | 1.55167 |
| 1.22189 | 1.89939 |
| 1.23275 | 1.85176 |
| 1.23614 | 1.7272 |
| 1.24208 | 2.03058 |
| 1.24334 | 1.45576 |
| 1.24514 | 1.47507 |
| 1.25673 | 1.46272 |
| 1.27061 | 1.42544 |
| 1.2788 | 2.88017 |
| 1.28404 | 2.32082 |
| 1.30467 | 1.55515 |
| 1.31068 | 2.03338 |
| 1.32629 | 1.85292 |
| 1.32943 | 1.75262 |
| 1.33274 | 1.51569 |
| 1.35295 | 1.87668 |
| 1.35555 | 1.77433 |
| 1.3623 | 3.0893 |
| 1.36877 | 1.51558 |
| 1.39156 | 4.03256 |
| 1.40446 | 2.4323 |
| 1.42845 | 1.65054 |
| 1.43594 | 3.46604 |
| 1.43815 | 1.42433 |
| 1.45991 | 2.86475 |
| 1.47383 | 2.27558 |
| 1.47738 | 2.10706 |
| 1.47847 | 1.435 |
| 1.50863 | 1.58999 |
| 1.54272 | 2.13005 |
| 1.5731 | 1.8544 |
| 1.59717 | 1.85296 |
| 1.60154 | 2.25005 |
| 1.63283 | 2.41679 |
| 1.63766 | 2.49995 |
| 1.66863 | 2.41146 |
| 1.67148 | 2.07767 |
| 1.70227 | 3.56838 |
| 1.73657 | 1.85187 |
| 1.76197 | 2.55642 |
| 1.77686 | 2.89634 |

| Log 2 Fold-change | "Decreased" |
| --- | --- |
| -5.55929 | 3.04418 |
| -3.80447 | 1.65116 |
| -3.6193 | 2.7699 |
| -3.58503 | 1.93526 |
| -3.46679 | 2.32739 |
| -3.38103 | 1.63275 |
| -3.34314 | 2.88554 |
| -3.29542 | 3.02225 |
| -2.71638 | 2.40641 |
| -2.64252 | 4.38062 |
| -2.5785 | 2.47581 |
| -2.57304 | 3.0226 |
| -2.52041 | 3.40619 |
| -2.44419 | 2.36773 |
| -2.40432 | 2.37717 |
| -2.3653 | 2.22124 |
| -2.35864 | 2.86458 |
| -2.31625 | 1.67791 |
| -2.27358 | 2.23066 |
| -2.25967 | 2.61464 |
| -2.25313 | 2.43233 |
| -2.24484 | 1.77119 |
| -2.21998 | 1.30672 |
| -2.21858 | 2.08575 |
| -2.16313 | 2.06131 |
| -2.14763 | 2.19177 |
| -2.14306 | 1.48467 |
| -2.11572 | 3.41652 |
| -2.09127 | 2.65164 |
| -2.04981 | 3.21483 |
| -2.04393 | 2.41182 |
| -2.02923 | 2.26361 |
| -2.00999 | 1.43651 |
| -1.97847 | 2.28606 |
| -1.95355 | 2.92465 |
| -1.92221 | 2.7836 |
| -1.89382 | 4.16592 |
| -1.87613 | 3.03717 |
| -1.84183 | 1.766 |
| -1.83094 | 2.8004 |
| -1.80796 | 2.18077 |
| -1.80739 | 2.37404 |
| -1.78831 | 2.33943 |
| -1.77545 | 2.22774 |
| -1.77491 | 2.50478 |
| -1.7088 | 2.23746 |
| -1.70221 | 2.47909 |
| -1.70028 | 1.59868 |
| -1.65608 | 2.15146 |
| -1.6468 | 2.41742 |
| -1.61988 | 1.58255 |
| -1.60662 | 2.45586 |
| -1.5966 | 2.03635 |
| -1.57393 | 1.30299 |
| -1.57383 | 2.42173 |
| -1.56814 | 2.68212 |
| -1.56482 | 1.62853 |
| -1.55505 | 1.59058 |
| -1.5213 | 1.3142 |
| -1.51855 | 1.65503 |
| -1.48636 | 2.43586 |
| -1.4796 | 1.93551 |
| -1.47667 | 1.38033 |
| -1.45204 | 1.30306 |
| -1.45132 | 2.90229 |
| -1.44527 | 2.19647 |
| -1.4303 | 2.08694 |
| -1.4293 | 2.35713 |
| -1.42878 | 3.28505 |
| -1.41808 | 3.08771 |
| -1.41549 | 2.62334 |
| -1.41111 | 1.82132 |
| -1.40608 | 1.38116 |
| -1.38179 | 2.95723 |
| -1.37636 | 2.06626 |
| -1.37584 | 1.51458 |
| -1.35997 | 2.7987 |
| -1.31347 | 1.60658 |
| -1.30907 | 1.76323 |
| -1.30752 | 1.40637 |
| -1.30645 | 3.07321 |
| -1.30111 | 2.24929 |
| -1.30098 | 1.5945 |
| -1.29433 | 3.10814 |
| -1.29375 | 2.73399 |
| -1.27716 | 2.47656 |
| -1.2726 | 1.70657 |
| -1.27132 | 2.11501 |
| -1.26746 | 3.53058 |
| -1.26717 | 1.45214 |
| -1.26699 | 1.86776 |
| -1.26195 | 1.64872 |
| -1.25636 | 2.78033 |
| -1.25225 | 1.92997 |
| -1.24471 | 2.0736 |
| -1.24264 | 3.04483 |
| -1.24229 | 1.63758 |
| -1.23756 | 1.44466 |
| -1.2333 | 2.02573 |
| -1.23107 | 2.11015 |
| -1.22889 | 1.84294 |
| -1.22395 | 1.60985 |
| -1.22335 | 2.24106 |
| -1.2153 | 2.11152 |
| -1.21305 | 1.3524 |
| -1.20704 | 1.8888 |
| -1.20619 | 2.29443 |
| -1.20537 | 1.77698 |
| -1.20112 | 1.81719 |
| -1.19474 | 1.98917 |
| -1.18714 | 2.78662 |
| -1.18057 | 2.35885 |
| -1.17762 | 1.79727 |
| -1.17557 | 2.01785 |
| -1.17255 | 1.65743 |
| -1.17185 | 1.4233 |
| -1.1673 | 3.38125 |
| -1.16564 | 1.49271 |
| -1.16553 | 2.19491 |
| -1.16547 | 1.683 |
| -1.16224 | 1.88626 |
| -1.16056 | 2.12413 |
| -1.15357 | 1.4651 |
| -1.15204 | 2.32859 |
| -1.13672 | 1.88665 |
| -1.13363 | 2.50445 |
| -1.12733 | 1.97349 |
| -1.12674 | 2.85616 |
| -1.12664 | 2.30144 |
| -1.12085 | 1.97035 |
| -1.11968 | 2.81009 |
| -1.11942 | 2.78953 |
| -1.11918 | 1.86998 |
| -1.10553 | 1.56457 |
| -1.09977 | 2.17515 |
| -1.09254 | 1.91143 |
| -1.09167 | 1.41869 |
| -1.0879 | 1.90098 |
| -1.08624 | 1.59051 |
| -1.07971 | 1.57193 |
| -1.07218 | 2.03085 |
| -1.07205 | 2.27304 |
| -1.0608 | 2.64617 |
| -1.05799 | 1.41105 |
| -1.05333 | 2.09688 |
| -1.03076 | 1.99124 |
| -1.01933 | 1.90401 |
| -1.01583 | 1.62287 |
| -1.01326 | 2.79898 |
| -1.01217 | 1.4479 |
| -1.00421 | 1.32256 |
| -1.00016 | 2.09607 |
| -0.995055 | 1.55328 |
| -0.988849 | 1.37302 |
| -0.979265 | 1.75242 |
| -0.979161 | 1.82518 |
| -0.979061 | 1.35198 |
| -0.97713 | 1.91728 |
| -0.976633 | 1.74755 |
| -0.970053 | 1.56914 |
| -0.964795 | 2.09173 |
| -0.964032 | 1.37771 |
| -0.962251 | 3.09891 |
| -0.958655 | 1.50532 |
| -0.953069 | 1.7599 |
| -0.950849 | 1.36557 |
| -0.9498 | 1.39218 |
| -0.949031 | 1.88837 |
| -0.944106 | 2.33111 |
| -0.941664 | 1.62878 |
| -0.941578 | 1.37859 |
| -0.941018 | 1.31586 |
| -0.93817 | 1.31779 |
| -0.936083 | 2.25207 |
| -0.933438 | 1.8024 |
| -0.925428 | 1.69542 |
| -0.923871 | 2.26534 |
| -0.923047 | 2.1175 |
| -0.921829 | 1.51111 |
| -0.919485 | 1.41398 |
| -0.918708 | 1.39741 |
| -0.917884 | 1.57333 |
| -0.908659 | 1.44412 |
| -0.907436 | 2.29908 |
| -0.895875 | 1.32691 |
| -0.894788 | 1.6837 |
| -0.888363 | 1.33431 |
| -0.887949 | 2.02035 |
| -0.879859 | 1.96607 |
| -0.875301 | 1.42971 |
| -0.869037 | 2.33092 |
| -0.867007 | 1.49911 |
| -0.863181 | 1.33918 |
| -0.861624 | 1.36999 |
| -0.860229 | 1.62314 |
| -0.854513 | 1.30426 |
| -0.852725 | 2.12138 |
| -0.849566 | 1.4921 |
| -0.847942 | 1.47984 |
| -0.847787 | 1.53523 |

**Figure 2.** Volcano plot illustrates significantly differentially abundant proteins in brain. The −log10 (*P value*) is plotted against the log_2_ (fold change: diet B/diet A).

**Table 5.** Top ten DEPs in brain tissue

| **Row** | **Protein ID** | **Protein Name** | **Fold change*** |
| --- | --- | --- | --- |
| **Increased abundance in diet B** | | | |
| 1 | A0A4W6EX76 | Transporter | 7.49 |
| 2 | A0A4W6ER29 | Ferritin | 4.61 |
| 3 | A0A4W6D6I8 | Peptidyl-prolyl cis-trans isomerase (PPIase) 5.2.1.8) | 3.64 |
| 4 | A0A4W6D1Z6 | Adhesion G protein-coupled receptor L2b, tandem duplicate 1 | 3.54 |
| 5 | A0A4W6EE98 | Myosin X, like 1 | 3.42 |
| 6 | A0A4W6ETZ5 | Protein tyrosine phosphatase receptor type Nb | 3.42 |
| 7 | A0A4W6FSI9 | LIM zinc-binding domain-containing protein | 3.39 |
| 8 | A0A4W6EUJ3 | 3-oxoacyl-(acyl-carrier-protein) synthase | 3.33 |
| 9 | A0A4W6FB21 | NADH dehydrogenase (ubiquinone) iron-sulfur protein 5 | 3.25 |
| 10 | A0A4W6E7W2 | Sodium channel subunit beta-2 isoform X1 | 3.18 |
| **Decreased abundance in diet B** | | | |
| 1 | A0A4W6DCM7 | Guanine nucleotide-binding protein subunit alpha | 0.02 |
| 2 | A0A4W6EGV8 | 1-phosphatidylinositol 4,5-bisphosphate phosphodiesterase | 0.07 |
| 3 | A0A4W6DLB9 | Cell adhesion molecule 1a | 0.08 |
| 4 | A0A4W6CDB0 | Spermine synthase | 0.08 |
| 5 | A0A4W6DXP0 | Guanine nucleotide binding protein (G protein) | 0.09 |
| 6 | A0A4W6CNV2 | ATP synthase subunit beta (EC 7.1.2.2) | 0.09 |
| 7 | A0A4W6BVV5 | Small ribosomal subunit protein uS2 | 0.09 |
| 8 | A0A4W6DBS0 | ADP ribosylation factor like GTPase 3b | 0.10 |
| 9 | A0A4W6CBE8 | Solute carrier family 17 member 6 | 0.15 |
| 10 | A0A4W6DBM8 | Uncharacterized protein LOC108891021 | 0.16 |

*Values > 1.3 indicate more abundant in diet B, values < 0.76 indicate less abundant in diet B

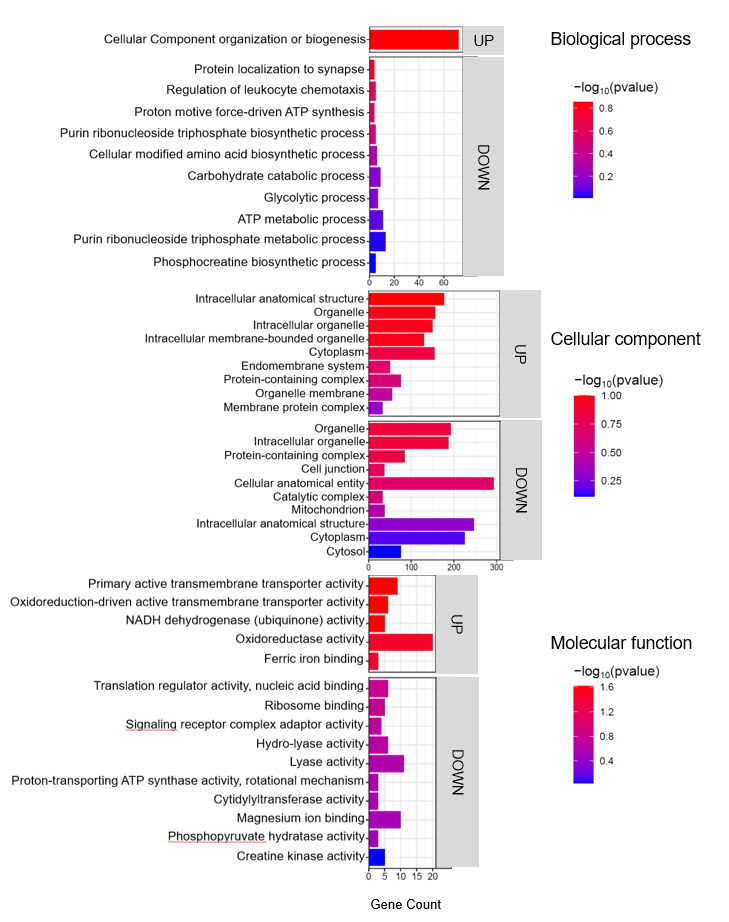
**Figure 3.** Top enriched GO terms in the DEPs of brain tissue from barramundi. A higher –log₁₀(p-value) (red) indicates greater statistical significance, while lower –log₁₀(p-value) (blue) values indicate less significant enrichment. Up indicates increased abundance in the brain of fish fed on diet B, while Down indicates decreased abundance in the brain of fish fed on diet B.

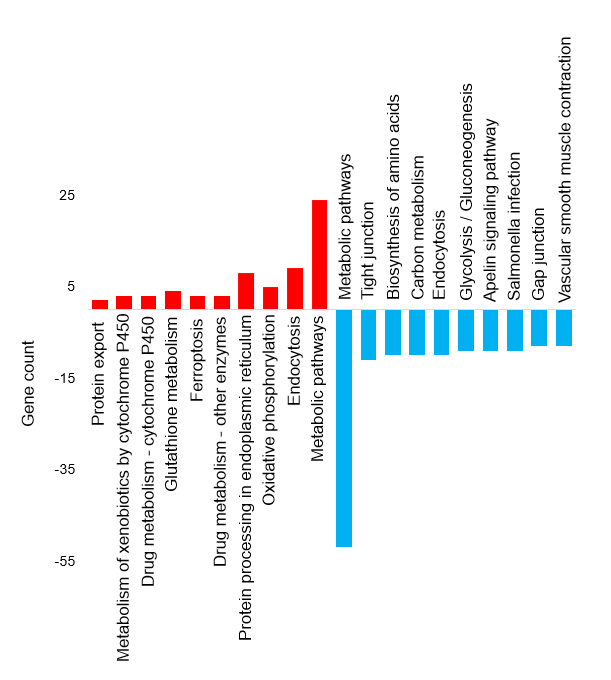

**Figure 4.** Top KEGG pathways of the differentially abundant proteins in brain tissue

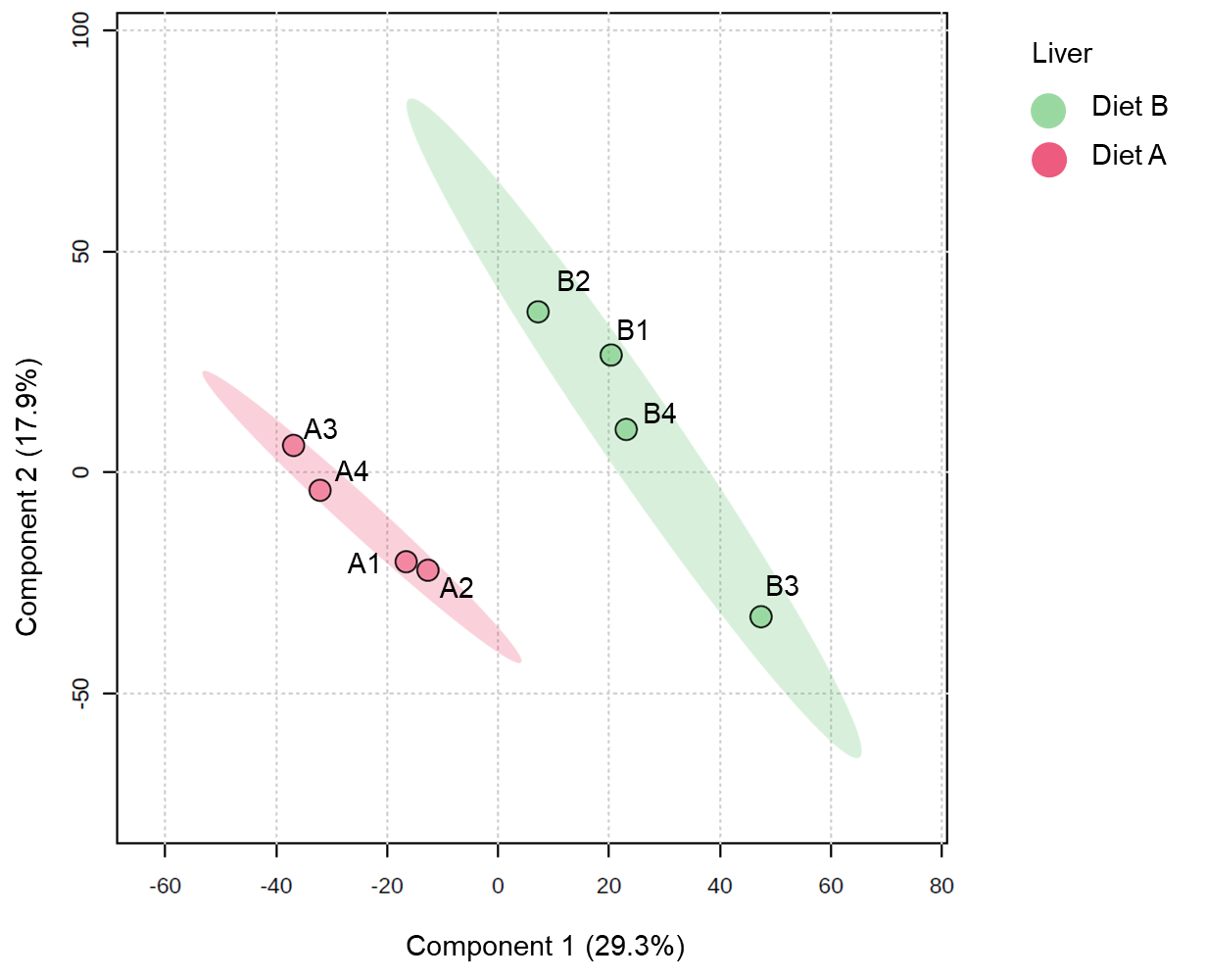

**Figure 5**. Partial least squares discriminant analysis (PLS-DA) of liver of barramundi fed on diet A and diet B displayed at 95% confidence intervals highlights the clustering of the sample based on their response to different diets (n= 4).

B vs A (Liver)

-Log10(P-value)

| Log Fold-change | "Not changed" |
| --- | --- |
| 1.96186 | 3.67951 |
| -1.29682 | 3.65595 |
| -1.56298 | 3.5303 |
| -2.50936 | 3.51374 |
| 1.04187 | 3.25242 |
| -1.39745 | 3.24723 |
| -1.43941 | 3.22678 |
| -1.51839 | 3.14382 |
| 0.934693 | 3.03216 |
| -1.95524 | 3.02271 |
| -1.20115 | 3.01942 |
| -1.15706 | 3.00771 |
| -1.96519 | 2.944 |
| 1.52622 | 2.89784 |
| -0.844665 | 2.89261 |
| -1.09046 | 2.81425 |
| -1.42696 | 2.81388 |
| -1.25355 | 2.8084 |
| -1.74775 | 2.80054 |
| -1.18188 | 2.78932 |
| 0.871807 | 2.73628 |
| 0.771188 | 2.71692 |
| -1.07709 | 2.7137 |
| 1.4356 | 2.71073 |
| 1.01653 | 2.69214 |
| -1.33026 | 2.66969 |
| -0.902637 | 2.66902 |
| 0.726807 | 2.65941 |
| -1.47446 | 2.63826 |
| 0.868626 | 2.56556 |
| -0.938828 | 2.55506 |
| 1.2566 | 2.54349 |
| -1.73427 | 2.47505 |
| 2.28327 | 2.46339 |
| -2.73102 | 2.46075 |
| -2.09944 | 2.42545 |
| 1.63634 | 2.41786 |
| -1.35735 | 2.40322 |
| -0.848675 | 2.40132 |
| -1.52678 | 2.4013 |
| -0.860722 | 2.37562 |
| -2.52606 | 2.36994 |
| 0.767139 | 2.36924 |
| -1.91416 | 2.36304 |
| -0.848869 | 2.36105 |
| -0.742687 | 2.35647 |
| 1.68136 | 2.35567 |
| -0.72794 | 2.3486 |
| -0.940078 | 2.3447 |
| 1.34298 | 2.33924 |
| 1.21237 | 2.33801 |
| -0.838459 | 2.33229 |
| 1.5578 | 2.31612 |
| -1.30645 | 2.30787 |
| 0.981319 | 2.29328 |
| 1.25824 | 2.288 |
| -1.72511 | 2.27827 |
| 1.11551 | 2.27492 |
| -1.45577 | 2.26981 |
| -1.20578 | 2.26759 |
| 0.85091 | 2.25754 |
| -0.835089 | 2.25559 |
| -1.42273 | 2.23418 |
| -1.17721 | 2.22252 |
| -1.38505 | 2.21819 |
| 1.09024 | 2.21772 |
| -0.581727 | 2.21733 |
| -0.69307 | 2.21161 |
| 1.93186 | 2.20213 |
| -1.0871 | 2.19655 |
| 0.779627 | 2.191 |
| -0.984241 | 2.18959 |
| -1.02024 | 2.17711 |
| 1.58722 | 2.17483 |
| -0.953844 | 2.17412 |
| -0.600469 | 2.17139 |
| 1.20213 | 2.1686 |
| 1.2517 | 2.16548 |
| -1.0484 | 2.15384 |
| 0.720555 | 2.15137 |
| -0.869805 | 2.15058 |
| -1.08903 | 2.14934 |
| -0.992723 | 2.14426 |
| -1.45598 | 2.13928 |
| 1.09377 | 2.13558 |
| 1.29986 | 2.1276 |
| -0.815124 | 2.12594 |
| 1.54488 | 2.10605 |
| -1.09155 | 2.10506 |
| -1.00846 | 2.09462 |
| -0.680187 | 2.08788 |
| 0.83169 | 2.08623 |
| -1.03953 | 2.08557 |
| 0.668358 | 2.07585 |
| 0.636646 | 2.0706 |
| -1.26615 | 2.06952 |
| 0.940469 | 2.06818 |
| -1.09669 | 2.06723 |
| -1.49737 | 2.06165 |
| 1.14853 | 2.06143 |
| -0.863805 | 2.05589 |
| -1.26373 | 2.05402 |
| -1.08152 | 2.05132 |
| -2.21284 | 2.0405 |
| 0.751485 | 2.03239 |
| -0.688614 | 2.03127 |
| -0.873695 | 2.02894 |
| 0.672348 | 2.02869 |
| -0.648041 | 2.02793 |
| 1.17258 | 2.02526 |
| -0.694017 | 2.01851 |
| -1.08087 | 2.01374 |
| 0.860691 | 2.00939 |
| -1.45199 | 2.00629 |
| 0.657162 | 1.99938 |
| 0.702955 | 1.99604 |
| 0.727953 | 1.99579 |
| -1.87662 | 1.99459 |
| 0.76465 | 1.99115 |
| 0.730846 | 1.98786 |
| 0.890483 | 1.98576 |
| -1.32046 | 1.98512 |
| 0.831481 | 1.98408 |
| -1.17154 | 1.98172 |
| 0.660436 | 1.98089 |
| -1.55607 | 1.96819 |
| -1.78727 | 1.9676 |
| -0.859607 | 1.9664 |
| -0.559678 | 1.96123 |
| 1.83379 | 1.95447 |
| 0.904575 | 1.95093 |
| 0.822632 | 1.94591 |
| -0.689034 | 1.94254 |
| -0.770649 | 1.94213 |
| 0.78076 | 1.93078 |
| 1.63305 | 1.92602 |
| 0.79735 | 1.922 |
| -0.725929 | 1.92185 |
| 0.786141 | 1.92075 |
| 1.18145 | 1.91865 |
| 0.871763 | 1.91633 |
| 1.46912 | 1.91428 |
| -0.660714 | 1.91033 |
| 0.985259 | 1.90814 |
| -1.11818 | 1.90712 |
| -0.73317 | 1.89468 |
| 0.742771 | 1.89459 |
| -0.760335 | 1.88599 |
| -1.2508 | 1.88195 |
| 0.965943 | 1.8698 |
| -0.733661 | 1.86859 |
| 1.54463 | 1.86701 |
| -2.3683 | 1.86399 |
| 0.892928 | 1.85734 |
| -0.679711 | 1.85545 |
| 1.10924 | 1.85034 |
| -0.911595 | 1.84629 |
| 1.61512 | 1.84571 |
| 0.581426 | 1.84404 |
| -0.945211 | 1.84338 |
| -0.710339 | 1.84283 |
| 0.621348 | 1.84209 |
| 1.25989 | 1.83835 |
| -0.764484 | 1.83455 |
| 0.675478 | 1.83074 |
| -1.55872 | 1.82722 |
| 1.2128 | 1.8231 |
| 1.04046 | 1.81891 |
| -0.861839 | 1.81755 |
| 1.05537 | 1.81006 |
| 1.31719 | 1.80488 |
| 1.14515 | 1.7954 |
| 1.1786 | 1.79305 |
| 0.939649 | 1.78901 |
| 0.899521 | 1.78673 |
| -0.593794 | 1.78404 |
| 1.14417 | 1.78321 |
| 1.59623 | 1.78244 |
| 0.929346 | 1.78132 |
| -0.873028 | 1.77547 |
| -1.55083 | 1.76753 |
| 1.3177 | 1.76718 |
| 1.1906 | 1.76668 |
| -0.770265 | 1.76457 |
| -0.664233 | 1.76391 |
| 0.874645 | 1.76295 |
| -0.792405 | 1.75458 |
| 1.27159 | 1.75311 |
| -0.908737 | 1.74809 |
| -1.26952 | 1.74732 |
| -1.53875 | 1.74247 |
| 1.14967 | 1.73805 |
| -0.613914 | 1.73766 |
| 0.758526 | 1.73711 |
| -0.642235 | 1.73201 |
| 0.757601 | 1.72928 |
| -1.36908 | 1.7292 |
| -0.678524 | 1.72264 |
| 1.76963 | 1.72057 |
| -0.669527 | 1.71711 |

| Log Fold-change | "Increased" |
| --- | --- |
| 0.394024 | 1.46154 |
| 0.398153 | 1.40306 |
| 0.414253 | 1.43915 |
| 0.421872 | 1.30325 |
| 0.429556 | 1.32502 |
| 0.435242 | 1.36979 |
| 0.446544 | 1.37335 |
| 0.448772 | 1.51351 |
| 0.44898 | 1.43454 |
| 0.462186 | 1.37762 |
| 0.462897 | 1.42143 |
| 0.467168 | 1.55679 |
| 0.478614 | 1.61708 |
| 0.479663 | 1.33424 |
| 0.486665 | 1.32811 |
| 0.507932 | 1.41354 |
| 0.513276 | 1.31028 |
| 0.51852 | 1.32288 |
| 0.521717 | 1.40155 |
| 0.538138 | 1.55416 |
| 0.542105 | 1.33318 |
| 0.543572 | 1.43263 |
| 0.564436 | 1.39359 |
| 0.576401 | 1.51084 |
| 0.580138 | 1.52044 |
| 0.581426 | 1.84404 |
| 0.591351 | 1.4296 |
| 0.597475 | 1.45315 |
| 0.601327 | 1.43487 |
| 0.601346 | 1.56441 |
| 0.601459 | 1.6992 |
| 0.602399 | 1.35089 |
| 0.604362 | 1.46741 |
| 0.607281 | 1.50818 |
| 0.611639 | 1.57285 |
| 0.616851 | 1.36055 |
| 0.621348 | 1.84209 |
| 0.621541 | 1.57078 |
| 0.628378 | 1.39464 |
| 0.634249 | 1.45066 |
| 0.636646 | 2.0706 |
| 0.652151 | 1.45827 |
| 0.653897 | 1.62749 |
| 0.65531 | 1.43053 |
| 0.655511 | 1.44759 |
| 0.655945 | 1.52396 |
| 0.657162 | 1.99938 |
| 0.660436 | 1.98089 |
| 0.668358 | 2.07585 |
| 0.672348 | 2.02869 |
| 0.675478 | 1.83074 |
| 0.675966 | 1.51092 |
| 0.676414 | 1.62803 |
| 0.677863 | 1.46533 |
| 0.687417 | 1.68917 |
| 0.68853 | 1.38652 |
| 0.690612 | 1.48723 |
| 0.690802 | 1.53534 |
| 0.691206 | 1.58552 |
| 0.695555 | 1.47054 |
| 0.695774 | 1.68391 |
| 0.698712 | 1.48711 |
| 0.702955 | 1.99604 |
| 0.705034 | 1.71605 |
| 0.705234 | 1.54526 |
| 0.705245 | 1.30363 |
| 0.714608 | 1.67141 |
| 0.71636 | 1.38803 |
| 0.719746 | 1.43824 |
| 0.720555 | 2.15137 |
| 0.724913 | 1.57902 |
| 0.726807 | 2.65941 |
| 0.727953 | 1.99579 |
| 0.730846 | 1.98786 |
| 0.735283 | 1.33983 |
| 0.742771 | 1.89459 |
| 0.751485 | 2.03239 |
| 0.756217 | 1.68516 |
| 0.757601 | 1.72928 |
| 0.758526 | 1.73711 |
| 0.761316 | 1.51661 |
| 0.76465 | 1.99115 |
| 0.767139 | 2.36924 |
| 0.769118 | 1.64932 |
| 0.771188 | 2.71692 |
| 0.771974 | 1.54507 |
| 0.776732 | 1.51304 |
| 0.779627 | 2.191 |
| 0.78076 | 1.93078 |
| 0.783668 | 1.53485 |
| 0.786141 | 1.92075 |
| 0.786556 | 1.67975 |
| 0.79735 | 1.922 |
| 0.797745 | 1.66358 |
| 0.805653 | 1.42996 |
| 0.808671 | 1.45588 |
| 0.80983 | 1.62918 |
| 0.811325 | 1.54052 |
| 0.815792 | 1.36808 |
| 0.822632 | 1.94591 |
| 0.823101 | 1.30305 |
| 0.831481 | 1.98408 |
| 0.83169 | 2.08623 |
| 0.833082 | 1.36608 |
| 0.837024 | 1.44585 |
| 0.846622 | 1.40156 |
| 0.847368 | 1.33907 |
| 0.847978 | 1.38084 |
| 0.848487 | 1.50448 |
| 0.85091 | 2.25754 |
| 0.858156 | 1.58703 |
| 0.859902 | 1.3896 |
| 0.860691 | 2.00939 |
| 0.861678 | 1.69406 |
| 0.864094 | 1.44309 |
| 0.864912 | 1.59767 |
| 0.868626 | 2.56556 |
| 0.871763 | 1.91633 |
| 0.871807 | 2.73628 |
| 0.873587 | 1.55225 |
| 0.874645 | 1.76295 |
| 0.87546 | 1.47151 |
| 0.885074 | 1.69721 |
| 0.888526 | 1.71473 |
| 0.890483 | 1.98576 |
| 0.892928 | 1.85734 |
| 0.899351 | 1.51169 |
| 0.899521 | 1.78673 |
| 0.904575 | 1.95093 |
| 0.905107 | 1.44295 |
| 0.910028 | 1.59507 |
| 0.923343 | 1.51603 |
| 0.923546 | 1.52315 |
| 0.924116 | 1.50314 |
| 0.929346 | 1.78132 |
| 0.929961 | 1.59378 |
| 0.934643 | 1.41078 |
| 0.934693 | 3.03216 |
| 0.937364 | 1.53393 |
| 0.939649 | 1.78901 |
| 0.940469 | 2.06818 |
| 0.965943 | 1.8698 |
| 0.968712 | 1.55878 |
| 0.981301 | 1.58295 |
| 0.981319 | 2.29328 |
| 0.981927 | 1.35118 |
| 0.985259 | 1.90814 |
| 0.996988 | 1.36346 |
| 1.00306 | 1.60757 |
| 1.00324 | 1.61618 |
| 1.01505 | 1.51861 |
| 1.01653 | 2.69214 |
| 1.01827 | 1.55802 |
| 1.0193 | 1.43082 |
| 1.04046 | 1.81891 |
| 1.04056 | 1.41222 |
| 1.04187 | 3.25242 |
| 1.05537 | 1.81006 |
| 1.07011 | 1.46033 |
| 1.09024 | 2.21772 |
| 1.09254 | 1.41152 |
| 1.09377 | 2.13558 |
| 1.1085 | 1.54984 |
| 1.10924 | 1.85034 |
| 1.11551 | 2.27492 |
| 1.11662 | 1.3733 |
| 1.11726 | 1.70436 |
| 1.12881 | 1.3486 |
| 1.13033 | 1.43607 |
| 1.13045 | 1.62725 |
| 1.13599 | 1.34827 |
| 1.14417 | 1.78321 |
| 1.14515 | 1.7954 |
| 1.14853 | 2.06143 |
| 1.14967 | 1.73805 |
| 1.15902 | 1.68708 |
| 1.16082 | 1.357 |
| 1.17258 | 2.02526 |
| 1.1786 | 1.79305 |
| 1.18145 | 1.91865 |
| 1.19022 | 1.43652 |
| 1.1906 | 1.76668 |
| 1.20213 | 2.1686 |
| 1.21237 | 2.33801 |
| 1.2128 | 1.8231 |
| 1.22004 | 1.55643 |
| 1.22416 | 1.40803 |
| 1.22841 | 1.42653 |
| 1.2517 | 2.16548 |
| 1.2566 | 2.54349 |
| 1.25824 | 2.288 |
| 1.25989 | 1.83835 |
| 1.27159 | 1.75311 |
| 1.29219 | 1.37807 |
| 1.29986 | 2.1276 |
| 1.31719 | 1.80488 |
| 1.3177 | 1.76718 |
| 1.34298 | 2.33924 |
| 1.34378 | 1.46433 |
| 1.34393 | 1.57421 |

| Log Fold-change | "Decreased" |
| --- | --- |
| -2.73102 | 2.46075 |
| -2.52606 | 2.36994 |
| -2.50936 | 3.51374 |
| -2.3683 | 1.86399 |
| -2.21284 | 2.0405 |
| -2.09944 | 2.42545 |
| -1.96519 | 2.944 |
| -1.95524 | 3.02271 |
| -1.91416 | 2.36304 |
| -1.87662 | 1.99459 |
| -1.83499 | 1.45702 |
| -1.79973 | 1.66129 |
| -1.78727 | 1.9676 |
| -1.74775 | 2.80054 |
| -1.73427 | 2.47505 |
| -1.72511 | 2.27827 |
| -1.56298 | 3.5303 |
| -1.55872 | 1.82722 |
| -1.55607 | 1.96819 |
| -1.55424 | 1.57555 |
| -1.55083 | 1.76753 |
| -1.53875 | 1.74247 |
| -1.52678 | 2.4013 |
| -1.51839 | 3.14382 |
| -1.50929 | 1.69378 |
| -1.49737 | 2.06165 |
| -1.48468 | 1.46107 |
| -1.47533 | 1.5996 |
| -1.47446 | 2.63826 |
| -1.46298 | 1.46819 |
| -1.45598 | 2.13928 |
| -1.45577 | 2.26981 |
| -1.45199 | 2.00629 |
| -1.43941 | 3.22678 |
| -1.42696 | 2.81388 |
| -1.42273 | 2.23418 |
| -1.39745 | 3.24723 |
| -1.38505 | 2.21819 |
| -1.37677 | 1.42811 |
| -1.36908 | 1.7292 |
| -1.35735 | 2.40322 |
| -1.33026 | 2.66969 |
| -1.32498 | 1.61624 |
| -1.32046 | 1.98512 |
| -1.30645 | 2.30787 |
| -1.29682 | 3.65595 |
| -1.29252 | 1.68524 |
| -1.28232 | 1.65158 |
| -1.27261 | 1.55517 |
| -1.26952 | 1.74732 |
| -1.26615 | 2.06952 |
| -1.26373 | 2.05402 |
| -1.25355 | 2.8084 |
| -1.2508 | 1.88195 |
| -1.23429 | 1.44616 |
| -1.20873 | 1.32769 |
| -1.20578 | 2.26759 |
| -1.20115 | 3.01942 |
| -1.19892 | 1.71414 |
| -1.18188 | 2.78932 |
| -1.17721 | 2.22252 |
| -1.17154 | 1.98172 |
| -1.17028 | 1.56934 |
| -1.15706 | 3.00771 |
| -1.15501 | 1.31686 |
| -1.13146 | 1.48092 |
| -1.12192 | 1.51018 |
| -1.11925 | 1.57154 |
| -1.11818 | 1.90712 |
| -1.10908 | 1.52259 |
| -1.10767 | 1.63307 |
| -1.10755 | 1.33594 |
| -1.09916 | 1.67269 |
| -1.09669 | 2.06723 |
| -1.09381 | 1.32412 |
| -1.09155 | 2.10506 |
| -1.09046 | 2.81425 |
| -1.08903 | 2.14934 |
| -1.0871 | 2.19655 |
| -1.08152 | 2.05132 |
| -1.08087 | 2.01374 |
| -1.07709 | 2.7137 |
| -1.06538 | 1.6377 |
| -1.0537 | 1.33068 |
| -1.0484 | 2.15384 |
| -1.03953 | 2.08557 |
| -1.02024 | 2.17711 |
| -1.00846 | 2.09462 |
| -0.992723 | 2.14426 |
| -0.992643 | 1.61244 |
| -0.984241 | 2.18959 |
| -0.969977 | 1.42027 |
| -0.953844 | 2.17412 |
| -0.945211 | 1.84338 |
| -0.940078 | 2.3447 |
| -0.939081 | 1.34954 |
| -0.938828 | 2.55506 |
| -0.931908 | 1.30412 |
| -0.926283 | 1.61349 |
| -0.919353 | 1.30991 |
| -0.918237 | 1.61541 |
| -0.911595 | 1.84629 |
| -0.908737 | 1.74809 |
| -0.906349 | 1.52589 |
| -0.902637 | 2.66902 |
| -0.893005 | 1.37404 |
| -0.8909 | 1.38546 |
| -0.888802 | 1.59751 |
| -0.886422 | 1.71478 |
| -0.884737 | 1.4042 |
| -0.875642 | 1.33419 |
| -0.873695 | 2.02894 |
| -0.873028 | 1.77547 |
| -0.869805 | 2.15058 |
| -0.865716 | 1.41588 |
| -0.864532 | 1.70189 |
| -0.863805 | 2.05589 |
| -0.861839 | 1.81755 |
| -0.860722 | 2.37562 |
| -0.859607 | 1.9664 |
| -0.854559 | 1.69443 |
| -0.848869 | 2.36105 |
| -0.848675 | 2.40132 |
| -0.846532 | 1.31654 |
| -0.845834 | 1.39768 |
| -0.844665 | 2.89261 |
| -0.838459 | 2.33229 |
| -0.835747 | 1.45299 |
| -0.835089 | 2.25559 |
| -0.8333 | 1.39363 |
| -0.825459 | 1.47119 |
| -0.820346 | 1.46577 |
| -0.819565 | 1.5301 |
| -0.816228 | 1.32097 |
| -0.815124 | 2.12594 |
| -0.814626 | 1.40699 |
| -0.813404 | 1.31489 |
| -0.805296 | 1.63874 |
| -0.804733 | 1.65125 |
| -0.799913 | 1.55434 |
| -0.798024 | 1.66671 |
| -0.792405 | 1.75458 |
| -0.788946 | 1.50795 |
| -0.783603 | 1.32574 |
| -0.775993 | 1.38297 |
| -0.7725 | 1.60775 |
| -0.770649 | 1.94213 |
| -0.770265 | 1.76457 |
| -0.769727 | 1.50724 |
| -0.765816 | 1.45647 |
| -0.764484 | 1.83455 |
| -0.760335 | 1.88599 |
| -0.756281 | 1.46715 |
| -0.756244 | 1.31618 |
| -0.748388 | 1.6941 |
| -0.746059 | 1.53054 |
| -0.742687 | 2.35647 |
| -0.741053 | 1.32754 |
| -0.733661 | 1.86859 |
| -0.733597 | 1.66007 |
| -0.733421 | 1.4836 |
| -0.73317 | 1.89468 |
| -0.72794 | 2.3486 |
| -0.725929 | 1.92185 |
| -0.711397 | 1.65334 |
| -0.710339 | 1.84283 |
| -0.705278 | 1.32421 |
| -0.701531 | 1.51721 |
| -0.69767 | 1.70563 |
| -0.696556 | 1.46239 |
| -0.694017 | 2.01851 |
| -0.69307 | 2.21161 |
| -0.689034 | 1.94254 |
| -0.688614 | 2.03127 |
| -0.686104 | 1.50675 |
| -0.681358 | 1.54618 |
| -0.680187 | 2.08788 |
| -0.679711 | 1.85545 |
| -0.678524 | 1.72264 |
| -0.677864 | 1.32995 |
| -0.670369 | 1.33366 |
| -0.669527 | 1.71711 |
| -0.668906 | 1.40519 |
| -0.667889 | 1.46432 |
| -0.664233 | 1.76391 |
| -0.660714 | 1.91033 |
| -0.65677 | 1.42098 |
| -0.650882 | 1.32011 |
| -0.648041 | 2.02793 |
| -0.644519 | 1.47718 |
| -0.642235 | 1.73201 |
| -0.638349 | 1.34521 |
| -0.637744 | 1.44652 |
| -0.632734 | 1.58531 |
| -0.632051 | 1.32475 |
| -0.623821 | 1.4175 |
| -0.616231 | 1.66474 |
| -0.613914 | 1.73766 |
| -0.600469 | 2.17139 |
| -0.600173 | 1.31177 |

**Figure 6**. Volcano plot illustrates significantly differentially abundant proteins in liver. The −log_10_ (*P value*) is plotted against the log_2_ (fold change: diet B/diet A).

**Table 6.** Top ten DEPs in the Liver tissue

| **Row** | **Protein ID** | **Protein Name** | **Fold* change** |
| --- | --- | --- | --- |
| **Increased abundance in diet B** | | | |
| 1 | A0A4W6G873 | pancreatic elastase II | 7.13 |
| 2 | A0A4W6CY37 | Chymotrypsin-like | 7.04 |
| 3 | A0A4W6FQC7 | Vesicle transport protein USE1 | 6.99 |
| 4 | A0A4W6D8E6 | Lin-9 DREAM MuvB core complex component | 5.05 |
| 5 | A0A4W6BK05 | RNA helicase | 4.86 |
| 6 | A0A4W6DXB6 | Dematin actin binding protein | 4.74 |
| 7 | A0A4W6FXS8 | Myoglobin | 3.91 |
| 8 | A0A4W6CMS6 | O-acyltransferase | 3.89 |
| 9 | A0A4W6F0G8 | SPRY domain containing 4 | 3.81 |
| 10 | A0A4W6E876 | Ring finger protein 7 | 3.64 |
| **Decreased abundance in diet B** | | | |
| 11 | A0A4W6CQU6 | C1q domain-containing protein | 0.15 |
| 12 | A0A4W6FZS4 | Zinc finger CCCH-type containing 7B | 0.17 |
| 13 | Q6ITU9 | Parvalbumin | 0.19 |
| 14 | A0A4W6EPI8 | Ig-like domain-containing protein | 0.21 |
| 15 | A0A4W6BTM5 | Uncharacterized protein | 0.25 |
| 16 | A0A4W6BKR3 | RNA helicase | 0.25 |
| 17 | A0A4W6BMF0 | CUB domain-containing protein | 0.26 |
| 18 | A0A4W6EUI1 | Si:dkey-69o16.5 | 0.27 |
| 19 | A0A4W6FRU8 | Prothrombin | 0.28 |
| 20 | A8D3J6 | Lysozyme g | 0.28 |

*Values > 1.3 indicate more abundant in diet B, values < 0.76 indicate less abundant in diet B

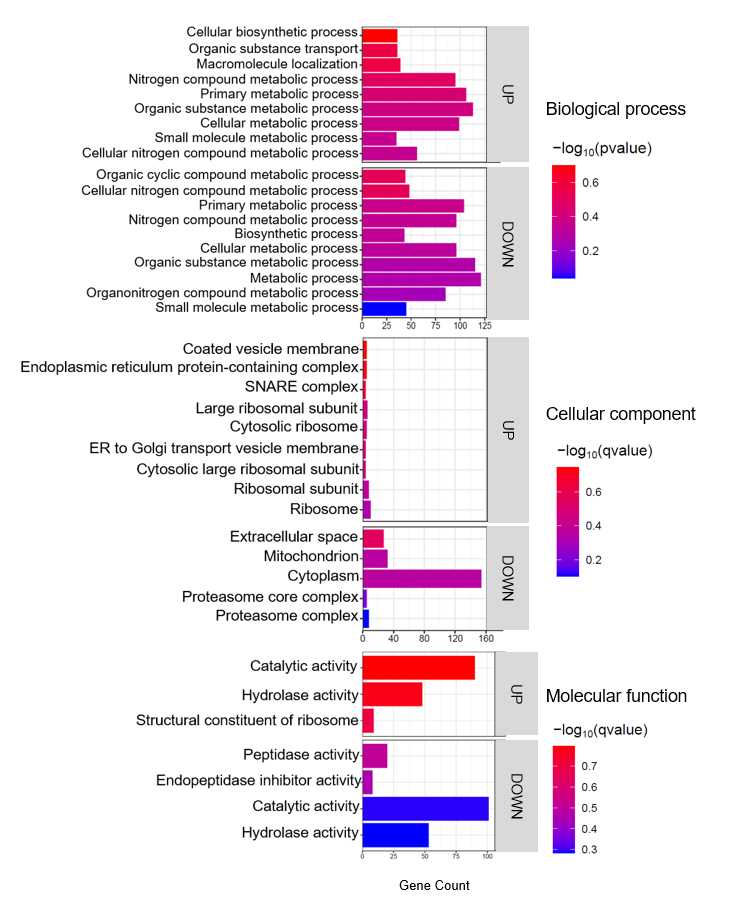

**Figure 7.** Top enriched GO terms in the DEPs of liver of barramundi. A higher –log₁₀(p-value) (red) indicates greater statistical significance, while lower –log₁₀(p-value) (blue) values indicate less significant enrichment. Up indicates increased abundance in the liver of fish fed on diet B, while Down indicates decreased abundance in the liver of fish fed on diet B.

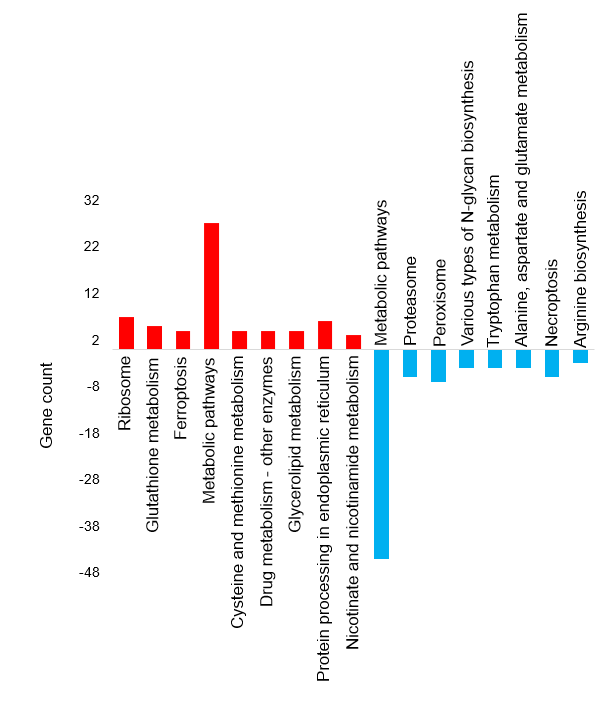

**Figure 8.** Top KEGG pathways of the differentially abundant proteins in liver tissue

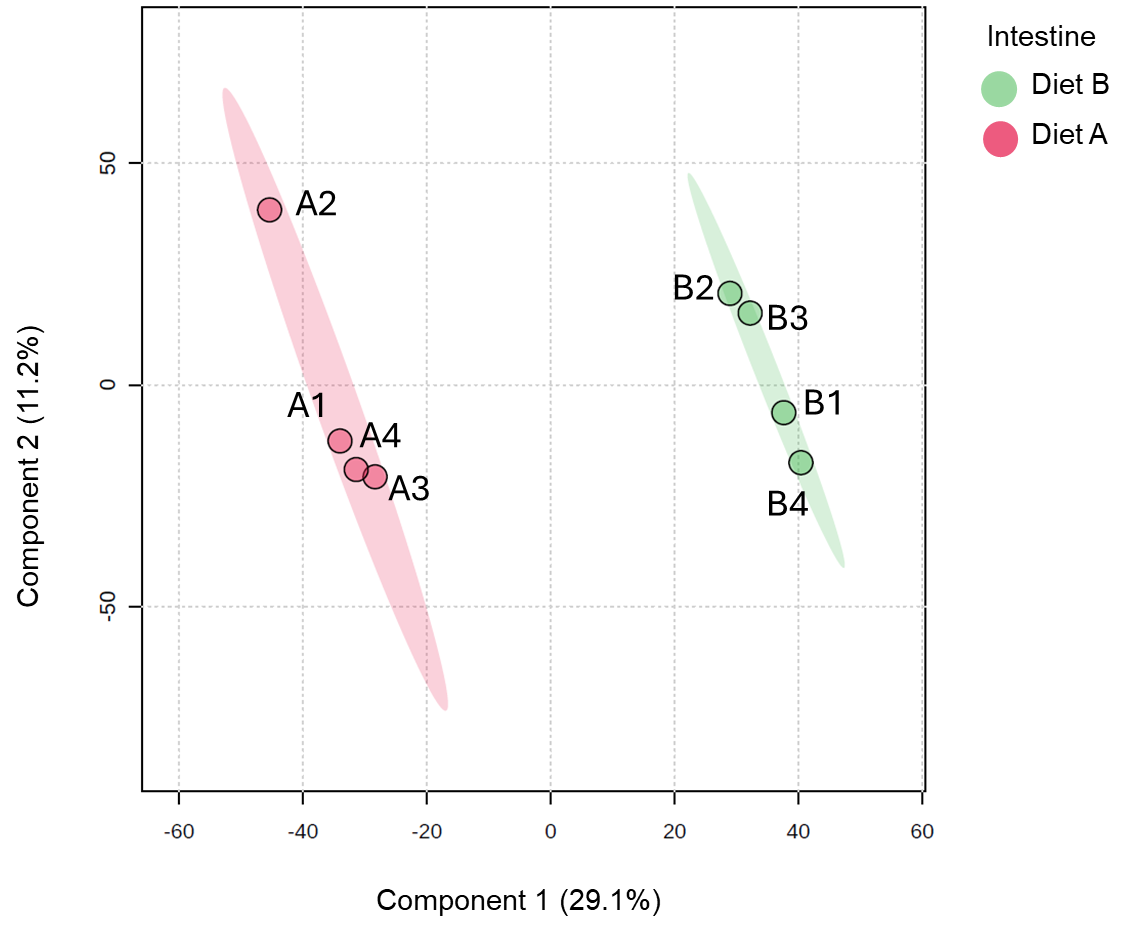

**Figure 9**. Partial least squares discriminant analysis (PLS-DA) of intestine of barramundi fed on diet A and diet B displayed at 95% confidence intervals highlights the clustering of the sample based on their response to different diets (n= 4).

B vs A (Intestine)

-Log10(P-value)

| Log Fold-change | "Not changed" |
| --- | --- |
| 1.50649 | 5.61932 |
| -3.86088 | 5.39384 |
| 1.49669 | 5.09942 |
| 1.50213 | 5.04656 |
| 2.24716 | 4.87347 |
| -1.03945 | 4.87159 |
| 1.13331 | 4.71404 |
| 1.1421 | 4.64615 |
| 1.76977 | 4.63952 |
| 1.34749 | 4.63684 |
| 2.79157 | 4.58757 |
| 1.19969 | 4.40986 |
| 0.894915 | 4.40601 |
| 1.23856 | 4.38189 |
| 1.56453 | 4.32326 |
| 1.26471 | 4.31299 |
| -2.30968 | 4.29741 |
| -1.44562 | 4.14434 |
| -1.96505 | 4.13595 |
| 0.839276 | 4.07457 |
| -0.889955 | 3.97907 |
| 1.04073 | 3.89488 |
| -1.20297 | 3.86603 |
| 0.873916 | 3.79852 |
| 3.18976 | 3.74981 |
| 1.05744 | 3.74781 |
| -0.920209 | 3.73558 |
| 0.886363 | 3.69984 |
| 0.67686 | 3.66184 |
| 0.784234 | 3.65522 |
| -1.18045 | 3.62838 |
| -0.992493 | 3.61875 |
| -1.35301 | 3.55696 |
| 1.0216 | 3.54661 |
| 1.07516 | 3.54487 |
| 0.96336 | 3.53672 |
| 1.40944 | 3.46998 |
| -0.998735 | 3.46671 |
| 1.12154 | 3.45278 |
| 0.977183 | 3.43158 |
| -0.621676 | 3.41001 |
| 1.2333 | 3.3976 |
| -1.2713 | 3.39225 |
| 1.3283 | 3.36566 |
| 0.774727 | 3.35595 |
| -1.31313 | 3.33764 |
| -0.695548 | 3.33323 |
| -1.3158 | 3.30177 |
| 0.798436 | 3.27252 |
| 0.849702 | 3.263 |
| -0.741869 | 3.24991 |
| 0.728129 | 3.23103 |
| 1.44028 | 3.20847 |
| 0.821995 | 3.19052 |
| 0.832739 | 3.18027 |
| -0.579908 | 3.1757 |
| 1.70896 | 3.17337 |
| 0.825502 | 3.16858 |
| -1.00236 | 3.15811 |
| 1.0197 | 3.132 |
| -0.577315 | 3.12516 |
| 0.527529 | 3.12169 |
| -0.837479 | 3.1193 |
| 0.677958 | 3.11662 |
| 0.617256 | 3.1111 |
| -0.758619 | 3.10776 |
| 1.20417 | 3.10178 |
| 0.757197 | 3.09535 |
| 1.02943 | 3.07634 |
| 0.684112 | 3.07614 |
| 0.694941 | 3.06623 |
| 1.35661 | 3.06262 |
| -1.33205 | 3.05255 |
| 0.678456 | 3.04271 |
| 0.637826 | 3.02912 |
| 1.19241 | 3.02717 |
| 0.792046 | 3.02503 |
| -0.924389 | 3.02281 |
| -1.67244 | 2.99531 |
| 0.741714 | 2.99422 |
| -0.64051 | 2.98696 |
| 0.653814 | 2.96028 |
| -0.568813 | 2.95816 |
| -1.36737 | 2.95722 |
| 0.82128 | 2.9571 |
| 1.52995 | 2.93991 |
| -0.751377 | 2.93889 |
| -1.28392 | 2.92582 |
| -1.36394 | 2.92368 |
| -0.751929 | 2.92067 |
| 0.512624 | 2.90929 |
| 0.788593 | 2.90612 |
| 0.858019 | 2.8995 |
| 0.728314 | 2.89364 |
| 0.618869 | 2.88571 |
| -0.772093 | 2.88456 |
| 1.53672 | 2.86631 |
| 0.92644 | 2.85919 |
| 0.893925 | 2.83998 |
| -0.888371 | 2.83084 |
| 0.707912 | 2.82898 |
| -1.35309 | 2.82677 |
| 0.648239 | 2.8232 |
| 0.85501 | 2.82069 |
| 0.799396 | 2.81592 |
| -0.705327 | 2.80473 |
| 2.07344 | 2.80315 |
| 1.42803 | 2.79572 |
| 0.858377 | 2.79362 |
| 0.866509 | 2.7908 |
| -0.772811 | 2.78998 |
| 0.691699 | 2.78342 |
| -0.867235 | 2.77197 |
| 0.725378 | 2.76697 |
| -0.904497 | 2.75549 |
| 1.20902 | 2.74581 |
| -0.555263 | 2.74257 |
| 0.599395 | 2.74065 |
| 0.867834 | 2.73771 |
| 0.996173 | 2.73325 |
| 0.717234 | 2.72834 |
| 0.601657 | 2.72663 |
| 0.742667 | 2.72213 |
| 1.11263 | 2.72196 |
| 0.783715 | 2.71584 |
| -1.22112 | 2.71505 |
| 0.727287 | 2.71204 |
| 0.715517 | 2.711 |
| 0.89876 | 2.70993 |
| 1.45369 | 2.70945 |
| 0.726402 | 2.7066 |
| -0.662388 | 2.70639 |
| -0.702956 | 2.70319 |
| -1.35541 | 2.69448 |
| -0.575691 | 2.69277 |
| -0.484754 | 2.68765 |
| 0.733767 | 2.68486 |
| -1.34084 | 2.68008 |
| 0.529148 | 2.67667 |
| 0.705135 | 2.67612 |
| 0.587968 | 2.674 |
| 0.674301 | 2.66817 |
| 0.601838 | 2.65792 |
| 2.36778 | 2.65652 |
| -0.944864 | 2.65385 |
| 0.937179 | 2.64179 |
| -0.549666 | 2.64041 |
| -1.01895 | 2.63788 |
| 2.40398 | 2.63211 |
| -0.974523 | 2.6316 |
| 0.685123 | 2.62676 |
| -0.4515 | 2.62291 |
| -0.878414 | 2.61991 |
| -0.824434 | 2.61616 |
| 0.850653 | 2.61532 |
| -0.464939 | 2.61452 |
| 1.20327 | 2.61262 |
| -0.856835 | 2.61122 |
| -1.42885 | 2.60943 |
| 1.03211 | 2.60858 |
| 1.30984 | 2.605 |
| -0.650524 | 2.60367 |
| 0.885031 | 2.59923 |
| 0.608034 | 2.59342 |
| 0.686395 | 2.59325 |
| 0.451604 | 2.5884 |
| 1.10389 | 2.58352 |
| -0.496462 | 2.57855 |
| 0.619835 | 2.57759 |
| -0.533364 | 2.57305 |
| 0.582424 | 2.57247 |
| -0.536319 | 2.56402 |
| -1.18556 | 2.5596 |
| 1.11665 | 2.55678 |
| 0.660112 | 2.55576 |
| -0.830519 | 2.55218 |
| 0.620585 | 2.54486 |
| -0.621856 | 2.54325 |
| -0.553939 | 2.54226 |
| 1.19778 | 2.54059 |
| -0.563505 | 2.53585 |
| -1.01981 | 2.53458 |
| 0.948758 | 2.53386 |
| -0.554202 | 2.52809 |
| 0.681548 | 2.52267 |
| -1.21575 | 2.52082 |
| 0.704463 | 2.51946 |
| 1.0067 | 2.51854 |
| -0.639003 | 2.51693 |
| 0.592785 | 2.51574 |
| 0.76896 | 2.50713 |
| 0.709675 | 2.50277 |
| 0.740303 | 2.48939 |
| -0.710084 | 2.48814 |
| 1.2535 | 2.48771 |
| 0.550546 | 2.48477 |
| -1.02964 | 2.47824 |
| 0.672743 | 2.46784 |
| 0.764537 | 2.46776 |
| -0.528214 | 2.46658 |

| Log Fold-change | "Increased" |
| --- | --- |
| 0.372507 | 1.99778 |
| 0.373286 | 1.95096 |
| 0.37465 | 2.07867 |
| 0.37525 | 1.40501 |
| 0.375356 | 1.60567 |
| 0.37729 | 1.65941 |
| 0.379536 | 1.57737 |
| 0.381631 | 2.0767 |
| 0.384076 | 2.21068 |
| 0.38438 | 1.47976 |
| 0.384901 | 2.1439 |
| 0.384931 | 1.32378 |
| 0.384984 | 1.3599 |
| 0.387615 | 1.55063 |
| 0.38837 | 1.49194 |
| 0.388533 | 1.6028 |
| 0.389672 | 1.87654 |
| 0.391336 | 2.0576 |
| 0.391929 | 1.52986 |
| 0.397851 | 1.59711 |
| 0.398524 | 1.51703 |
| 0.398674 | 1.35484 |
| 0.400015 | 1.589 |
| 0.400808 | 1.32331 |
| 0.401466 | 1.44567 |
| 0.401619 | 1.65008 |
| 0.403326 | 2.19206 |
| 0.403459 | 1.86678 |
| 0.40379 | 1.98271 |
| 0.404582 | 1.32478 |
| 0.404868 | 1.38254 |
| 0.405398 | 1.61639 |
| 0.406435 | 1.60243 |
| 0.407284 | 1.3033 |
| 0.407888 | 1.44135 |
| 0.409845 | 2.15848 |
| 0.410166 | 1.80971 |
| 0.410299 | 1.72733 |
| 0.412535 | 2.01952 |
| 0.41637 | 1.84783 |
| 0.417426 | 1.33607 |
| 0.426007 | 1.43834 |
| 0.431762 | 1.84023 |
| 0.434125 | 1.93069 |
| 0.434625 | 1.33303 |
| 0.434681 | 1.31337 |
| 0.436118 | 1.62383 |
| 0.436125 | 2.10306 |
| 0.438826 | 2.27886 |
| 0.439241 | 1.8103 |
| 0.440878 | 1.31609 |
| 0.441651 | 2.39046 |
| 0.442156 | 1.97096 |
| 0.446987 | 2.05683 |
| 0.4489 | 1.62684 |
| 0.450813 | 1.71674 |
| 0.451604 | 2.5884 |
| 0.451661 | 1.55466 |
| 0.453042 | 2.20589 |
| 0.453749 | 1.69813 |
| 0.459957 | 1.33057 |
| 0.46236 | 1.30452 |
| 0.464475 | 1.54001 |
| 0.465963 | 2.11579 |
| 0.470261 | 1.40808 |
| 0.473139 | 1.32947 |
| 0.476613 | 1.57147 |
| 0.476793 | 1.33628 |
| 0.476916 | 1.78551 |
| 0.480962 | 1.9371 |
| 0.481253 | 1.63099 |
| 0.481882 | 1.4129 |
| 0.482603 | 1.94002 |
| 0.48381 | 1.78884 |
| 0.486532 | 1.3112 |
| 0.491453 | 1.85261 |
| 0.493471 | 1.67971 |
| 0.494509 | 1.80313 |
| 0.498458 | 1.96913 |
| 0.498838 | 2.04494 |
| 0.499652 | 1.33182 |
| 0.499951 | 1.31911 |
| 0.506348 | 1.97123 |
| 0.507967 | 1.87693 |
| 0.508699 | 1.75552 |
| 0.510678 | 1.5488 |
| 0.512624 | 2.90929 |
| 0.512724 | 2.40231 |
| 0.512876 | 1.80532 |
| 0.513914 | 1.7396 |
| 0.514356 | 2.30878 |
| 0.515925 | 1.48354 |
| 0.516823 | 2.17088 |
| 0.522947 | 1.79133 |
| 0.524652 | 1.9122 |
| 0.525275 | 2.05689 |
| 0.525603 | 1.63008 |
| 0.527314 | 1.38092 |
| 0.527529 | 3.12169 |
| 0.528151 | 1.89676 |
| 0.528661 | 1.45443 |
| 0.529148 | 2.67667 |
| 0.530498 | 1.763 |
| 0.531196 | 1.78729 |
| 0.53156 | 1.77622 |
| 0.533942 | 2.22559 |
| 0.533993 | 1.89049 |
| 0.537588 | 1.86447 |
| 0.539022 | 1.3531 |
| 0.539589 | 1.62215 |
| 0.539665 | 1.31969 |
| 0.540308 | 1.82831 |
| 0.543704 | 1.75094 |
| 0.543766 | 1.40978 |
| 0.544664 | 1.46549 |
| 0.544803 | 1.46919 |
| 0.548484 | 1.7395 |
| 0.550546 | 2.48477 |
| 0.550619 | 1.35387 |
| 0.555425 | 1.46232 |
| 0.556368 | 1.52047 |
| 0.556428 | 2.16368 |
| 0.559676 | 1.62529 |
| 0.566486 | 1.77528 |
| 0.566495 | 1.47263 |
| 0.567315 | 1.34495 |
| 0.569001 | 2.22548 |
| 0.570746 | 1.97764 |
| 0.571808 | 2.01648 |
| 0.573787 | 1.43212 |
| 0.574223 | 1.64286 |
| 0.57448 | 1.94541 |
| 0.57562 | 1.82222 |
| 0.576208 | 2.23241 |
| 0.57767 | 2.30274 |
| 0.578257 | 2.10589 |
| 0.578876 | 1.65941 |
| 0.581706 | 1.72248 |
| 0.582424 | 2.57247 |
| 0.583024 | 1.67308 |
| 0.584754 | 1.86866 |
| 0.585043 | 1.36382 |
| 0.58567 | 2.22488 |
| 0.587304 | 1.91214 |
| 0.587968 | 2.674 |
| 0.590207 | 2.24081 |
| 0.590357 | 2.08965 |
| 0.591502 | 1.47886 |
| 0.592705 | 1.55544 |
| 0.592785 | 2.51574 |
| 0.593859 | 2.01005 |
| 0.595174 | 2.00104 |
| 0.595898 | 2.44705 |
| 0.599395 | 2.74065 |
| 0.59998 | 1.55608 |
| 0.601657 | 1.32976 |
| 0.601657 | 2.72663 |
| 0.601838 | 2.65792 |
| 0.604187 | 2.17013 |
| 0.60529 | 1.51349 |
| 0.608034 | 2.59342 |
| 0.609044 | 2.09568 |
| 0.611253 | 2.21973 |
| 0.611536 | 1.4493 |
| 0.611564 | 1.37089 |
| 0.6117 | 2.36261 |
| 0.612224 | 1.66126 |
| 0.612418 | 2.08678 |
| 0.615386 | 1.93585 |
| 0.617256 | 3.1111 |
| 0.618553 | 1.89394 |
| 0.618869 | 2.88571 |
| 0.619835 | 2.57759 |
| 0.620585 | 2.54486 |
| 0.621523 | 2.21919 |
| 0.621607 | 1.69856 |
| 0.624885 | 1.32665 |
| 0.625359 | 2.16289 |
| 0.631849 | 1.80108 |
| 0.631967 | 1.5927 |
| 0.633701 | 1.90816 |
| 0.633907 | 1.90723 |
| 0.635023 | 1.3214 |
| 0.635107 | 1.75013 |
| 0.63605 | 1.32472 |
| 0.637508 | 1.51371 |
| 0.637826 | 3.02912 |
| 0.640248 | 1.72451 |
| 0.641525 | 2.00804 |
| 0.641733 | 1.40509 |
| 0.642742 | 2.24119 |
| 0.64423 | 1.5496 |
| 0.644857 | 1.82956 |
| 0.644877 | 1.76258 |
| 0.645391 | 2.26195 |
| 0.648239 | 2.8232 |
| 0.648519 | 1.55648 |
| 0.649434 | 1.81036 |
| 0.653814 | 2.96028 |
| 0.654001 | 1.45665 |

| Log Fold-change | "Decreased" |
| --- | --- |
| -3.95445 | 2.42098 |
| -3.86088 | 5.39384 |
| -2.4715 | 1.87977 |
| -2.30968 | 4.29741 |
| -1.96505 | 4.13595 |
| -1.77089 | 1.8197 |
| -1.74927 | 2.28391 |
| -1.6909 | 2.35466 |
| -1.67679 | 2.44675 |
| -1.67417 | 1.73208 |
| -1.67244 | 2.99531 |
| -1.62159 | 2.14138 |
| -1.61535 | 1.40868 |
| -1.5808 | 2.346 |
| -1.55558 | 2.43036 |
| -1.5069 | 2.19302 |
| -1.49894 | 1.36729 |
| -1.46489 | 1.57454 |
| -1.45258 | 1.9503 |
| -1.44562 | 4.14434 |
| -1.44243 | 2.46415 |
| -1.42885 | 2.60943 |
| -1.42027 | 2.21902 |
| -1.3781 | 2.3949 |
| -1.36737 | 2.95722 |
| -1.36394 | 2.92368 |
| -1.35677 | 2.4228 |
| -1.35541 | 2.69448 |
| -1.35309 | 2.82677 |
| -1.35301 | 3.55696 |
| -1.34084 | 2.68008 |
| -1.33205 | 3.05255 |
| -1.3158 | 3.30177 |
| -1.31313 | 3.33764 |
| -1.29436 | 1.53111 |
| -1.28392 | 2.92582 |
| -1.27986 | 2.06768 |
| -1.2713 | 3.39225 |
| -1.24353 | 2.38421 |
| -1.2426 | 2.22809 |
| -1.22112 | 2.71505 |
| -1.21575 | 2.52082 |
| -1.20297 | 3.86603 |
| -1.1982 | 1.81949 |
| -1.18556 | 2.5596 |
| -1.18045 | 3.62838 |
| -1.1699 | 1.67185 |
| -1.13887 | 1.4141 |
| -1.12955 | 2.05442 |
| -1.1048 | 1.56192 |
| -1.08223 | 1.90708 |
| -1.07578 | 2.45717 |
| -1.0637 | 1.83392 |
| -1.0508 | 1.8822 |
| -1.04961 | 2.05014 |
| -1.04778 | 1.67514 |
| -1.03945 | 4.87159 |
| -1.02964 | 2.47824 |
| -1.02505 | 1.82539 |
| -1.02289 | 2.32828 |
| -1.01981 | 2.53458 |
| -1.01973 | 1.33455 |
| -1.01895 | 2.63788 |
| -1.01201 | 2.21558 |
| -1.01174 | 1.41468 |
| -1.00236 | 3.15811 |
| -0.998735 | 3.46671 |
| -0.992493 | 3.61875 |
| -0.986351 | 1.89493 |
| -0.978038 | 1.62556 |
| -0.974523 | 2.6316 |
| -0.971859 | 1.45265 |
| -0.961188 | 1.56347 |
| -0.956877 | 1.55675 |
| -0.949893 | 1.45438 |
| -0.949778 | 2.32372 |
| -0.948729 | 2.04482 |
| -0.944864 | 2.65385 |
| -0.941805 | 1.5903 |
| -0.929269 | 1.90896 |
| -0.925309 | 1.32245 |
| -0.925096 | 1.61871 |
| -0.924389 | 3.02281 |
| -0.920209 | 3.73558 |
| -0.904497 | 2.75549 |
| -0.902573 | 1.74702 |
| -0.889955 | 3.97907 |
| -0.888371 | 2.83084 |
| -0.887608 | 1.56033 |
| -0.880477 | 2.36522 |
| -0.878642 | 1.8068 |
| -0.878414 | 2.61991 |
| -0.868128 | 2.25586 |
| -0.867235 | 2.77197 |
| -0.862988 | 2.26305 |
| -0.861603 | 1.84429 |
| -0.85816 | 2.23142 |
| -0.856835 | 2.61122 |
| -0.856731 | 1.63989 |
| -0.851273 | 1.98817 |
| -0.85018 | 1.82333 |
| -0.850044 | 1.40462 |
| -0.841747 | 2.24653 |
| -0.837479 | 3.1193 |
| -0.836554 | 1.34586 |
| -0.830519 | 2.55218 |
| -0.824434 | 2.61616 |
| -0.822585 | 1.65956 |
| -0.80957 | 1.80097 |
| -0.808319 | 1.58814 |
| -0.807984 | 1.32589 |
| -0.807636 | 2.28516 |
| -0.806593 | 1.36626 |
| -0.800763 | 1.9698 |
| -0.78966 | 1.55929 |
| -0.780619 | 2.1129 |
| -0.777457 | 1.5695 |
| -0.775823 | 1.78803 |
| -0.774617 | 2.2237 |
| -0.772811 | 2.78998 |
| -0.772093 | 2.88456 |
| -0.770887 | 1.69382 |
| -0.769934 | 1.56688 |
| -0.767913 | 2.28084 |
| -0.765859 | 1.51052 |
| -0.764698 | 1.89615 |
| -0.763788 | 2.37782 |
| -0.758619 | 3.10776 |
| -0.758443 | 1.69235 |
| -0.75699 | 1.78875 |
| -0.754534 | 1.39654 |
| -0.751929 | 2.92067 |
| -0.751377 | 2.93889 |
| -0.749584 | 1.81158 |
| -0.749579 | 1.69754 |
| -0.743786 | 2.30176 |
| -0.741869 | 3.24991 |
| -0.740151 | 1.54659 |
| -0.73997 | 2.36777 |
| -0.739061 | 1.68249 |
| -0.735313 | 2.38454 |
| -0.733936 | 1.93074 |
| -0.723705 | 1.43768 |
| -0.72333 | 1.72106 |
| -0.717484 | 1.4796 |
| -0.712654 | 1.44988 |
| -0.710329 | 2.08686 |
| -0.710084 | 2.48814 |
| -0.705327 | 2.80473 |
| -0.704731 | 1.58121 |
| -0.702956 | 2.70319 |
| -0.700652 | 1.94251 |
| -0.698089 | 2.10556 |
| -0.697624 | 1.52417 |
| -0.695548 | 3.33323 |
| -0.688445 | 1.61459 |
| -0.681283 | 1.40738 |
| -0.678377 | 2.2003 |
| -0.676665 | 1.30834 |
| -0.670772 | 2.06414 |
| -0.670188 | 1.52032 |
| -0.662388 | 2.70639 |
| -0.660299 | 2.13389 |
| -0.658444 | 1.3143 |
| -0.656374 | 2.05475 |
| -0.655434 | 1.37719 |
| -0.654627 | 1.46923 |
| -0.650524 | 2.60367 |
| -0.645968 | 1.90902 |
| -0.644478 | 1.93262 |
| -0.642693 | 2.11514 |
| -0.64051 | 2.98696 |
| -0.639003 | 2.51693 |
| -0.635332 | 2.38578 |
| -0.633478 | 2.44184 |
| -0.629496 | 2.23007 |
| -0.623965 | 1.90109 |
| -0.622686 | 2.10475 |
| -0.621856 | 2.54325 |
| -0.621676 | 3.41001 |
| -0.620819 | 2.37073 |
| -0.619289 | 1.63949 |
| -0.614891 | 2.00452 |
| -0.610301 | 1.41859 |
| -0.609658 | 1.80189 |
| -0.607577 | 2.10185 |
| -0.606186 | 1.73982 |
| -0.605795 | 2.20066 |
| -0.603653 | 1.42404 |
| -0.602476 | 1.80715 |
| -0.601488 | 1.72074 |
| -0.600537 | 1.80873 |
| -0.597004 | 1.97917 |
| -0.597003 | 1.65748 |
| -0.596336 | 2.46082 |
| -0.593284 | 1.31889 |
| -0.590388 | 2.20714 |
| -0.589325 | 2.05779 |
| -0.589197 | 1.39439 |
| -0.588423 | 1.30801 |

**Figure 10**. Volcano plot illustrates significantly differentially abundant proteins in intestine. The −log_10_ (*P value*) is plotted against the log_2_ (fold change: diet B/diet A).

**Table 7.** Top ten DEPs in the Intestine tissue

| **Row** | **Protein ID** | **Protein Name** | **Fold change*** |
| --- | --- | --- | --- |
| **Increased abundance in diet B** | | | |
| 1 | A0A4W6DGN9 | Threonyl carbamoyl-AMP synthase | 9.12 |
| 2 | A0A4W6CQR5 | GIT ArfGAP 1 | 6.92 |
| 3 | A0A4W6DG98 | NACHT domain-containing protein | 5.29 |
| 4 | A0A4W6EI74 | Apolipoprotein Ea | 5.16 |
| 5 | A0A4W6FS56 | RUN and SH3 domain containing 1 | 4.74 |
| 6 | A0A4W6DP25 | YTH domain-containing family protein | 4.40 |
| 7 | A0A4W6D1L2 | Metallo endopeptidase | 4.20 |
| 8 | A0A4W6DT98 | DnaJ (Hsp40) homolog, subfamily C, member 8 | 4.03 |
| 9 | A0A4W6FGH5 | Bridging integrator 2b | 4.00 |
| 10 | A0A4W6CAB3 | AP complex subunit sigma | 3.92 |
| **Decreased abundance in diet B** | | | |
| 11 | A0A4W6DJG2 | C1q domain-containing protein | 0.06 |
| 12 | A0A4W6EI75 | non-specific serine/threonine protein kinase | 0.06 |
| 13 | A0A4W6BTQ3 | UPAR/Ly6 domain-containing protein | 0.18 |
| 14 | A0A4W6FXK4 | phosphatidylinositol-4,5-bisphosphate 3-kinase | 0.20 |
| 15 | A0A4W6G0X6 | H1.0 linker histone | 0.25 |
| 16 | A0A4W6CTG1 | Uncharacterized protein | 0.29 |
| 17 | A0A4W6F148 | Zmp:0000000846 | 0.29 |
| 18 | A0A4W6BT82 | Tyrosine-protein kinase (EC 2.7.10.2) | 0.30 |
| 19 | A0A4W6BTD9 | Solute carrier family 25-member 18 | 0.31 |
| 20 | A0A4W6ECB4 | Fatty acid-binding protein | 0.31 |

*Values > 1.3 indicate more abundant in diet B, values < 0.76 indicate less abundant in diet B

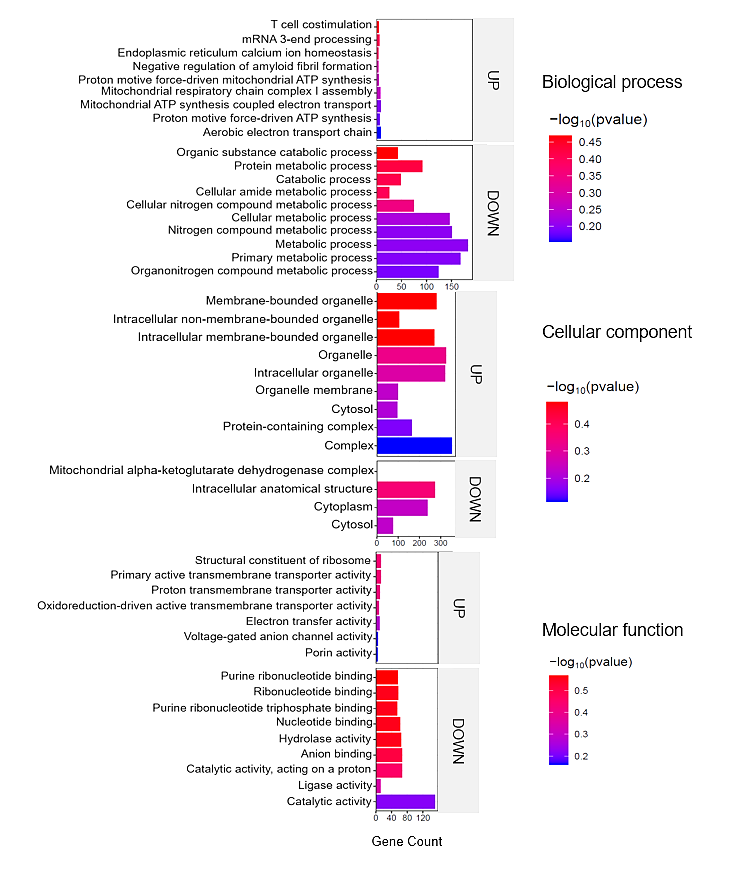

**Figure 11.** Top enriched GO terms in the DEPS of the intestine of barramundi. A higher –log₁₀(p-value) (red) indicates greater statistical significance, while lower –log₁₀(p-value) (blue) values indicate less significant enrichment. Up indicates increased abundance in the intestine of fish fed on diet B, while Down indicates decreased abundance in the intestine of fish fed on diet B.

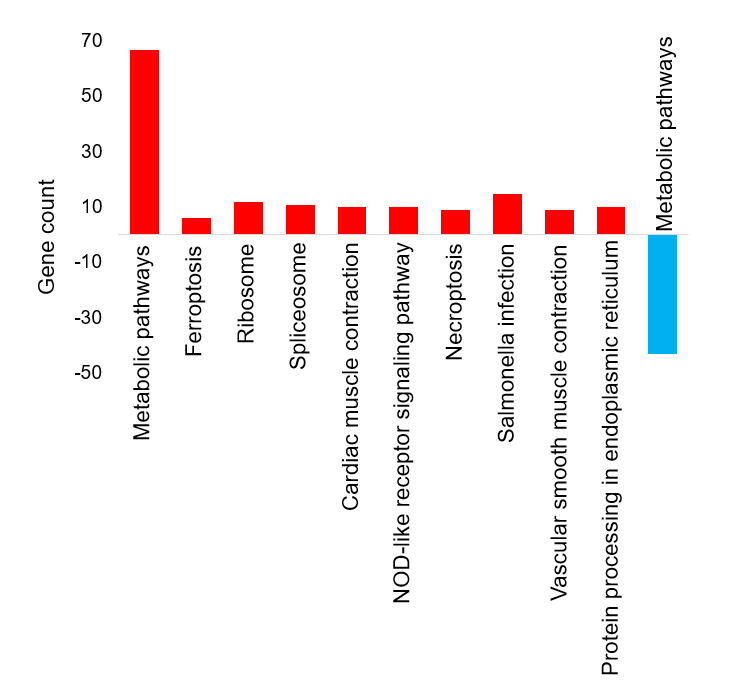

**Figure 12.** Top KEGG pathways of the differentially abundant proteins in Intestine tissue

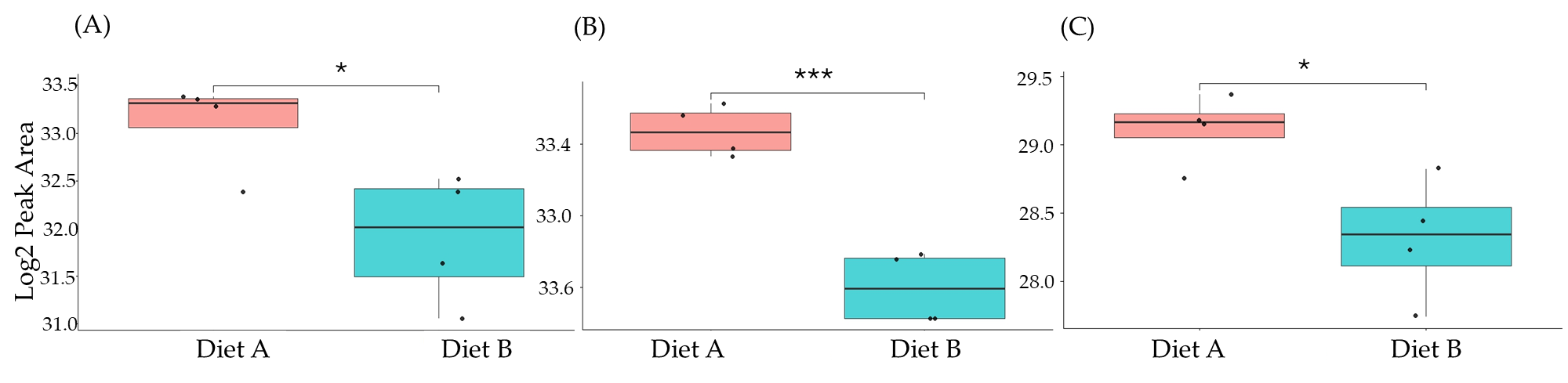
**Figure 13.** PRM validation of three proteins changed in abundance in response to dietary composition in brain tissue in Barramundi. (A) = Guanine nucleotide binding protein (G protein), (B) = ATP synthase subunit beta (EC 7.1.2.2), (C) = ADP ribosylation factor like GTPase 3b. An asterisk (*, **, ***) indicates a statistically significant difference between diet A and diet B, according to a Student t-test (p-value ≤ 0.05, ≤ 0.01, ≤ 0.001).

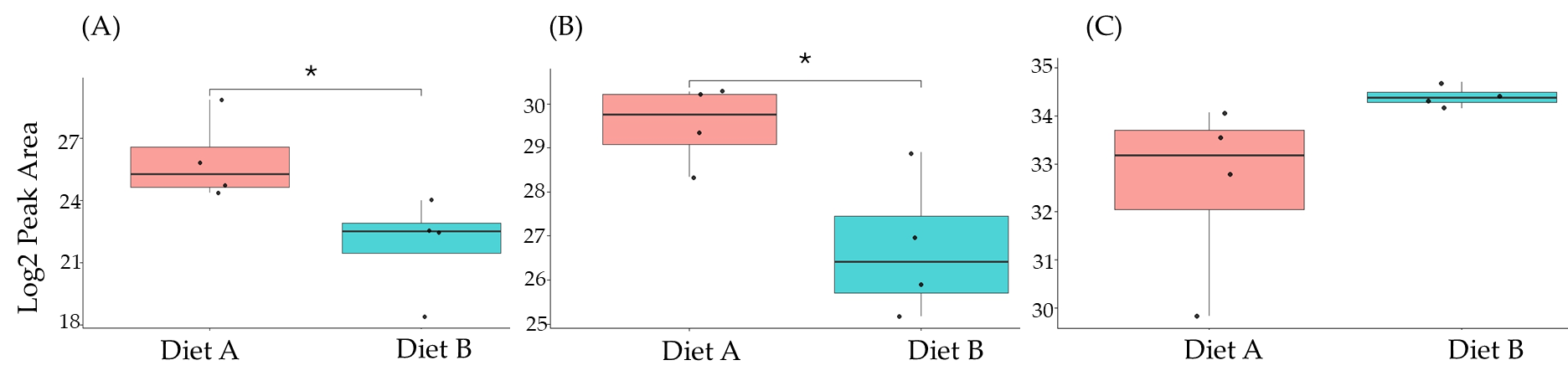

**Figure 14.** PRM validation of two proteins changed in abundance in response to dietary composition in liver tissue in Barramundi. (A) = Lysozyme g, (B) = C1q domain-containing protein, (C) = pancreatic elastase. An asterisk (*, **, ***) indicates a statistically significant difference between diet A and diet B, according to a Student t-test (p-value ≤0.05, ≤ 0.01, ≤ 0.001)

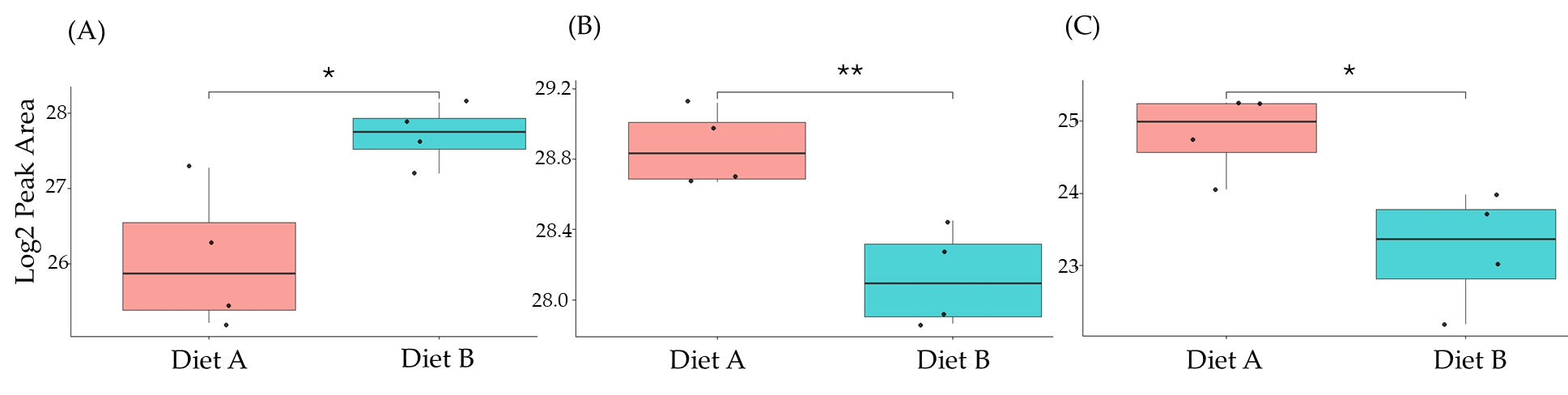

**Figure 15.** PRM validation of three proteins changed in abundance in response to dietary composition in intestine tissue in Barramundi. (A) = Apolipoprotein Ea, (B) = non-specific serine/threonine protein kinase, (C) = phosphatidylinositol-4,5-bisphosphate 3-kinase. An asterisk (*, **, ***) indicates a statistically significant difference between diet A and diet B, according to a Student t-test (p-value ≤ 0.05, ≤ 0.01, ≤ 0.001).

|  | Iron Concentration (µg/g) |
| --- | --- |
| Diet A | 0.514984 |
| Diet B | 0.605038 |

**Figure 16.** Total iron concentration (µg/g) brain tissue of the barramundi fed on diet A and B. Error bars represent the standard deviation (n= 4).

|  | Iron Concentration (µg/g) |
| --- | --- |
| Diet A | 64.9667 |
| Diet B | 123.867 |

**Figure 17.** Total iron concentration (µg/g) liver tissue of the barramundi fed on diet A and B. Bars represent the standard deviation (n= 4).

|  | Iron Concentration (µg/g) |
| --- | --- |
| Diet A | 0.425445 |
| Diet B | 0.803549 |

**Figure 18.** Total iron concentration (µg/g) in intestine tissue of the barramundi fed on diet A and B. Bars represent the standard deviation (n= 4). Asterisk (*) indicates a statistically significant difference between groups as determined by Student’s t-test (p ≤ 0.05).
